# BugSeq: a highly accurate cloud platform for long-read metagenomic analyses

**DOI:** 10.1101/2020.10.08.329920

**Authors:** Jeremy Fan, Steven Huang, Samuel D Chorlton

## Abstract

**Background:** As the use of nanopore sequencing for metagenomic analysis increases, tools capable of performing long-read taxonomic classification in a fast and accurate manner are needed. Existing tools were either designed for short-read data (eg. Centrifuge) or take days to analyse modern sequencer outputs (eg. MetaMaps).

**Results:** We present BugSeq, a novel, highly accurate metagenomic classifier for nanopore reads. BugSeq (F1=0.91-0.95) offers better read classification than MetaMaps (F1=0.89-0.94) in a fraction of the time. BugSeq significantly improves on the accuracy of Centrifuge (F1=0.79-0.93) while offering competitive run times. We apply BugSeq to metagenomic sequencing of 41 samples from patients with lower respiratory tract infections and show that it produces greater concordance with microbiological culture and qPCR compared with “What’s In My Pot” analysis.

**Conclusion:** BugSeq is deployed to the cloud for easy and scalable long-read metagenomic analyses. BugSeq is freely available for non-commercial use at https://bugseq.com/free.

## Background

Nanopore sequencing has seen a dramatic increase in read quality and throughput over the last few years, leading to increased adoption and novel applications. Recently, nanopore sequencing has been used for metagenomics in clinical, environmental and agricultural settings (Petersen *et al*., 2019; Edwards *et al*., 2019; Stewart *et al*., 2019). Many metagenomic read classifiers, originally designed for short (<300bp), high quality reads rely on k-mers for sequence classification (Breitwieser *et al*., 2017). Due to the high error rate of nanopore sequencing and the low likelihood of many consecutive error-free bases, k-mer methods are unlikely to be optimal for nanopore read classification. Alternative methods have been explored: EPI2ME, a platform operated by Metrichor, uses Centrifuge as its classifier (Kim *et al*., 2016). Centrifuge can start with short k-mer matches and extend them until the first nucleotide difference in alignment, enabling variable length matches. By default, Centrifuge starts this extension with 22bp seeds, however this parameter can be set down to 16bp for increased sensitivity. MetaMaps relies on approximate read alignment with minimizers and a probabilistic model to estimate sample composition (Dilthey *et al*., 2019). Finally, deSAMBA performs pseudo-alignment against De Bruijn graphs and assigns reads to the top scoring hit (Li *et al*., 2019).

## Implementation

We combined a fast and accurate read mapper, Bayesian reassignment of reads based on mapping quality, a new lowest-common ancestor process, and an advanced visualization tool to build a better metagenomic classifier for nanopore reads. This pipeline, which we call BugSeq, has been packaged with Nextflow and made available as an online service (https://bugseq.com/free) for easy cloud analyses. In brief, reads are quality controlled with fastp, mapped with minimap2 to an index, and reassigned based on a Bayesian statistical framework using Pathoscope (Chen *et al*., 2018; Li, 2018; Francis *et al*., 2013). Finally, the lowest common ancestor of reassigned reads is calculated and inputted into Recentrifuge for summarization and visualization (Martí, 2019). Quality control results are summarized with MultiQC (Ewels *et al*, 2016). All dependencies are packaged in Docker images, and jobs are executed on Amazon Web Services Batch in a secure, private environment.

## Results

We assessed the performance of BugSeq and compared it with three competing tools: Centrifuge, MetaMaps and deSAMBA (Kim *et al*., 2016; Dilthey *et al*., 2019; Li *et al*., 2019). We used nanopore sequencing data from two microbial communities with known composition. The ZymoBIOMICS mock communities contain 8 bacteria and 2 yeast in even (hereafter referred to as “Even”) and logarithmic (hereafter referred to as “Log”) concentrations (Nicholls *et al*., 2019). Recall, precision and F-scores were determined for the three classifiers at the read-level; reads classified to any of the expected taxonomic nodes were considered correct at that taxonomic rank and any rank above, otherwise the read was considered incorrect. Full results at each taxonomic rank, along with definitions for each performance metric, can be found in Supplementary Table 1. All tools were evaluated using 32 threads, 280Gb of RAM (our server capacity) and default tool parameters unless otherwise specified. All evaluations used a reference database from RefSeq (see supplementary information). deSAMBA failed to index the RefSeq database with 280Gb of RAM, excluding it from further analysis.

At the species level, BugSeq attained the top precision and recall compared with MetaMaps and Centrifuge across both Even and Log datasets (F1-score_Even_: 0.91, F1-score_Log_: 0.95) (Table 1). On average, BugSeq had 2% better recall than MetaMaps while maintaining superior precision, and 2-5% better precision than Centrifuge while maintaining superior recall. When analyzing the number of unique species identified by each tool (true count=10), BugSeq found a total of 68 and 31 species in the Zymo Even and Log dataset, respectively. In comparison, MetaMaps identified 2144 (Log) and 1386 (Even) unique species using the “miniSeq+H” database and exceeded our RAM threshold with the RefSeq database. Centrifuge identified 5380 and 5513 species in the Even and Log dataset, respectively.

**Table 1.** Performance of four metagenomic classifiers (BugSeq, MetaMaps, Centrifuge, and deSamba) on GridION ZymoBIOMICS Mock Log and Even communities. Different k-mer seed sizes were explored for Centrifuge (16 bp and 22 bp), and two different databases were examined for MetaMaps (miniSeq+H and RefSeq); otherwise, default parameters were used. The best result for each column and dataset combination is bolded.

|  | Dataset Used | F1 score at species level | Precision at species level (%) | Recall at species level (%) | Number of unique species detected (true count=10) | Run time (wall clock) [hh:mm] | Classification memory requirement [GB] | Indexing time (wall clock) [hh:mm] | Indexing memory requirement [GB] |
| --- | --- | --- | --- | --- | --- | --- | --- | --- | --- |
| BugSeq | Even | **0.91** | **99.57** | **83.69** | **68** | 4:09 | 118 | 1:15 | 243 |
|  | Log | **0.95** | **99.92** | **90.47** | **31** | 4:25 | 113 |  |  |
| MetaMaps (miniSeq+H) | Even | 0.89 | 99.13 | 80.71 | 2144 | 86:20 | 172.6 | **Prebuilt** | **Prebuilt** |
|  | Log | 0.94 | 99.46 | 88.25 | 1386 | 125:22 | 184.0 |  |  |
| MetaMaps (RefSeq) | Even | Out of memory |  |  |  |  |  | NA | NA |
|  | Log | 0.94 | 99.73 | 88.66 | 2094 | 154:29 | 270 |  |  |
| Centrifuge – 16bp minimum hit length, -k 1 | Even | 0.79 | 93.00 | 69.17 | 4337 | **00:19** | **67.7** | 19:45 | 168 |
|  | Log | 0.92 | 95.93 | 89.07 | 5777 | **00:19** | **64.4** |  |  |
| Centrifuge – default (22bp) minimum hit length, -k 1 | Even | 0.80 | 94.71 | 68.53 | 5380 | **00:16** | **56.9** |  |  |
|  | Log | 0.93 | 97.14 | 88.65 | 5513 | **00:14** | **55.7** |  |  |
| DeSAMBA | Even | Out of memory |  |  |  |  |  |  | >280 GB |
|  | Log | Out of memory |  |  |  |  |  |  | >280 GB |

We next measured computational performance for all tools. BugSeq is an order of magnitude faster than MetaMaps, which took over 5 days using 32 cores and their “miniSeq+H” database. BugSeq took up to 4 hours and 25 minutes to analyse the same amount of data. Notably, MetaMap’s miniSeq+H (26 GB) is significantly smaller than Refeq (86 GB). BugSeq offered marginally longer run times than Centrifuge, which took 14 to 19 minutes for all analyses. All tools required more than 50GB of RAM for execution, precluding their use on modern laptops.

To evaluate BugSeq on real clinical samples, we applied it to nanopore metagenomic sequencing of 41 lower respiratory tracts samples from patients with bacterial lower respiratory infections. Sample characteristics and data generation have been previously reported (Charalampous *et al*, 2019). We used the original authors’ 1% abundance threshold to report pathogenic microbes, ensuring comparability across methods. The results of quality control and metagenomic classification are visualized in the supplementary HTML files. BugSeq reached better concordance with traditional culture results, as compared with the original “What’s In My Pot” (WIMP) analysis, in 3/41 (7.3%) samples (Supplementary Table 2). Specifically, samples S8, S15 and S21 each had *S. pneumoniae* detected by WIMP analysis but not by BugSeq or microbial culture. Pathogen-specific qPCR on these samples failed to detect *S. pneumoniae*, confirming these findings (Charalampous *et al*, 2019). Additionally, BugSeq reached better concordance with qPCR, but not microbial culture, in 1/41 samples (sample S12), where WIMP detected a false-positive *H. influenzae* not detected by BugSeq or qPCR. Sample S32 was the only other discordant sample between BugSeq and WIMP: BugSeq detected an *E. coli* and *S. flexneri*, WIMP detected an *E. coli*, and the cultures were reported as “no significant growth”. As there was no qPCR for *S. flexneri* performed on this sample, the implications of this discordance are unclear.

## Discussion

Here we present BugSeq, an accurate and fast metagenomic classifier for nanopore reads. On mock microbial communities, BugSeq was found to outperform both MetaMaps and Centrifuge, sometimes by large margins, in terms of precision and recall. BugSeq was also faster than MetaMaps by an order of days, while offering minimal time trade-off with Centrifuge. BugSeq achieves better classification performance with its reliance on underlying performant tools. Minimap2, which performs BugSeq’s alignment step, is over 30 times faster than most long-read aligners and demonstrated the highest alignment accuracy at the time of its publication (Li, 2018, 2). Preprocessing relies on fastp, which again is optimized for speed by relying on C++ under the hood (Chen *et al*, 2018).

The results of our study are concordant with existing literature on long-read metagenomic classifiers (Li *et al*., 2019). We found a lower sensitivity for Centrifuge, which could be attributed to cases in which Centrifuge returns multiple assignments for a single read and collapses these up the taxonomic tree via lowest common ancestor. Similarly, the original MetaMaps paper identified a RAM use of 262 GB and 209 CPU hours for a random sample of a third of the Zymo dataset (Dilthey *et al*, 2019). In our experience, MetaMaps mapped reads relatively quickly, in accordance with published data on its MinHash-based aligner, but stalled on its “classification” step (Jain *et al*, 2018).

In addition to superior performance, BugSeq is deployed to the cloud to enable automatic metagenomic analysis from raw reads to report. BugSeq’s user interface only requires a simple upload of FASTQ files to its website, and returns to the user intuitive HTML files for visualization in their browser. We demonstrate the ease of use of BugSeq by uploading metagenomic data from 41 lower respiratory tract samples. Resulting data, including quality control and metagenomic classification, was packaged into two HTML files (Supplementary Material), and showed superior accuracy compared with the original WIMP analysis on the same data.

## Conclusion

BugSeq is a rapid, scalable and accurate metagenomics classifier that outperforms alternatives such as MetaMaps and Centrifuge across a range of performance indicators. BugSeq is deployed to the cloud for easy metagenomic analyses.

## Supporting information

Quality Control Information

Metagenomic Classification

## Availability and requirements

Project name: BugSeq

Project home page: https://bugseq.com/free

Operating system(s): Platform independent

Programming language: Nextflow

Other requirements: Modern internet browser such as Firefox, Chrome, Safari or Edge

Any restrictions to use by non-academics: Licence required

## List of abbreviations

HTML: Hypertext Markup Language
qPCR: Quantitative Polymerase Chain Reaction
WIMP: What’s In My Pot

