## Supplementary material for "BugSeq: a highly accurate cloud platform for long-read metagenomic analyses": Metagenomic Classification

 Javascript must be enabled to view this page.

countunassignedtidrankscoreS1.fastqS2.fastqS3.fastqS4.fastqS5.fastqS6.fastqS7.fastqS8.fastqS9.fastqS10.fastqS11.fastqS12.fastqS13.fastqS14.fastqS15.fastqS16.fastqS17.fastqS18.fastqS19.fastqS20.fastqS21.fastqS22.fastqS23.fastqS24.fastqS25.fastqS26.fastqS27.fastqS28.fastqS29.fastqS30.fastqS31.fastqS32.fastqS33.fastqS34.fastqS35.fastqS36.fastqS37.fastqS38.fastqS39.fastqS40.fastqS41.fastqS2.fastq\_EXCLUSIVE\_speciesS3.fastq\_EXCLUSIVE\_speciesS4.fastq\_EXCLUSIVE\_speciesS5.fastq\_EXCLUSIVE\_speciesS7.fastq\_EXCLUSIVE\_speciesS8.fastq\_EXCLUSIVE\_speciesS9.fastq\_EXCLUSIVE\_speciesS11.fastq\_EXCLUSIVE\_speciesS12.fastq\_EXCLUSIVE\_speciesS13.fastq\_EXCLUSIVE\_speciesS14.fastq\_EXCLUSIVE\_speciesS15.fastq\_EXCLUSIVE\_speciesS16.fastq\_EXCLUSIVE\_speciesS17.fastq\_EXCLUSIVE\_speciesS19.fastq\_EXCLUSIVE\_speciesS20.fastq\_EXCLUSIVE\_speciesS21.fastq\_EXCLUSIVE\_speciesS22.fastq\_EXCLUSIVE\_speciesS23.fastq\_EXCLUSIVE\_speciesS24.fastq\_EXCLUSIVE\_speciesS26.fastq\_EXCLUSIVE\_speciesS27.fastq\_EXCLUSIVE\_speciesS28.fastq\_EXCLUSIVE\_speciesS29.fastq\_EXCLUSIVE\_speciesS30.fastq\_EXCLUSIVE\_speciesS31.fastq\_EXCLUSIVE\_speciesS32.fastq\_EXCLUSIVE\_speciesS33.fastq\_EXCLUSIVE\_speciesS34.fastq\_EXCLUSIVE\_speciesS36.fastq\_EXCLUSIVE\_speciesS38.fastq\_EXCLUSIVE\_speciesS39.fastq\_EXCLUSIVE\_speciesS40.fastq\_EXCLUSIVE\_speciesS41.fastq\_EXCLUSIVE\_speciesS3.fastq\_EXCLUSIVE\_genusS4.fastq\_EXCLUSIVE\_genusS5.fastq\_EXCLUSIVE\_genusS9.fastq\_EXCLUSIVE\_genusS11.fastq\_EXCLUSIVE\_genusS12.fastq\_EXCLUSIVE\_genusS13.fastq\_EXCLUSIVE\_genusS14.fastq\_EXCLUSIVE\_genusS15.fastq\_EXCLUSIVE\_genusS16.fastq\_EXCLUSIVE\_genusS19.fastq\_EXCLUSIVE\_genusS22.fastq\_EXCLUSIVE\_genusS23.fastq\_EXCLUSIVE\_genusS26.fastq\_EXCLUSIVE\_genusS28.fastq\_EXCLUSIVE\_genusS31.fastq\_EXCLUSIVE\_genusS34.fastq\_EXCLUSIVE\_genusS36.fastq\_EXCLUSIVE\_genusS38.fastq\_EXCLUSIVE\_genusS39.fastq\_EXCLUSIVE\_genusSHARED\_genusS3.fastq\_EXCLUSIVE\_familyS4.fastq\_EXCLUSIVE\_familyS14.fastq\_EXCLUSIVE\_familyS16.fastq\_EXCLUSIVE\_familyS26.fastq\_EXCLUSIVE\_familyS28.fastq\_EXCLUSIVE\_familyS31.fastq\_EXCLUSIVE\_familyS34.fastq\_EXCLUSIVE\_familySHARED\_familyS3.fastq\_EXCLUSIVE\_orderS4.fastq\_EXCLUSIVE\_orderS26.fastq\_EXCLUSIVE\_orderS28.fastq\_EXCLUSIVE\_orderS31.fastq\_EXCLUSIVE\_orderS34.fastq\_EXCLUSIVE\_orderSHARED\_orderS28.fastq\_EXCLUSIVE\_classS31.fastq\_EXCLUSIVE\_classS34.fastq\_EXCLUSIVE\_classSHARED\_classS26.fastq\_EXCLUSIVE\_phylumS31.fastq\_EXCLUSIVE\_phylumS34.fastq\_EXCLUSIVE\_phylumSHARED\_phylumS2.fastq\_EXCLUSIVE\_SUMMARYS3.fastq\_EXCLUSIVE\_SUMMARYS4.fastq\_EXCLUSIVE\_SUMMARYS5.fastq\_EXCLUSIVE\_SUMMARYS7.fastq\_EXCLUSIVE\_SUMMARYS8.fastq\_EXCLUSIVE\_SUMMARYS9.fastq\_EXCLUSIVE\_SUMMARYS11.fastq\_EXCLUSIVE\_SUMMARYS12.fastq\_EXCLUSIVE\_SUMMARYS13.fastq\_EXCLUSIVE\_SUMMARYS14.fastq\_EXCLUSIVE\_SUMMARYS15.fastq\_EXCLUSIVE\_SUMMARYS16.fastq\_EXCLUSIVE\_SUMMARYS17.fastq\_EXCLUSIVE\_SUMMARYS19.fastq\_EXCLUSIVE\_SUMMARYS20.fastq\_EXCLUSIVE\_SUMMARYS21.fastq\_EXCLUSIVE\_SUMMARYS22.fastq\_EXCLUSIVE\_SUMMARYS23.fastq\_EXCLUSIVE\_SUMMARYS24.fastq\_EXCLUSIVE\_SUMMARYS26.fastq\_EXCLUSIVE\_SUMMARYS27.fastq\_EXCLUSIVE\_SUMMARYS28.fastq\_EXCLUSIVE\_SUMMARYS29.fastq\_EXCLUSIVE\_SUMMARYS30.fastq\_EXCLUSIVE\_SUMMARYS31.fastq\_EXCLUSIVE\_SUMMARYS32.fastq\_EXCLUSIVE\_SUMMARYS33.fastq\_EXCLUSIVE\_SUMMARYS34.fastq\_EXCLUSIVE\_SUMMARYS36.fastq\_EXCLUSIVE\_SUMMARYS38.fastq\_EXCLUSIVE\_SUMMARYS39.fastq\_EXCLUSIVE\_SUMMARYS40.fastq\_EXCLUSIVE\_SUMMARYS41.fastq\_EXCLUSIVE\_SUMMARYSHARED\_SUMMARY 1083021740025447666019782132711871638179266995018736487367822716837463324228416420442384693655142769413213171230101538463575234302766572969992682716510680477891299514046296893181359986128635896572352078351711091326321103972471243322121390418705322756232084251942671632143016285131380671171616281101116141263051911440875706410921957912870821595486288159477193044753810318173519510913264211039724712434381215905187053227602320842599428816321463162951313318171196252561274125611118132613168448111331156181no\_rank3665.73361.12706.83932.23120.33829.43001.23062.53458.93586.12796.13196.43537.73351.03627.13656.63476.73336.43232.32991.23388.73138.23243.23591.63654.33440.52873.52544.74409.84187.62897.12467.63429.74010.83458.24666.53129.74795.03221.63510.73138.63037.42424.12582.63211.4739.2319.23293.7662.51924.61684.6707.94630.52921.72141.7495.22046.41181.81347.31495.6486.22677.13048.6700.91799.84439.52285.12185.43099.71119.91891.24170.03050.01501.94234.12750.91711.42787.81704.2561.62053.81483.9224.7261.2664.9857.82114.01578.11170.81024.71887.8878.6416.66074.33050.03243.6400.0592.0231.0571.11170.81429.62254.0112.93237.3343.3592.01170.8626.52349.5112.93189.6626.52615.0135.23439.91170.81782.5121.73416.83037.42342.02582.63211.4739.2319.23293.7662.51924.61675.3505.74623.02919.12141.7495.22046.41181.81347.31397.6486.22677.13048.6709.21799.84439.52210.52185.43099.7592.41891.24230.23050.01501.94234.13416.8108301173992518966581978213271187113815126694501683643736775271643743432408841432041738468365164276941293316903009353840357513429976596296389255271611067247788129461396429684318045998312802589517205201235221091326321103972381243322121389618645322756232083294267163214221622131386711716162811011166663051511424870641092157911670821595486168159477113044753863181735331091326421103972381243438121589618645322760232084094288163214431622131331817142113318142953462091511101914618222852836329112superkingdom3665.73361.22708.03932.93120.33829.43001.63064.03459.53586.82798.73196.73538.03351.63628.13657.03479.03336.53233.62991.23389.63139.33243.93591.93654.43440.62873.62546.54414.24187.72898.72467.73441.14029.13458.74667.33129.74805.23221.93524.33156.53037.41250.52582.63211.4739.2319.23293.7651.41924.61684.6707.94630.52942.72141.7390.82046.41181.81347.31495.6486.22677.10528.91799.84439.52285.12185.43099.7106.81891.23643.401501.94234.1412.21711.42787.81704.2561.62053.81483.9224.7261.2787.3107.02114.01578.11170.8639.51887.8118.4416.6003243.6400.0592.0231.001170.81429.62254.0109.53237.3343.3592.01170.8626.52349.5109.53189.6626.52615.0137.73439.91170.81782.5143.43416.83037.4943.82582.63211.4739.2319.23293.7651.41924.61675.3505.74623.02942.72141.7390.82046.41181.81347.31397.6486.22677.10575.51799.84439.52210.52185.43099.7108.41891.23643.401501.94234.13416.8107553168712184960151948554961926615423409032461298092654610255521238224373621100300832540716451053634304350504304357972063788727143250646370241130296622584559946469523205010953522109132631624688331178184145162214822138671171664510371581093771593773918817192243533109132641624688341178184147162223822131922422831132574312311132312531132192244071224phylum3673.83383.22790.54018.93130.62970.43184.13767.83644.53742.63148.63190.13555.33653.63851.73091.43047.23360.82786.43201.93736.63291.03716.03947.13666.13989.72868.82532.44407.34187.83357.72484.53030.81582.83458.34836.23130.14083.23305.93325.52097.33037.41250.52582.63211.4483.602289.4524.02463.91684.604756.70000616.600000004439.53015.62183.90118.3416.63643.4004234.1412.21711.42787.81704.2524.03211.81483.9000000003434.9108.2416.6000400.0592.000003434.965.90343.3592.0003434.965.9003426.203483.80003467.23037.4943.82582.63211.4483.602289.4524.02463.91675.304756.70000616.600000004439.53019.02183.90103.1416.63643.4004234.13467.2107551168692166560121948443311839515418406468272730926546100872143355337317528300802469997210329335733504742253577382478872712117924636723511129662254525993245452262504693510109132632473331145184162214822136711716451015897971881735101091326424733411451841622238221318817314338855519534421222391262315318817188171236class3673.93383.22798.64020.03130.63459.53740.83869.03645.03750.33286.43293.73555.33668.54142.63050.23465.73362.62779.33202.13762.43489.23760.03966.43666.34015.82869.32629.04407.34187.13790.52484.53061.41824.33458.34864.73130.34124.53307.33325.53879.73037.4692.32582.63211.4002289.402654.01684.604823.70000616.600000004439.502183.90118.3416.63643.4004234.101711.42787.81704.203211.81483.9000000000108.2416.60000592.00000065.900592.000065.900003483.800003037.4692.32582.63211.4002289.402654.01675.304823.70000616.600000004439.502183.90103.1416.63643.4004234.13483.85211716253382352067659644246291174126252278662710191752435375990736135949239106422310642232112721152272274order02488.82810.8882.5003885.03887.52787.7003499.703711.93448.43161.7000004376.405355.54971.74017.92534.73057.74410.34188.53382.24741.84530.03845.03958.13668.43130.04296.13364.13021.02550.90692.30000000003724.30000000000004439.5000003643.4004614.00000000000000000000000000000000000000000000000692.30000000003724.30000000000004439.5000003643.4004614.008235967649644954112161010193589677334111468family00000003887.52804.4003499.303711.93441.33785.60000000004017.91357.800000003724.54849.803361.43363.53880.03045.70000000000000000000000000000000004614.00000000000000000000000000000000000000000000000000000000000000000000000000000004614.0055733553469genus000000001955.6000003520.200000000000000000000000003045.70000000000000000000000000000000004614.00000000000000000000000000000000000000000000000000000000000000000000000000000004614.004331909768species\_group00000000000000000000000000000000000000003931.50000000000000000000000000000000004614.00000000000000000000000000000000000000000000000000000000000000000000000000000004614.00333333470species00000000000000000000000000000000000000004614.00000000000000000000000000000000004614.00000000000000000000000000000000000000000000000000000000000000000000000000000004614.00823546764964454111169101935895726251475genus00000003887.53865.5003499.303711.903785.60000000004018.81357.800000004063.34849.803361.43363.63880.000000000000000000000000000000000000000000000000000000000000000000000000000000000000000000000000000000000000000000009696476species0000000000000852.4000000000000788.500000000000000000000000000000000000000000000000000000000000000000000000000000000000000000000000000000000000000000000000000000000823346764962954109591019358947769446731959053804591019357557480species00000003887.83865.5003499.303716.503785.60000000004019.82424.800000004063.34849.803361.43363.73880.00000000000000000000000000000000000000000000000000000000000000000000000000000000000000000000000000000000000000000000539333930513953933393051391236608no\_rank00000004275.70003434.703871.9000000000004227.70000000000003776.80000000000000000000000000000000000000000000000000000000000000000000000000000000000000000000000000000000000000000000052117162531111529914920786527101817524252759902342511610642210642211135621family02488.82810.8882.5003886.402774.0000003452.70000004376.405395.904033.54626.93323.84410.34188.53827.04741.84530.03845.04051.53230.83129.94348.03799.62645.100692.30000000003724.30000000000004439.5000003643.40000000000000000000000000000000000000000000000000692.30000000003724.30000000000004439.5000003643.40000521171625311115299139207865271018175242527599013425116106422106422150659131231912220227286genus02488.82810.8882.5003886.402774.0000003452.70000004376.405395.904335.94626.93323.84410.34188.53827.04741.84530.03845.04051.53230.83130.04348.03799.62645.100692.30000000003724.30000000000004439.5000003643.40000000000000000000000000000000000000000000000000692.30000000003724.30000000000004439.5000003643.40000421067625256299129177842222906175825275961828151161081081111136841species\_group02418.02813.3882.5003890.103434.70000000000004376.405395.904687.14626.93884.94416.44075.14010.74741.84530.04331.94051.53230.83132.94275.83799.62645.100692.300000000000000000000000000003321.20000000000000000000000000000000000000000000000000692.300000000000000000000000000003321.200004210566252462991281778412229061758252759618273511641861062155629912817758720114958157589961932816287species02418.02814.4882.5003891.603434.70000000000004376.405395.904687.15037.03884.94416.84075.14010.74741.84530.04331.94051.53230.83132.94303.73799.62645.1000000000000000000000000000000000000000000000000000000000000000000000000000000000000000000000000000000000000000000021622162208963no\_rank002922.200000000000000000000000000000000000000000000000000000000000000000000000000000000000000000000000000000000000000000000000000000000000000000000000000000000478478557722no\_rank00000000000000000000000000003828.5004631.2000000000000000000000000000000000000000000000000000000000000000000000000000000000000000000000000000000000000000000000000000369369910265no\_rank0000004505.50000000000000000000000000000000000000000000000000000000000000000000000000000000000000000000000000000000000000000000000000000000000000000000000000000991279007no\_rank00000000000000000000000000005145.200000000000000000000000000000000000000000000000000000000000000000000000000000000000000000000000000000000000000000000000000000049491280938no\_rank00000000000000000000000000005795.100000000000000000000000000000000000000000000000000000000000000000000000000000000000000000000000000000000000000000000000000000010205744523102057445231340851no\_rank00000000000000000000000000000000003931.62873.91867.63048.93813.30000000000000000000000000000000000000000000000000000000000000000000000000000000000000000000000000000000000000000000013131400868no\_rank0000000000000000000000000000000000001353.800000000000000000000000000000000000000000000000000000000000000000000000000000000000000000000000000000000000000000000006516511408272no\_rank003042.000000000000000000000000004079.40000000000000000000000000000000000000000000000000000000000000000000000000000000000000000000000000000000000000000000000000000002781121766352781121766351411700no\_rank002451.500000000000000000000000003718.64853.64010.70000004513.600000000000000000000000000000000000000000000000000000000000000000000000000000000000000000000000000000000000000000000087871415629no\_rank00000000000000000000000000005349.200000000000000000000000000000000000000000000000000000000000000000000000000000000000000000000000000000000000000000000000000000035351427342no\_rank0000000000000000000000000000000000001368.000000000000000000000000000000000000000000000000000000000000000000000000000000000000000000000000000000000000000000000008888853412species00000000000000000000000000000000000003321.20000000000000000000000000000000003321.20000000000000000000000000000000000000000000000000000000000000000000000000000003321.20000881245471no\_rank00000000000000000000000000000000000003321.20000000000000000000000000000000000000000000000000000000000000000000000000000000000000000000000000000000000000000000001010101232139species\_subgroup00692.3000000000000000000000000000000000000000692.3000000000000000000000000000000000000000000000000000000000000000000000000000000692.3000000000000000000000000000000000101010101010301species00692.3000000000000000000000000000000000000000692.3000000000000000000000000000000000000000000000000000000000000000000000000000000692.30000000000000000000000000000000003881136843species\_group002270.30000000000000000000000000000000002150.0000000000000000000000000000000000000000000000000000000000000000000000000000000000000000000000000000000000000000000000038813881294species002270.30000000000000000000000000000000002150.000000000000000000000000000000000000000000000000000000000000000000000000000000000000000000000000000000000000000000000007552136845species\_group00000000000000000000000000000000000003329.30000000000000000000000000000000002617.20000000000000000000000000000000000000000000000000000000000000000000000000000002617.20000555155303species00000000000000000000000000000000000002917.00000000000000000000000000000000002617.20000000000000000000000000000000000000000000000000000000000000000000000000000002617.2000044231023no\_rank00000000000000000000000000000000000002542.20000000000000000000000000000000000000000000000000000000000000000000000000000000000000000000000000000000000000000000004444441306993species000000000000000000000000000004439.5000000000000000000000000000000000004439.50000000000000000000000000000000000000000000000000000000000000000000000000000004439.500000000001661116161427696922583993no\_rank002622.9000000000003724.300000000000003258.44798.90003582.90004842.7000000000000003724.30000000000000000004499.8000000000000000000000000000000000000000000000000000000000003724.30000000000000000004499.800009999991415630species00000000000000000000000000000000000004499.80000000000000000000000000000000004499.80000000000000000000000000000000000000000000000000000000000000000000000000000004499.800001611161614161611161614161978467species002622.900000000000000000000000003258.44798.90003582.90004851.50000000000000000000000000000000000000000000000000000000000000000000000000000000000000000000000000000000000000000000006666662018067species000000000000003724.300000000000000000000000000000000000003724.30000000000000000000000000000000000000000000000000000000000000000000000000000003724.3000000000000000000000001075481683175996194844141539182654611595555165201746348215629587913652184235100132632433113916226105811716103510013264243411391622610211586123135152121191347order3673.93384.94191.44028.73130.63459.52949.703646.903064.003555.304801.73138.03237.21930.42639.60003823.300002805.202340.72812.62483.62173.71752.93456.93612.23494.23076.63759.02800.53345.13037.402761.73211.4002289.4001684.604829.5000000000000002183.9099.300004120.101885.12787.81704.2001483.9000000000000000000000000000000000000003037.402761.73211.4002289.4001675.304829.5000000000000002183.9099.300004120.10107546166606126719445413153518141156455201746347194429586913652084235132348911391622101031035132358101139162210971113240429141245521361434431131543family3674.03389.24097.72731.13132.63459.52925.703650.403064.001567.904804.403274.51930.40000000002805.202340.72812.62483.72391.92202.73457.03612.23494.23076.63941.72800.53345.13037.4003214.1003459.6001496.904829.5000000000000002183.90000004120.1002903.90001483.9000000000000000000000000000000000000003037.4003214.0003459.6001483.904829.5000000000000002183.90000004120.10612643295811449181139918113975544genus04055.002730.93287.40003459.6000004821.5000000000000000000000000000002855.7003459.600004829.500000000000000000000000000000000000000000000000000000000000000000000002855.7003459.600004829.500000000000000000000000113911391139113911391139545species000000000000004829.500000000000000000000000000000000000004829.50000000000000000000000000000000000000000000000000000000000000000000000000000004829.50000000000000000000000061264326987387381121344959species\_group04055.002730.93290.30003459.6000000000000000000000000000000000002907.8003459.60000000000000000000000000000000000000000000000000000000000000000000000000002907.8003459.60000000000000000000000000000612633184612633184546species04055.002730.53303.100000000000000000000000000000000000000000000000000000000000000000000000000000000000000000000000000000000000000000000000000000000000000000000000000000088888867827species00001409.50000000000000000000000000000000000000001409.50000000000000000000000000000000000000000000000000000000000000000000000000000001409.500000000000000000000000000000003131313131311639133species00002151.30000000000000000000000000000000000000002151.30000000000000000000000000000000000000000000000000000000000000000000000000000002151.300000000000000000000000000000003434343434342077147species00003950.00000000000000000000000000000000000000003950.00000000000000000000000000000000000000000000000000000000000000000000000000000003950.000000000000000000000000000000008888882529121species000000003459.6000000000000000000000000000000000000003459.60000000000000000000000000000000000000000000000000000000000000000000000000000003459.6000000000000000000000000000019181812644389no\_rank00002725.20000000000000000000000000000000000000002644.60000000000000000000000000000000000000000000000000000000000000000000000000000002644.600000000000000000000000000000001313131313131703250species00002108.80000000000000000000000000000000000000002108.80000000000000000000000000000000000000000000000000000000000000000000000000000002108.800000000000000000000000000000005555551920110species00004037.80000000000000000000000000000000000000004037.80000000000000000000000000000000000000000000000000000000000000000000000000000004037.800000000000000000000000000000006014134737331547genus02708.4002287.90002832.3000000000000000000000000000000000002845.40000000000000000000000000000000000000000000000000000000000000000000000000000002845.4000000000000000000000000000000057102343838262354276species\_group02838.4002150.90002832.3000000000000000000000000000000000002925.40000000000000000000000000000000000000000000000000000000000000000000000000000002925.400000000000000000000000000000003232732327550species03186.9001733.10002809.70000000000000000000000000000000000000000000000000000000000000000000000000000000000000000000000000000000000000000000000000000000000000000000000000017172517176158836species02814.2001668.30002860.3000000000000000000000000000000000000000000000000000000000000000000000000000000000000000000000000000000000000000000000000000000000000000000000000001919301105subspecies000000002778.000000000000000000000000000000000000000000000000000000000000000000000000000000000000000000000000000000000000000000000000000000000000000000000000000555555299767species00002258.80000000000000000000000000000000000000002258.80000000000000000000000000000000000000000000000000000000000000000000000000000002258.800000000000000000000000000000003333333333331812935species00003026.50000000000000000000000000000000000000003026.50000000000000000000000000000000000000000000000000000000000000000000000000000003026.5000000000000000000000000000000069691915310species01314.8001462.8000000000000000000000000000000000000000000000000000000000000000000000000000000000000000000000000000000000000000000000000000000000000000000000000000000181818181818881260species00002567.60000000000000000000000000000000000000002567.60000000000000000000000000000000000000000000000000000000000000000000000000000002567.6000000000000000000000000000000020171732608935no\_rank00002710.50000000000000000000000000000000000000002960.80000000000000000000000000000000000000000000000000000000000000000000000000000002960.800000000000000000000000000000001717171717171560339species00002960.80000000000000000000000000000000000000002960.80000000000000000000000000000000000000000000000000000000000000000000000000000002960.80000000000000000000000000000000107410196610913152577511111011615412905781262207114561genus3673.73004.14097.702477.102925.703652.603345.700000000000000002805.202499.13097.82208.12335.92256.23460.34053.83513.23155.23941.73166.92831.3000000000000000000000000000000000000000000000000000000000000000000000000000000000000000000000000000000000000000000107410196610913152577511111011615412905381262207112902119661093151837511119385154124359354912211562species3673.73004.14097.702477.102925.703652.603345.700000000000000002805.202499.13097.82208.12335.92256.23460.34053.83513.23155.23941.73166.92831.3000000000000000000000000000000000000000000000000000000000000000000000000000000000000000000000000000000000000000000101811083333no\_rank0000002821.60000000000000000000000002262.7000000000000000000000000000000000000000000000000000000000000000000000000000000000000000000000000000000000000000000000000000181181511145no\_rank00000000000000000000000000000002262.7000000000000000000000000000000000000000000000000000000000000000000000000000000000000000000000000000000000000000000000000000243243199310no\_rank00000000000000000000000000000000004960.80000000000000000000000000000000000000000000000000000000000000000000000000000000000000000000000000000000000000000000000004831748317316435no\_rank000000003938.500000000000000000000000005225.800000000000000000000000000000000000000000000000000000000000000000000000000000000000000000000000000000000000000000000000055409438no\_rank0000000000000000000000000000000000666.80000000000000000000000000000000000000000000000000000000000000000000000000000000000000000000000000000000000000000000000001111439184no\_rank000000003995.5000000000000000000000000000000000000000000000000000000000000000000000000000000000000000000000000000000000000000000000000000000000000000000000000002020566546no\_rank00000000000000000000000000000000001537.30000000000000000000000000000000000000000000000000000000000000000000000000000000000000000000000000000000000000000000000001313595495no\_rank00000000000000000000000000000000000002129.500000000000000000000000000000000000000000000000000000000000000000000000000000000000000000000000000000000000000000000010081008655817no\_rank00000000000000000000000000000000004247.60000000000000000000000000000000000000000000000000000000000000000000000000000000000000000000000000000000000000000000000004685037no\_rank00000000000000000000000000000000001519.500000000000000000000000000000000000000000000000000000000000000000000000000000000000000000000000000000000000000000000000044685038no\_rank00000000000000000000000000000000001519.50000000000000000000000000000000000000000000000000000000000000000000000000000000000000000000000000000000000000000000000001997519975885275no\_rank00000000000000000000000000000000004836.803898.6003226.8000000000000000000000000000000000000000000000000000000000000000000000000000000000000000000000000000000000000000000010441044885276no\_rank00000000000000000000000000000000004193.40000000000000000000000000000000000000000000000000000000000000000000000000000000000000000000000000000000000000000000000001111941280no\_rank4101.1000000000000000000000000000000000000000000000000000000000000000000000000000000000000000000000000000000000000000000000000000000000000000000000000000000000055941322no\_rank6150.800000000000000000000000000000000000000000000000000000000000000000000000000000000000000000000000000000000000000000000000000000000000000000000000000000000005959941323no\_rank000000001474.000000000000000000000002908.700000000000000000000000000000000000000000000000000000000000000000000000000000000000000000000000000000000000000000000000000013441038927no\_rank2640.400000003681.50000000000000000000000000000000000000000000000000000000000000000000000000000000000000000000000000000000000000000000000000000000000000000000000000013131134782no\_rank2640.4000000000000000000000000000000000000000000000000000000000000000000000000000000000000000000000000000000000000000000000000000000000000000000000000000000000051072458no\_rank2338.40000000000000000000000000000000000000000000000000000000000000000000000000000000000000000000000000000000000000000000000000000000000000000000000000000000000551072459no\_rank2338.4000000000000000000000000000000000000000000000000000000000000000000000000000000000000000000000000000000000000000000000000000000000000000000000000000000000085851274814no\_rank3039.2000000000000000000000000000000000000000000000000000000000000000000000000000000000000000000000000000000000000000000000000000000000000000000000000000000000058581322345no\_rank000000000000000000000000000000000004251.6004124.8000000000000000000000000000000000000000000000000000000000000000000000000000000000000000000000000000000000000000000007782767782761355100no\_rank3644.000000003323.00000000000000000000000000000000000000000000000000000000000000000000000000000000000000000000000000000000000000000000000000000000000000000000000000045451355101no\_rank4298.400000000000000000000000000000000000000000000000000000000000000000000000000000000000000000000000000000000000000000000000000000000000000000000000000000000001441441392854no\_rank00000000000000000000000000000002049.0003808.000000000000000000000000000000000000000000000000000000000000000000000000000000000000000000000000000000000000000000000000058581401688no\_rank2630.2000000000000000000000000000000000000000000000000000000000000000000000000000000000000000000000000000000000000000000000000000000000000000000000000000000000018181502658no\_rank00000000000000000000000000000000003688.600000000000000000000000000000000000000000000000000000000000000000000000000000000000000000000000000000000000000000000000017171954351no\_rank00000000000000000000000000000000002911.3000000000000000000000000000000000000000000000000000000000000000000000000000000000000000000000000000000000000000000000000527181552718152126982no\_rank00000000000000000000000000000002733.0003630.30000000000000000000000000000000000000000000000000000000000000000000000000000000000000000000000000000000000000000000000003403402592065no\_rank3976.0000000000000000000000000000000000000000000000000000000000000000000000000000000000000000000000000000000000000000000000000000000000000000000000000000000000024162371554119220221412962141296214520623570genus4375.03401.4003123.20003747.300000000000000000000002470.8003107.60000003081.1003221.50000000000000000000000000000000000000000000000000000000000000000000000000003081.1003221.50000000000000000000000000000000141414141414548species03081.10000000000000000000000000000000000000003081.10000000000000000000000000000000000000000000000000000000000000000000000000000003081.100000000000000000000000000000000001023510235571species03215.2002046.100000000000000000000000000000000000000000000000000000000000000000000000000000000000000000000000000000000000000000000000000000000000000000000000000000015160599651222019151504994712919573species4879.73407.5002780.80003658.800000000000000000000002470.8003153.600000000000000000000000000000000000000000000000000000000000000000000000000000000000000000000000000000000000000000000000099574subspecies03141.900000000000000000000000000000000000000000000000000000000000000000000000000000000000000000000000000000000000000000000000000000000000000000000000000000000089318211767321172407subspecies03327.7002703.5000000000000000000000000002493.60000000000000000000000000000000000000000000000000000000000000000000000000000000000000000000000000000000000000000000000000005050272620no\_rank03138.000000000000000000000000000000000000000000000000000000000000000000000000000000000000000000000000000000000000000000000000000000000000000000000000000000000026261123862no\_rank03339.6000000000000000000000000000000000000000000000000000000000000000000000000000000000000000000000000000000000000000000000000000000000000000000000000000000000361536151328324no\_rank02855.8003159.900000000000000000000000000000000000000000000000000000000000000000000000000000000000000000000000000000000000000000000000000000000000000000000000000000014141392499no\_rank03797.900000000000000000000000000000000000000000000000000000000000000000000000000000000000000000000000000000000000000000000000000000000000000000000000000000000015151049565no\_rank04039.300000000000000000000000000000000000000000000000000000000000000000000000000000000000000000000000000000000000000000000000000000000000000000000000000000000076761284798no\_rank03307.9000000000000000000000000000000000000000000000000000000000000000000000000000000000000000000000000000000000000000000000000000000000000000000000000000000000661380908no\_rank04559.700000000000000000000000000000000000000000000000000000000000000000000000000000000000000000000000000000000000000000000000000000000000000000000000000000000011111392500no\_rank03642.200000000000000000000000000000000000000000000000000000000000000000000000000000000000000000000000000000000000000000000000000000000000000000000000000000000083128312244366species02641.3001942.2000000000000000000000000000000000000000000000000000000000000000000000000000000000000000000000000000000000000000000000000000000000000000000000000000000816962119451134687species3534.03256.1002704.00000000000000000000000000000000000000000000000000000000000000000000000000000000000000000000000000000000000000000000000000000000000000000000000000000001551551006551no\_rank03416.0002255.6000000000000000000000000000000000000000000000000000000000000000000000000000000000000000000000000000000000000000000000000000000000000000000000000000000771191061no\_rank3858.0000000000000000000000000000000000000000000000000000000000000000000000000000000000000000000000000000000000000000000000000000000000000000000000000000000000012121308980no\_rank00002054.800000000000000000000000000000000000000000000000000000000000000000000000000000000000000000000000000000000000000000000000000000000000000000000000000000041014921463165species04082.5002681.8000000000000000000000000000000000000000000000000000000000000000000000000000000000000000000000000000000000000000000000000000000000000000000000000000000991667327subspecies00002082.70000000000000000000000000000000000000000000000000000000000000000000000000000000000000000000000000000000000000000000000000000000000000000000000000000006060606060602058152species00002196.80000000000000000000000000000000000000002196.80000000000000000000000000000000000000000000000000000000000000000000000000000002196.800000000000000000000000000000006666662153354species00001482.50000000000000000000000000000000000000001482.50000000000000000000000000000000000000000000000000000000000000000000000000000001482.5000000000000000000000000000000061299451289612896632608929no\_rank03630.0003219.70004250.8000000000000000000000000000000000003227.10000000000000000000000000000000000000000000000000000000000000000000000000000003227.100000000000000000000000000000001151151151151151151905288species00002998.20000000000000000000000000000000000000002998.20000000000000000000000000000000000000000000000000000000000000000000000000000002998.200000000000000000000000000000007575757575751934254species00002258.60000000000000000000000000000000000000002258.60000000000000000000000000000000000000000000000000000000000000000000000000000002258.600000000000000000000000000000001270612706127061270612706127062488567species00003234.80000000000000000000000000000000000000003234.80000000000000000000000000000000000000000000000000000000000000000000000000000003234.800000000000000000000000000000009559552697371species00002293.30004250.8000000000000000000000000000000000000000000000000000000000000000000000000000000000000000000000000000000000000000000000000000000000000000000000000001550161342161192161191590genus2282.12882.2002355.2000000000000000000000000002075.00000000003008.2002388.700000000000000000000002114.200000000000000000000000000000000000000000000000000003008.2002388.700000000000000000000002114.20000000014291015914328901species2409.12790.9002335.2000000000000000000000000001764.200000000000000000000000000000000000000000000000000000000000000000000000000000000000000000000000000000000000000000000000000052861256359201subspecies3528.02790.1003109.0000000000000000000000000001602.90000000000000000000000000000000000000000000000000000000000000000000000000000000000000000000000000000000000000000000000000005598360no\_rank3528.0000000000000000000000000000000000000000000000000000000000000000000000000000000000000000000000000000000000000000000000000000000000000000000000000000000000099149539no\_rank00000000000000000000000000000001957.30000000000000000000000000000000000000000000000000000000000000000000000000000000000000000000000000000000000000000000000000002323260678no\_rank02889.60000000000000000000000000000000000000000000000000000000000000000000000000000000000000000000000000000000000000000000000000000000000000000000000000000000006666654736species00002388.70000000000000000000000000000000000000002388.70000000000000000000000000000000000000000000000000000000000000000000000000000002388.70000000000000000000000000000000661197719no\_rank00002388.70000000000000000000000000000000000000000000000000000000000000000000000000000000000000000000000000000000000000000000000000000000000000000000000000000002111921119211192614656no\_rank03008.2000000000000000000000000000002114.20000000003008.200000000000000000000000002114.200000000000000000000000000000000000000000000000000003008.200000000000000000000000002114.2000000002121212121212500543species03008.20000000000000000000000000000000000000003008.20000000000000000000000000000000000000000000000000000000000000000000000000000003008.200000000000000000000000000000000001191191191191191192664291species00000000000000000000000000000002114.2000000000000000000000000000000000002114.20000000000000000000000000000000000000000000000000000000000000000000000000000002114.200000000444345166431150310150310411620genus000000004675.0000000000000000000001493.52950.02574.3003565.1000003527.4000000000000000000000000002189.50000004120.1000000000000000000000000000000000000000000000000000000000000000000000002189.50000004120.10464464622species000000004675.000000000000000000000000003565.10000000000000000000000000000000000000000000000000000000000000000000000000000000000000000000000000000000000000000000000004330122043301220623species0000000000000000000000000000002950.02593.6000000003280.4000000000000000000000000000000000000000000000000000000000000000000000000000000000000000000000000000000000000000000150315031503140015031503624species00000000000000000000000000000002152.1000000000000000000000000000000000002189.50000000000000000000000000000000000000000000000000000000000000000000000000000002189.500000000103103300269no\_rank00000000000000000000000000000002696.80000000000000000000000000000000000000000000000000000000000000000000000000000000000000000000000000000000000000000000000000001010102629414no\_rank00000000000000000000000000000000000000004120.10000000000000000000000000000000004120.10000000000000000000000000000000000000000000000000000000000000000000000000000004120.101010101010101813821species00000000000000000000000000000000000000004120.10000000000000000000000000000000004120.10000000000000000000000000000000000000000000000000000000000000000000000000000004120.10939293931931160674genus00002792.40000000000000000000000000000000000000002821.2000000000000000000000000000000002820.80000000000000000000000000000000000000000000002820.80000000000000000000000000000000161616161616577species0000747.5000000000000000000000000000000000000000747.5000000000000000000000000000000000000000000000000000000000000000000000000000000747.5000000000000000000000000000000070707070707054291species00003239.70000000000000000000000000000000000000003239.70000000000000000000000000000000000000000000000000000000000000000000000000000003239.700000000000000000000000000000006666661259973species00003467.30000000000000000000000000000000000000003467.30000000000000000000000000000000000000000000000000000000000000000000000000000003467.3000000000000000000000000000000010910101101413496genus0000000000001366.600000000000000000000000000000000000001496.90000000000000000000000000000001483.9000000000000000000000000000000000000000000000001483.900000000000000000000000009999928141species0000000000001496.900000000000000000000000000000000000001496.90000000000000000000000000000000000000000000000000000000000000000000000000000001496.9000000000000000000000000099290339no\_rank0000000000001496.9000000000000000000000000000000000000000000000000000000000000000000000000000000000000000000000000000000000000000000000000000000000000000000000010101010101330547genus00003676.60000000000000000000000000000000000000003676.6000000000000000000000000000000003676.60000000000000000000000000000000000000000000003676.600000000000000000000000000000001010101010283686species00003676.60000000000000000000000000000000000000003676.60000000000000000000000000000000000000000000000000000000000000000000000000000003676.6000000000000000000000000000000010101177180no\_rank00003676.600000000000000000000000000000000000000000000000000000000000000000000000000000000000000000000000000000000000000000000000000000000000000000000000000000058145814581458141903409family0001885.11933.5000000000000000000000000000000000000001885.11933.500000000000000000000000000000001885.11933.5000000000000000000000000000000000000000000001885.11933.50000000000000000000000000000000585858585853335genus0001885.10000000000000000000000000000000000000001885.1000000000000000000000000000000001885.10000000000000000000000000000000000000000000001885.100000000000000000000000000000000585858585866269species0001885.10000000000000000000000000000000000000001885.10000000000000000000000000000000000000000000000000000000000000000000000000000001885.1000000000000000000000000000000005866271subspecies0001885.100000000000000000000000000000000000000000000000000000000000000000000000000000000000000000000000000000000000000000000000000000000000000000000000000000005858660596no\_rank0001885.1000000000000000000000000000000000000000000000000000000000000000000000000000000000000000000000000000000000000000000000000000000000000000000000000000000014141414142100764genus00001933.50000000000000000000000000000000000000001933.5000000000000000000000000000000001933.50000000000000000000000000000000000000000000001933.500000000000000000000000000000001414141414141458355species00001933.50000000000000000000000000000000000000001933.50000000000000000000000000000000000000000000000000000000000000000000000000000001933.50000000000000000000000000000000171451319265121442155421552211903411family02968.904468.92187.700000003558.30000000003972.6000000000000000000003972.42047.3000004482.8000000000000000000000000000000000000000000000000000000000000000000000003972.42047.3000004482.8000000000000000000000000017145111726511144215542155822613genus02968.904470.52034.400000003558.40000000003972.6000000000000000000003972.42047.3000004482.8000000000000000000000000000000000000000000000000000000000000000000000003972.42047.3000004482.80000000000000000000000000171436926472146137062638814615species02968.904496.7000000003562.10000000003972.6000000000000000000000000000000000000000000000000000000000000000000000000000000000000000000000000000000000000000000000000000000000000663211759subspecies0005114.30000000000000000000000000000000000000000000000000000000000000000000000000000000000000000000000000000000000000000000000000000000000000000000000000000000663663273526no\_rank0005114.30000000000000000000000000000000000000000000000000000000000000000000000000000000000000000000000000000000000000000000000000000000000000000000000000000000258258435998no\_rank0988.300000000003783.5000000000000000000000000000000000000000000000000000000000000000000000000000000000000000000000000000000000000000000000000000000000000000000000069691334564no\_rank04068.300000000002250.40000000000000000000000000000000000000000000000000000000000000000000000000000000000000000000000000000000000000000000000000000000000000000000000796779671401254no\_rank03169.900000000003467.80000000000000000000000000000000000000000000000000000000000000000000000000000000000000000000000000000000000000000000000000000000000000000000000131313131328151species0003676.80000000000000000000000000000000000000003676.80000000000000000000000000000000000000000000000000000000000000000000000000000003676.8000000000000000000000000000000001313399741no\_rank0003676.8000000000000000000000000000000000000000000000000000000000000000000000000000000000000000000000000000000000000000000000000000000000000000000000000000000091316882996species0003577.400000000425.2000000000000000000000000000000000000000000000000000000000000000000000000000000000000000000000000000000000000000000000000000000000000000000000088768492no\_rank0003878.5000000000000000000000000000000000000000000000000000000000000000000000000000000000000000000000000000000000000000000000000000000000000000000000000000000031311154756no\_rank000000000000425.2000000000000000000000000000000000000000000000000000000000000000000000000000000000000000000000000000000000000000000000000000000000000000000000015151348660no\_rank0003632.70000000000000000000000000000000000000000000000000000000000000000000000000000000000000000000000000000000000000000000000000000000000000000000000000000000301562915529155112647522no\_rank0004125.92047.300000004048.70000000000000000000000000000004104.92047.3000004482.8000000000000000000000000000000000000000000000000000000000000000000000004104.92047.3000004482.80000000000000000000000000111111111111768490species0003733.70000000000000000000000000000000000000003733.70000000000000000000000000000000000000000000000000000000000000000000000000000003733.700000000000000000000000000000000666666768493species0004195.00000000000000000000000000000000000000004195.00000000000000000000000000000000000000000000000000000000000000000000000000000004195.0000000000000000000000000000000005555551327989species0000000000004482.800000000000000000000000000000000000004482.80000000000000000000000000000000000000000000000000000000000000000000000000000004482.800000000000000000000000001212121212122420306species0004400.10000000000000000000000000000000000000004400.10000000000000000000000000000000000000000000000000000000000000000000000000000004400.1000000000000000000000000000000001515151515152447890species00002047.30000000000000000000000000000000000000002047.30000000000000000000000000000000000000000000000000000000000000000000000000000002047.30000000000000000000000000000000171912161961616196131903414family000000001931.60001037.200000000000000000000104.000000000000001704.2001037.200000000000000000099.3000000001704.2000000000000000000000000000000000000000000000001704.2001037.200000000000000000099.30000001616161616581genus000000001704.2000000000000000000000000000000000000001704.20000000000000000000000000000001704.2000000000000000000000000000000000000000000000001704.20000000000000000000000000000161616161616582species000000001704.2000000000000000000000000000000000000001704.20000000000000000000000000000000000000000000000000000000000000000000000000000001704.200000000000000000000000000001991961963583genus0000000000001037.20000000000000000000087.100000000000000001037.200000000000000000099.3000000000000000000000000000000000000000000000000000000000001037.200000000000000000099.300000077777584species0000000000001072.600000000000000000000000000000000000001072.60000000000000000000000000000000000000000000000000000000000000000000000000000001072.6000000000000000000000000077529507no\_rank0000000000001072.60000000000000000000000000000000000000000000000000000000000000000000000000000000000000000000000000000000000000000000000000000000000000000000000121212121212585species0000000000001016.500000000000000000000000000000000000001016.50000000000000000000000000000000000000000000000000000000000000000000000000000001016.50000000000000000000000000666666626774species00000000000000000000000000000000099.30000000000000000000000000000000000099.300000000000000000000000000000000000000000000000000000000000000000000000000000099.3000000451145845104510135614order000000000003211.8000000000000000000000125.80000000000000003211.80000000000000000000132.600000000003211.80000000000131.40000000000000000000000000000000000003211.80000000000000000000131.4000000451145845104510132033family000000000003211.8000000000000000000000125.80000000000000003211.80000000000000000000132.600000000003211.80000000000131.40000000000000000000000000000000000003211.80000000000000000000131.400000010810102102338genus000000000000000000000000000000000126.600000000000000000000000000000000000132.6000000000000000000000131.400000000000000000000000000000000000000000000000000000000131.4000000888388339species000000000000000000000000000000000129.500000000000000000000000000000000000132.6000000000000000000000000000000000000000000000000000000000000000000000000000000132.60000005359385no\_rank000000000000000000000000000000000134.4000000000000000000000000000000000000000000000000000000000000000000000000000000000000000000000000000000000000000000000000055990315no\_rank000000000000000000000000000000000134.40000000000000000000000000000000000000000000000000000000000000000000000000000000000000000000000000000000000000000000000000454545454540323genus000000000003211.800000000000000000000000000000000000003211.80000000000000000000000000000003211.8000000000000000000000000000000000000000000000003211.800000000000000000000000000454545995085species\_group000000000003211.800000000000000000000000000000000000003211.80000000000000000000000000000000000000000000000000000000000000000000000000000003211.80000000000000000000000000045454545454540324species000000000003211.800000000000000000000000000000000000003211.80000000000000000000000000000000000000000000000000000000000000000000000000000003211.800000000000000000000000000551159618135615order00000002280.80000000000002844.82216.4000001343.5001342.700001570.100000000000000000000000000000000000000000000000000000000000000000000000000000000000000000000000000000000000000000000000551159618868family00000002280.80000000000002844.82216.4000001343.5001342.700001570.1000000000000000000000000000000000000000000000000000000000000000000000000000000000000000000000000000000000000000000000005511596182717genus00000002280.80000000000002844.82216.4000001343.5001342.700001570.1000000000000000000000000000000000000000000000000000000000000000000000000000000000000000000000000000000000000000000000005511596185511596182718species00000002280.80000000000002844.82216.4000001343.5001342.700001570.100000000000000000000000000000000000000000000000000000000000000000000000000000000000000000000000000000000000000000000000757777135623order00000000000000000000000000000000075.700000000000000000000000000000000000000000000000000000000062.00000000000065.900000065.9000000000000000000000000000000000000065.90000007577272641family00000000000000000000000000000000075.700000000000000000000000000000000000000000000000000000000062.00000000000065.90000000000000000000000000000000000000000000065.9000000555555662genus00000000000000000000000000000000062.000000000000000000000000000000000000000000000000000000000062.00000000000000000000000000000000000000000000000000000000062.00000009999999135624order000592.0000000000000000000000000000000000000000592.000000000000000000000000000000000592.000000000000000000000592.000000000592.0000000000000000592.00000000000000000000000000000000099999984642family000592.0000000000000000000000000000000000000000592.000000000000000000000000000000000592.000000000000000000000592.0000000000000000000000000592.00000000000000000000000000000000099999642genus000592.0000000000000000000000000000000000000000592.000000000000000000000000000000000592.0000000000000000000000000000000000000000000000592.000000000000000000000000000000000999999654species000592.0000000000000000000000000000000000000000592.0000000000000000000000000000000000000000000000000000000000000000000000000000000592.000000000000000000000000000000000334877601524406388112049443895524473731251430080246899291031333558350369735747736191760206428253821025162771941528188828188135625order02637.82249.20003292.52934.83078.83750.13320.33225.802749.83377.83004.03495.13362.82813.23202.13763.13484.53759.93966.13666.23965.92869.42724.43239.403810.303159.12299.05158.34869.54735.94241.23180.83915.13966.4000000001757.50000000616.6000000000000416.6000000000000000000000416.6000000000000000000000000000000000001757.50000000616.6000000000000416.6000003348776015244063881120494438955244737312514300802468992910313335583503697357477361917602064282538210251627719415281888281881232116132714124584521712family02637.82249.20003292.52934.83078.83750.13320.33225.802749.83377.83004.03495.13362.82813.23202.13763.13484.53759.93966.13666.23965.92869.42724.43239.403810.303159.12299.05158.34869.54735.94241.23180.83915.13966.4000000001757.50000000616.6000000000000416.6000000000000000000000416.6000000000000000000000000000000000001757.50000000616.6000000000000416.60000012244818818464713genus00000000000378.300000000605.6000000850.8000000000000000000000397.50000000616.60000000000000000000000000000000000000000000000000000000000000000000000397.50000000616.6000000000000000000888888715species00000000000000000000274.2000000000000000000000000000000000000274.2000000000000000000000000000000000000000000000000000000000000000000000000000000274.2000000000000000000888888716species00000000000397.50000000000000000000000000000000000000397.5000000000000000000000000000000000000000000000000000000000000000000000000000000397.500000000000000000000000000101010101010189834species00000000000000000000890.5000000000000000000000000000000000000890.5000000000000000000000000000000000000000000000000000000000000000000000000000000890.50000000000000000003248571710844046681020335201947244737310406300802428592810311335363503197357426411917522062825329102516258194142020422642761251291267104018311643257712724genus02656.42253.40003376.23241.93078.83763.33321.83238.403106.43400.93004.03495.13362.93243.83202.13809.73486.83760.53967.93666.33965.92869.72889.13239.403808.803159.105158.34876.64735.94241.23183.63915.13975.7000000002301.60000000000000000000000000000000000000000000000000000000000000000000000000000002301.6000000000000000000000000009617307871051971891311138896173078710519718913111388726species001677.30000001205.31642.81904.903131.51879.400002878.52710.82031.203309.60002433.900002210.4003864.60000000000000000000000000000000000000000000000000000000000000000000000000000000000000000000000000000000000000000000000042551039789233285372271342889121222981031130167350309135738156141752528244621025162511841242191039546182325209771342801528058010018300863422786355351421417525282313110251546213394727species001264.50003864.92766.703775.13316.13345.500001121.23366.93258.83210.13911.43193.63760.54047.03666.34020.12869.83001.13100.703808.801291.405158.34911.44735.94241.23183.73835.83984.00000000000000000000000000000000000000000000000000000000000000000000000000000000000000000000000000000000000000000007619726305876197263058725no\_rank0000000002260.300000002013.502713.0003767.84976.8000000000004903.5003257.7000000000000000000000000000000000000000000000000000000000000000000000000000000000000000000000000000000000000000000005571421no\_rank000000000000000000000000000000000005240.400000000000000000000000000000000000000000000000000000000000000000000000000000000000000000000000000000000000000000000000301452120511301452120511262727no\_rank0000004025.500000000003212.503608.00000000000000000004224.7000000000000000000000000000000000000000000000000000000000000000000000000000000000000000000000000000000000000000000005101451538902851014515389028262728no\_rank00000000003740.84066.8000003054.203827.63202.703194.20000000000005825.3003693.20000000000000000000000000000000000000000000000000000000000000000000000000000000000000000000000000000000000000000000067825161016613803128146782516101661380312814281310no\_rank0000003636.7003806.503500.3000003264.603288.0005314.803968.603193.2000000006093.2000000000000000000000000000000000000000000000000000000000000000000000000000000000000000000000000000000000000000000000002418396624183966374930no\_rank00000000000000000003114.83786.303311.300000000000000000000000000000000000000000000000000000000000000000000000000000000000000000000000000000000000000000000000000000000000053958142495395814249374931no\_rank000000000003390.2000000000002858.903738.42365.22852.400000005108.400000000000000000000000000000000000000000000000000000000000000000000000000000000000000000000000000000000000000000000000825689825689862964no\_rank000000000001535.9000003767.400002514.70000000000003819.0002165.800000000000000000000000000000000000000000000000000000000000000000000000000000000000000000000000000000000000000000000812812866630no\_rank000000000000000000000000000000000005241.2003488.70000000000000000000000000000000000000000000000000000000000000000000000000000000000000000000000000000000000000000000096111781509611178150935897no\_rank0000000003368.900000000002523.7000001491.5000000003822.2002269.20000000000000000000000000000000000000000000000000000000000000000000000000000000000000000000000000000000000000000000014956280576431175149562805764311751232659no\_rank0000000004256.200000003751.003768.1004099.40001941.7000000005708.0003713.64027.0000000000000000000000000000000000000000000000000000000000000000000000000000000000000000000000000000000000000000000097665060840411976650608404111295140no\_rank00000000000000000003137.30003837.1003086.4000000004810.0003461.404287.90000000000000000000000000000000000000000000000000000000000000000000000000000000000000000000000000000000000000000001880181019871880181019871334187no\_rank000000000000000003496.103623.103531.73620.00000000000005554.300004895.700000000000000000000000000000000000000000000000000000000000000000000000000000000000000000000000000000000000000000028461460926357641920092917244271232266356731864015188377182784216041465617947585742442141188234618392315170241729species02752.92270.90003113.03357.303110.03370.03265.103117.53430.93004.03777.71494.73336.103137.23598.103269.30002916.63627.80003275.1004256.60000000000000000000000000000000000000000000000000000000000000000000000000000000000000000000000000000000000000000000000010183393222110812533434371917812211347170181361018339322211081253343437191781221134717018136862965no\_rank02758.52262.30002800.53016.703271.83164.72944.702954.43463.5001494.73231.303294.53505.703284.20002985.600003016.8004240.300000000000000000000000000000000000000000000000000000000000000000000000000000000000000000000000000000000000000000000000202020202020730species000000000002301.600000000000000000000000000000000000002301.60000000000000000000000000000000000000000000000000000000000000000000000000000002301.6000000000000000000000000001119263139243211192631392432735species000000000000002786.50002339.403039.93349.3000002200.000000001561.80000000000000000000000000000000000000000000000000000000000000000000000000000000000000000000000000000000000000000000000094289139428913197575species0000000002855.70000000002734.40004454.8000000000004883.4000000000000000000000000000000000000000000000000000000000000000000000000000000000000000000000000000000000000000000000002824928249249188species00000000000901.4000000000001977.90003086.00000000000000000000000000000000000000000000000000000000000000000000000000000000000000000000000000000000000000000000000000000000932710158828172609962no\_rank002512.0000000003346.603417.200003577.7003722.003474.50002919.800000000000000000000000000000000000000000000000000000000000000000000000000000000000000000000000000000000000000000000000000000009327101588281793271015882817712310species002512.0000000003346.603417.200003577.7003722.003474.50002919.80000000000000000000000000000000000000000000000000000000000000000000000000000000000000000000000000000000000000000000000000000000140155462111114745genus000000000345.4000366.9312.8000326.60373.900765.6000586.30000000000000000000000000000000000000000000000000000000000000000000000000000000000000000000000000000000000000000000000000000000140155462111114140155462111114747species000000000345.4000366.9312.8000326.60373.900765.6000586.30000000000000000000000000000000000000000000000000000000000000000000000000000000000000000000000000000000000000000000000000000000782375984genus00000000000766.3000000001171.90000000000000000000000000000000000000000000000000000000000000000000000000000000000000000000000000000000000000000000000000000000000000055575985species00000000000641.6000000001651.800000000000000000000000000000000000000000000000000000000000000000000000000000000000000000000000000000000000000000000000000000000000000551222034no\_rank000000000000000000001651.8000000000000000000000000000000000000000000000000000000000000000000000000000000000000000000000000000000000000000000000000000000000000001210214906genus000000000123.7000487.20000000000000000000000000000000000000000000000000000000000000000000000000000000000000000000000000000000000000000000000000000000000000000000001210731species000000000123.7000487.200000000000000000000000000000000000000000000000000000000000000000000000000000000000000000000000000000000000000000000000000000000000000000000012101210205914no\_rank000000000123.7000487.2000000000000000000000000000000000000000000000000000000000000000000000000000000000000000000000000000000000000000000000000000000000000000000000404219134211481477562840171281117541416916genus0000001955.22237.603215.901807.502699.200002275.201816.1001889.02552.4002204.6004154.800001840.000874.9000000000000000000000000000000000000000000000000000000000000000000000000000000000000000000000000000000000000000000004451744517714species000000000001840.6000000001375.000000000000000000874.900000000000000000000000000000000000000000000000000000000000000000000000000000000000000000000000000000000000000000000730195761299188227303111914822732species0000001531.32412.403215.901956.502998.300002409.201792.90000001999.1004154.80000862.4000000000000000000000000000000000000000000000000000000000000000000000000000000000000000000000000000000000000000000000001281412814634176no\_rank000000000001959.8000000001560.40000002076.2000000000000000000000000000000000000000000000000000000000000000000000000000000000000000000000000000000000000000000000000000000019145111914511985008no\_rank0000000003215.902721.703479.200002373.9000000000000000000000000000000000000000000000000000000000000000000000000000000000000000000000000000000000000000000000000000000000000000032102520435424017739species0000002063.72016.60001591.702691.900002237.701960.50000002364.300000003135.2000000000000000000000000000000000000000000000000000000000000000000000000000000000000000000000000000000000000000000000003210252043542401732102520435424017888057no\_rank0000002063.72016.60001591.702691.900002237.701960.50000002364.300000003135.200000000000000000000000000000000000000000000000000000000000000000000000000000000000000000000000000000000000000000000000888881960084genus00000000000000000000000000000000000416.60000000000000000000000000000000000416.6000000000000000000000416.600000000000000000000000000000000000000000000000000000000416.600000888888758species00000000000000000000000000000000000416.60000000000000000000000000000000000416.6000000000000000000000000000000000000000000000000000000000000000000000000000000416.600000772094023genus000000000000000000247.700000000294.00000000000000000000000000000000000000000000000000000000000000000000000000000000000000000000000000000000000000000000000000000000772629322no\_rank000000000000000000247.700000000294.0000000000000000000000000000000000000000000000000000000000000000000000000000000000000000000000000000000000000000000000000000000077772030797species000000000000000000247.700000000294.0000000000000000000000000000000000000000000000000000000000000000000000000000000000000000000000000000000000000000000000000000000015848101515828211class00318.300000000000000000000000000000086.800000000379.500000000000000000000000000000000412.200000000000000000000400.000000000343.3000000000000000343.3000000000000000000000000000000000555555356order00230.0000000000000000000000000000000000000000000000000000000000000000000000000000000000000000000000000000000230.0000000000000000230.00000000000000000000000000000000004444444204455order00379.5000000000000000000000000000000000000000379.500000000000000000000000000000000379.500000000000000000000379.500000000379.5000000000000000379.500000000000000000000000000000000044444431989family00379.5000000000000000000000000000000000000000379.500000000000000000000000000000000379.500000000000000000000379.5000000000000000000000000379.500000000000000000000000000000000044444265genus00379.5000000000000000000000000000000000000000379.500000000000000000000000000000000379.5000000000000000000000000000000000000000000000379.5000000000000000000000000000000000444444147645species00379.5000000000000000000000000000000000000000379.5000000000000000000000000000000000000000000000000000000000000000000000000000000379.5000000000000000000000000000000000646666204457order00351.0000000000000000000000000000000000000000000000000000000000000000000000000445.000000000000000000000413.700000000413.7000000000000000413.7000000000000000000000000000000000646626241297family00351.0000000000000000000000000000000000000000000000000000000000000000000000000445.000000000000000000000413.7000000000000000000000000413.7000000000000000000000000000000000444444165695genus00445.0000000000000000000000000000000000000000000000000000000000000000000000000445.0000000000000000000000000000000000000000000000445.000000000000000000000000000000000014213238085243160324931522883145453494535341855126619103654170128624816615336816615332213122111161428216class001962.60001907.42819.32396.82518.43104.22057.302864.43663.302912.81816.22800.802965.52999.41393.83071.60475.002665.205054.42308.902173.4164.503610.702864.61527.80754.202383.900483.600524.01538.9002432.400000000000000000000000000524.0000000000000000000000000000000000000000002383.900483.600524.01538.9002432.4000000000000000000000002725185535812410151412825825361687280840order002579.20002745.52214.80000002840.20003923.9003002.8000003080.005054.42087.600003480.102864.600002383.90000000002449.6000000000000000000000000000000000000000000000000000000000000000000002383.90000000002449.600000000000000000000000849108258251506family002383.9000000000002840.900000000000000000000002718.500002383.90000000002449.6000000000000000000000000000000000000000000000000000000000000000000002383.90000000002449.600000000000000000000000848108258256222genus002383.9000000000002892.100000000000000000000002718.500002383.90000000002449.6000000000000000000000000000000000000000000000000000000000000000000002383.90000000002449.60000000000000000000000088888832002species002383.90000000000000000000000000000000000000002383.90000000000000000000000000000000000000000000000000000000000000000000000000000002383.90000000000000000000000000000000001710171085698species000000000000003615.900000000000000000000002718.5000000000000000000000000000000000000000000000000000000000000000000000000000000000000000000000000000000000000000000000151515151515217203species000000000000002148.700000000000000000000000000000000000002148.70000000000000000000000000000000000000000000000000000000000000000000000000000002148.7000000000000000000000001010102626865no\_rank000000000000002901.100000000000000000000000000000000000002901.10000000000000000000000000000000000000000000000000000000000000000000000000000002901.1000000000000000000000001010101010102282475species000000000000002901.100000000000000000000000000000000000002901.10000000000000000000000000000000000000000000000000000000000000000000000000000002901.10000000000000000000000011475682family003475.7000000000000000000000000004442.000000000000000000000000000000000000000000000000000000000000000000000000000000000000000000000000000000000000000000000000000000114303379genus003475.7000000000000000000000000004442.000000000000000000000000000000000000000000000000000000000000000000000000000000000000000000000000000000000000000000000000000000114114204773species003475.7000000000000000000000000004442.000000000000000000000000000000000000000000000000000000000000000000000000000000000000000000000000000000000000000000000000000000524573512468141211119060family001333.60002806.91864.300000000003923.9000000003080.005462.73517.500003480.1000000000000000000000000000000000000000000000000000000000000000000000000000000000000000000000000000000000000000000000004632008genus001645.0000000000000000000000000005462.700000000000000000000000000000000000000000000000000000000000000000000000000000000000000000000000000000000000000000000000000000464687882species\_group001645.0000000000000000000000000005462.70000000000000000000000000000000000000000000000000000000000000000000000000000000000000000000000000000000000000000000000000000024363512381447670genus0000002828.52153.700000000003923.9000000003103.8003517.500003480.10000000000000000000000000000000000000000000000000000000000000000000000000000000000000000000000000000000000000000000000024363512381424363512381447671species0000002828.52153.700000000003923.9000000003103.8003517.500003480.100000000000000000000000000000000000000000000000000000000000000000000000000000000000000000000000000000000000000000000000113107080042431600249215028281434431445252618451149521415686241661586166158206351order001848.80001714.02825.42308.22518.43106.92058.002902.43679.302948.01801.82675.602965.02999.31400.93077.40002561.2002467.002325.8003622.5001527.80754.20000483.600524.01538.9002378.500000000000000000000000000524.000000000000000000000000000000000000000000000483.600524.01538.9002378.5000000000000000000000001131070800424316002492150282814344314452526184511495214156862416615861661581371322134353562131481family001848.80001714.02825.42308.22518.43106.92058.002902.43679.302948.01801.82675.602965.02999.31400.93077.40002561.2002467.002325.8003622.5001527.80754.20000483.600524.01538.9002378.500000000000000000000000000524.000000000000000000000000000000000000000000000483.600524.01538.9002378.5000000000000000000000001121058786424215252363148282714037310449484171506458194149562316158161588861541110947147594214621152010741152482genus001856.70001719.92848.22308.22527.63142.82038.502896.43680.202991.31733.42697.502975.73031.11368.33095.00002585.6002339.002325.8003636.9001306.80751.10000483.60001538.9002378.500000000000000000000000000000000000000000000000000000000000000000000000483.60001538.9002378.500000000000000000000000175885237915303653175885237915303653484species0000001157.21671.2003303.5425.2003666.103352.80840.403154.22894.3000002161.90000000000000000000000000000000000000000000000000000000000000000000000000000000000000000000000000000000000000000000000000000000329196255883295625588485species000000420.30001718.31249.2000000360.301168.21145.500000441.000000000000000000000000000000000000000000000000000000000000000000000000000000000000000000000000000000000000000000000000000000001414528354no\_rank000000000001189.10000000000000000000000000000000000000000000000000000000000000000000000000000000000000000000000000000000000000000000000000000000000000000000000010513134251051313425486species0000001072.62089.2001718.2923.8002576.302416.200000000000000000000000000000000000000000000000000000000000000000000000000000000000000000000000000000000000000000000000000000000000000000050553833220535839537425205302487species0000001766.51820.802768.41935.91539.6001977.80001750.800003539.8000000000000000000000000000000000000000000000000000000000000000000000000000000000000000000000000000000000000000000000000000000000005491no\_rank00000001820.800000000000000000000000000000000000000000000000000000000000000000000000000000000000000000000000000000000000000000000000000000000000000000000000000055630588no\_rank00000001820.80000000000000000000000000000000000000000000000000000000000000000000000000000000000000000000000000000000000000000000000000000000000000000000000000004865699no\_rank0000003284.500000000000000003961.5000000000000000000000000000000000000000000000000000000000000000000000000000000000000000000000000000000000000000000000000000000000004848122587no\_rank0000003284.500000000000000003961.50000000000000000000000000000000000000000000000000000000000000000000000000000000000000000000000000000000000000000000000000000000000015135720no\_rank000000000000000000000003883.9000000000000000000000000000000000000000000000000000000000000000000000000000000000000000000000000000000000000000000000000000000000001010374833no\_rank000000000000000000000003848.40000000000000000000000000000000000000000000000000000000000000000000000000000000000000000000000000000000000000000000000000000000000055604162no\_rank000000000000000000000003954.80000000000000000000000000000000000000000000000000000000000000000000000000000000000000000000000000000000000000000000000000000000000077662598no\_rank0000001443.9000000000000000000000000000000000000000000000000000000000000000000000000000000000000000000000000000000000000000000000000000000000000000000000000000099935588no\_rank000000000000000000000003678.80000000000000000000000000000000000000000000000000000000000000000000000000000000000000000000000000000000000000000000000000000000000055935590no\_rank000000000000000000000004234.4000000000000000000000000000000000000000000000000000000000000000000000000000000000000000000000000000000000000000000000000000000000001919935599no\_rank000000000000000000000003721.2000000000000000000000000000000000000000000000000000000000000000000000000000000000000000000000000000000000000000000000000000000000009797942513no\_rank00000000001884.01472.300000000000000000000000000000000000000000000000000000000000000000000000000000000000000000000000000000000000000000000000000000000000000000000000814915911787204741442115693156621814915911787204741442115693156621488species001850.90001639.42891.102512.11692.02126.602617.5002471.31966.22978.000002316.00001594.100000002554.70000748.4000000000000000000000000000000000000000000000000000000000000000000000000000000000000000000000000000000000000000000888888489species000000000000002378.500000000000000000000000000000000000002378.50000000000000000000000000000000000000000000000000000000000000000000000000000002378.500000000000000000000000444444492species000000589.000000000000000000000000000000000000000589.0000000000000000000000000000000000000000000000000000000000000000000000000000000589.0000000000000000000000000000000151515151515493species000000000001538.900000000000000000000000000000000000001538.90000000000000000000000000000000000000000000000000000000000000000000000000000001538.900000000000000000000000000531225771227123516712113341219651227724121274495species0000001598.91765.6002069.41925.5000001370.52182.702251.02437.41528.100002131.1002540.900003992.3000000000000000000000000000000000000000000000000000000000000000000000000000000000000000000000000000000000000000000000001961251115539211251113988719subspecies0000001763.30002401.81302.4000000002674.41453.51569.300002148.400000003696.000000000000000000000000000000000000000000000000000000000000000000000000000000000000000000000000000000000000000000000000175145175145546263no\_rank0000001823.50002819.4000000000001607.900002148.400000000000000000000000000000000000000000000000000000000000000000000000000000000000000000000000000000000000000000000000000000002326915311559093244573803363092936519232691531155909324457380336309293651928449species001893.50002366.42950.3003393.71383.803118.83762.703357.202960.903128.43648.802133.90002778.000000003165.800000000000000000000000000000000000000000000000000000000000000000000000000000000000000000000000000000000000000000000000444444326522species000000501.200000000000000000000000000000000000000501.2000000000000000000000000000000000000000000000000000000000000000000000000000000501.2000000000000000000000000000000201220121853278species000000551.30000361.00000000000000000000000000000000000000000000000000000000000000000000000000000000000000000000000000000000000000000000000000000000000000000000000014366251406212623750no\_rank000000495.20001247.4345.00000000001819.81410.1737.7000000000000000000000000000000000000000000000000000000000000000000000000000000000000000000000000000000000000000000000000000000000003625140536251405655307species00000000001247.400000000001819.81410.1798.4000000000000000000000000000000000000000000000000000000000000000000000000000000000000000000000000000000000000000000000000000000000001261261853276species000000534.40000345.0000000000000000000000000000000000000000000000000000000000000000000000000000000000000000000000000000000000000000000000000000000000000000000000008888882666100species000000422.000000000000000000000000000000000000000422.0000000000000000000000000000000000000000000000000000000000000000000000000000000422.00000000000000000000000000000009766127737103161212538genus0000001166.22258.3002605.52419.6000002163.70002865.92036.200002464.500000003534.200000000000000000000000000000000000000000000000000000000000000000000000000000000000000000000000000000000000000000000000576511373610296576511373610296539species0000001369.02258.3002641.72456.3000002163.70002869.72036.200002419.100000003534.2000000000000000000000000000000000000000000000000000000000000000000000000000000000000000000000000000000000000000000000004124122528037species000000912.800002096.0000000000000000000000000000000000000000000000000000000000000000000000000000000000000000000000000000000000000000000000000000000000000000000000006666632257genus0000000000524.00000000000000000000000000000000000000524.0000000000000000000000000000000524.000000000000000000000000000000000000000000000000524.0000000000000000000000000000666666504species0000000000524.00000000000000000000000000000000000000524.0000000000000000000000000000000000000000000000000000000000000000000000000000000524.000000000000000000000000000019324611426121862522125513821218712461767372221450145373737391471121168525subphylum001527.30002722.62469.902746.72045.7002111.43420.702967.102803.702603.23038.42701.03041.402836.72122.52271.2002285.100274.302520.62221.102145.30000000000000000000000000003015.6000000000000000000000003434.9000000000003434.90000003434.90003426.200000000000000000000000000000003019.000000000039373737373939239228221class0000000000000000000000000000003265.6000000000000000000000000000000000003434.9000000000000000000000003434.9000000000003434.90000003434.90003426.200000000000000000000000000000003426.200000000037373737373737213118order0000000000000000000000000000003434.9000000000000000000000000000000000003434.9000000000000000000000003434.9000000000003434.90000003434.9000000000000000000000000000000000003434.9000000000373737373737213121family0000000000000000000000000000003434.9000000000000000000000000000000000003434.9000000000000000000000003434.9000000000003434.90000000000000000000000000000000000000000003434.90000000003737373737893genus0000000000000000000000000000003434.9000000000000000000000000000000000003434.9000000000000000000000003434.90000000000000000000000000000000000000000000000000000003434.90000000003737373737371986146species0000000000000000000000000000003434.9000000000000000000000000000000000003434.90000000000000000000000000000000000000000000000000000000000000000000000000000003434.90000000001932361142611186252212551382121871246156346222144910810829547class001527.30002730.62469.902746.72045.7002303.93420.702967.102803.702603.23038.42701.03041.402836.72122.52273.4002224.800303.702520.62221.102151.90000000000000000000000000002872.00000000000000000000000000000000000000000000000000000000000000000000000000000002872.0000000000193236114261118625221255138212187124615634622214491081081213849order001527.30002730.62469.902746.72045.7002303.93420.702967.102803.702603.23038.42701.03041.402836.72122.52273.4002224.800303.702520.62221.102151.90000000000000000000000000002872.00000000000000000000000000000000000000000000000000000000000000000000000000000002872.000000000019323611426111862522125513821218712461563452221449108108172294family001527.30002730.62469.902746.72045.7002303.93420.702967.102803.702603.23038.42701.03041.402836.72122.52273.4002224.800416.002520.62221.102151.90000000000000000000000000002872.00000000000000000000000000000000000000000000000000000000000000000000000000000002872.0000000000193236114261118625221255138212187124615634422214491081083222231204115219124712194genus001527.30002730.62469.902746.72045.7002303.93420.702967.102803.702603.23038.42701.03041.402836.72122.52273.4002224.800529.502520.62221.102151.90000000000000000000000000002872.00000000000000000000000000000000000000000000000000000000000000000000000000000002872.000000000059196species000000937.491.600000000000000000000000000000000000000000000000000000000000000000000000000000000000000000000000000000000000000000000000000000000000000000000000000059532020subspecies000000937.491.6000000000000000000000000000000000000000000000000000000000000000000000000000000000000000000000000000000000000000000000000000000000000000000000000000991273266no\_rank000000091.60000000000000000000000000000000000000000000000000000000000000000000000000000000000000000000000000000000000000000000000000000000000000000000000000001030240127917724191227114112012334115937285162226745713771365811262320692329199species001628.80002817.22976.102979.73223.9002327.93456.602957.803035.402641.23175.92719.63091.5002106.82475.9002898.600003556.1002093.60000000000000000000000000000000000000000000000000000000000000000000000000000000000000000000000000000000000000000000051401865120111149156175135675195140186512011114915617513567519360104no\_rank001193.60002857.12921.703124.30002474.43478.803425.002900.402528.02838.703096.60002429.7002933.600004024.2002236.500000000000000000000000000000000000000000000000000000000000000000000000000000000000000000000000000000000000000000000108108108108108108200species0000000000000000000000000000002872.0000000000000000000000000000000000002872.00000000000000000000000000000000000000000000000000000000000000000000000000000002872.00000000004106101554219327501041061015542193275010204species001389.500000001840.20001970.80002100.602567.02447.22327.500002176.2001758.100002546.9001665.900000000000000000000000000000000000000000000000000000000000000000000000000000000000000000000000000000000000000000000513107918512694628128139513107918512694628128139824species001434.40001743.82531.1001577.10000000002302.82496.43387.22559.902888.401609.6001920.900002284.32149.702815.600000000000000000000000000000000000000000000000000000000000000000000000000000000000000000000000000000000000000000000123725013311252768559758582323no\_rank00000000001032.60001191.10001162.201112.0858.2475.300001244.6001113.500003781.9001771.10000000000000000473.6000000000000003365.8000000000000000000000000000000000000000000000000000000000000000473.6000000000000003365.80000012372501331125276855975858111783234no\_rank00000000001032.60001191.10001162.201112.0858.2475.300001244.6001113.500003781.9001771.10000000000000000473.6000000000000003365.8000000000000000000000000000000000000000000000000000000000000000473.6000000000000003365.8000001237249133112527585597585812631401187137795818phylum00000000001032.60001191.10001166.401112.0858.2475.300001245.3001113.500003781.9001771.10000000000000000473.6000000000000003365.8000000000000000000000000000000000000000000000000000000000000000473.6000000000000003365.800000141613141613713059species0000000000000000000000000001470.9001167.000004435.9000000000000000000000000000000000000000000000000000000000000000000000000000000000000000000000000000000000000000000000001912121107181912121107181476577species000000000000001066.10001321.601544.00000001162.7001184.700000000000000000000000000000000000000000000000000000000000000000000000000000000000000000000000000000000000000000000000000002991124223958582979721895827no\_rank000000000000000000658.20651.20374.400001258.500961.600003728.00000000000000000000473.6000000000000003365.8000000000000000000000000000000000000000000000000000000000000000473.6000000000000003365.8000008888882699178species000000000000000000000000000000000003365.800000000000000000000000000000000003365.80000000000000000000000000000000000000000000000000000000000000000000000000000003365.8000005295292699179species000000000000000000952.000000000000000003765.5000000000000000000000000000000000000000000000000000000000000000000000000000000000000000000000000000000000000000000000005555552699180species000000000000000000473.6000000000000000000000000000000000000473.6000000000000000000000000000000000000000000000000000000000000000000000000000000473.6000000000000000000008568562699182species000000000000000000432.0000000002049.600749.7000000000000000000000000000000000000000000000000000000000000000000000000000000000000000000000000000000000000000000000000000094109941092699183species000000000000000000859.4000377.500001362.2001261.600000000000000000000000000000000000000000000000000000000000000000000000000000000000000000000000000000000000000000000000000006226222572088species0000000000000000000000819.700001571.1000000000000000000000000000000000000000000000000000000000000000000000000000000000000000000000000000000000000000000000000000000012686310116126863101162572089species000000000000001348.80001224.001107.70000001291.7001422.60000000000000000000000000000000000000000000000000000000000000000000000000000000000000000000000000000000000000000000000000000721892351532325810231611637351144653008826870933251411551411532066phylum002174.63393.403812.01632.22811.3002979.7002676.63401.203282.31785.73108.703064.32799.13143.43290.203482.42193.42477.5002903.100003926.3002637.50000001557.2000000000001908.61274.700000001950.00000000000000000000000000000000000000000000000000000000001557.2000000000001908.61274.700000001950.00000000007218923515323258102316116373511446530088268709332514115514115203490class002174.63393.403812.01632.22811.3002979.7002676.63401.203282.31785.73108.703064.32799.13143.43290.203482.42193.42477.5002903.100003926.3002637.50000001557.2000000000001908.61274.700000001950.00000000000000000000000000000000000000000000000000000000001557.2000000000001908.61274.700000001950.000000000072189235153232581023161163735114465300882687093325141155141152222143203491order002174.63393.403812.01632.22811.3002979.7002676.63401.203282.31785.73108.703064.32799.13143.43290.203482.42193.42477.5002903.100003926.3002637.50000001557.2000000000001908.61274.700000001950.00000000000000000000000000000000000000000000000000000000001557.2000000000001908.61274.700000001950.0000000000198123210382144199512927415247532555203492family0003450.403812.002190.8000002680.83763.6001808.53049.203096.53454.13344.7003542.102643.9002985.600004476.2002675.20000000000000000000000000001950.00000000000000000000000000000000000000000000000000000000000000000000000000000001950.0000000000198123210382144199512927415247532555133471247831215422848genus0003450.403812.002190.8000002680.83763.6001808.53049.203096.53454.13344.7003542.102643.9002985.600004476.2002675.20000000000000000000000000001950.00000000000000000000000000000000000000000000000000000000000000000000000000000001950.00000000001979104671226421728266333851232466723837184214851species0003450.404152.602347.8000002999.13442.9001690.73398.002880.43364.93384.6003568.003146.6003184.800004528.2002724.900000000000000000000000000000000000000000000000000000000000000000000000000000000000000000000000000000000000000000000543910976856subspecies0002918.40000000002781.3000000000003979.0000000000000000000000000000000000000000000000000000000000000000000000000000000000000000000000000000000000000000000000000000000000525525525283no\_rank0002918.40000000002716.8000000000000000000000000000000000000000000000000000000000000000000000000000000000000000000000000000000000000000000000000000000000000000000000881307428no\_rank00000000000002495.40000000000000000000000000000000000000000000000000000000000000000000000000000000000000000000000000000000000000000000000000000000000000000000002741491248274149124876857subspecies00000000000003143.50000003016.42887.4000003536.2001663.200004431.100000000000000000000000000000000000000000000000000000000000000000000000000000000000000000000000000000000000000000000000452971023115142317205451312811539674576859subspecies0003675.20002305.2000002999.70001875.43451.202886.43684.03414.2003521.702546.9003077.800003936.2000000000000000000000000000000000000000000000000000000000000000000000000000000000000000000000000000000000000000000000009584541695845416457405no\_rank00000000000003121.70001700.000003791.5003647.900002909.9000000000000000000000000000000000000000000000000000000000000000000000000000000000000000000000000000000000000000000000000000071157817115781469607no\_rank00000000000003361.200003990.30003292.1003481.50000000000000000000000000000000000000000000000000000000000000000000000000000000000000000000000000000000000000000000000000000000008736697814103698223897141174155615subspecies0003765.404152.600000002995.70000003088.703852.1003903.803112.4003520.900004790.3002858.100000000000000000000000000000000000000000000000000000000000000000000000000000000000000000000000000000000000000000000591195591195469602no\_rank000004139.600000003107.80000000000000000000004924.3003090.4000000000000000000000000000000000000000000000000000000000000000000000000000000000000000000000000000000000000000000005710957109469604no\_rank00000000000003552.8000000000004384.602839.8003376.3000000000000000000000000000000000000000000000000000000000000000000000000000000000000000000000000000000000000000000000000000032321307427no\_rank00000000000002993.700000000000000000000000000000000000000000000000000000000000000000000000000000000000000000000000000000000000000000000000000000000000000000000055555859species0000000000000000000000000000001950.0000000000000000000000000000000000001950.00000000000000000000000000000000000000000000000000000000000000000000000000000001950.000000000055143387subspecies0000000000000000000000000000001950.000000000000000000000000000000000000000000000000000000000000000000000000000000000000000000000000000000000000000000000000000001232201612322016860species00000000000002516.60000003545.60000003294.6000000000000000000000000000000000000000000000000000000000000000000000000000000000000000000000000000000000000000000000000000000077561541583098species00000000000002364.20001580.800000002673.701477.500000000000000000000000000000000000000000000000000000000000000000000000000000000000000000000000000000000000000000000000000000001141141307442no\_rank00000000000002090.50000000000000870.000000000000000000000000000000000000000000000000000000000000000000000000000000000000000000000000000000000000000000000000000000005456115456111307443no\_rank00000000000002295.70001580.800000002673.701698.50000000000000000000000000000000000000000000000000000000000000000000000000000000000000000000000000000000000000000000000000000000881307444no\_rank00000000000002693.500000000000000000000000000000000000000000000000000000000000000000000000000000000000000000000000000000000000000000000000000000000000000000000050819111293173508191112931732663009species00000000000002455.43958.00002671.703222.53411.9000002724.80000000000000000000000000000000000000000000000000000000000000000000000000000000000000000000000000000000000000000000000000000000590231522201011614383001156126851162340751411514111321129771family001921.20001580.43135.0002979.7002070.32712.603282.303151.603059.92687.83092.73471.302853.72171.82259.3002734.600003197.6002502.70000001557.2000000000001908.61274.7000000000000000000000000000000000000000000000000000000000000000001557.2000000000001908.61274.7000000000000000005902314222010113143830011561268511423407514115141152025361214956310544732067genus001921.20001580.43135.0003171.6002070.32712.603282.303202.103059.92687.83092.73471.302853.72171.82292.1002734.600003197.6002502.70000001557.2000000000001908.61274.7000000000000000000000000000000000000000000000000000000000000000001557.2000000000001908.61274.7000000000000000001291840542species000000000000000000002161.22033.4000002632.300000000000000000000000000000000000000000000000000000000000000000000000000000000000000000000000000000000000000000000000000000001291812918523794no\_rank000000000000000000002161.22033.4000002632.3000000000000000000000000000000000000000000000000000000000000000000000000000000000000000000000000000000000000000000000000000000033741145113374114511109328species000000898.0000000000002503.002122.21929.42502.000002315.0000000000000000000000000000000000000000000000000000000000000000000000000000000000000000000000000000000000000000000000000000000022154141334774349675371572215414133477434967537157157687species0000002258.33192.9000000004531.203470.903135.53183.23375.43501.602318.52356.42796.9002699.800004200.00000000000000000000000000000000000000000000000000000000000000000000000000000000000000000000000000000000000000000000000053262265326226157688species00000000002780.200000003081.202385.73146.1000001873.50000000000000000000000000000000000000000000000000000000000000000000000000000000000000000000000000000000000000000000000000000000141414141414157691species000000000000000000001908.60000000000000000000000000000000000001908.60000000000000000000000000000000000000000000000000000000000000000000000000000001908.6000000000000000000111111111111157692species0000000000000000000001274.70000000000000000000000000000000000001274.70000000000000000000000000000000000000000000000000000000000000000000000000000001274.7000000000000000001091623626171610916236261716554406species0000003613.00003389.100000003118.3003520.72974.93803.702880.60000000003236.800000000000000000000000000000000000000000000000000000000000000000000000000000000000000000000000000000000000000000000000561717321713537413551112633022no\_rank0000001557.23956.3000002002.82662.50003514.702055.72492.9000001752.5003419.000003258.40000000001557.20000000000000000000000000000000000000000000000000000000000000000000000000000001557.20000000000000000000000000000006161732111343141361617321113431413712357species00000003956.3000002046.82662.50003514.702357.52499.1000001830.2003419.000003258.4000000000000000000000000000000000000000000000000000000000000000000000000000000000000000000000000000000000000000000000006565712368species000000000000000000001502.30000001493.200000000000000000000000000000000000000000000000000000000000000000000000000000000000000000000000000000000000000000000000000000005555551785996species0000001557.2000000000000000000000000000000000000001557.20000000000000000000000000000000000000000000000000000000000000000000000000000001557.2000000000000000000000000000000383138383838383838203691phylum0000000000000000000000000000001724.9000000000000000000000000000000000001795.5000000000000000000000001782.5000000000001782.50000001782.50001782.50001782.50000000000000000000000000001782.50000000003831383838383838203692class0000000000000000000000000000001724.9000000000000000000000000000000000001795.5000000000000000000000001782.5000000000001782.50000001782.50001782.500000000000000000000000000000001782.500000000038313838383838136order0000000000000000000000000000001724.9000000000000000000000000000000000001795.5000000000000000000000001782.5000000000001782.50000001782.5000000000000000000000000000000000001782.5000000000383138383838137family0000000000000000000000000000001724.9000000000000000000000000000000000001795.5000000000000000000000001782.5000000000001782.50000000000000000000000000000000000000000001782.5000000000383138387387157genus0000000000000000000000000000001724.9000000000000000000000000000000000001795.5000000000000000000000001782.50000000000000000000000000000000000000000000000000000001782.50000000002121212121158species0000000000000000000000000000002042.9000000000000000000000000000000000002042.90000000000000000000000000000000000000000000000000000000000000000000000000000002042.90000000002121243275no\_rank0000000000000000000000000000002042.900000000000000000000000000000000000000000000000000000000000000000000000000000000000000000000000000000000000000000000000000001010102638727no\_rank0000000000000000000000000000001275.9000000000000000000000000000000000001275.90000000000000000000000000000000000000000000000000000000000000000000000000000001275.90000000001010101010101539298species0000000000000000000000000000001275.9000000000000000000000000000000000001275.90000000000000000000000000000000000000000000000000000000000000000000000000000001275.9000000000215710187349132412130312102372012249016028368231832492117598742173543321886673622765836116552155454536387645141783270no\_rank1881.501789.62153.82304.02894.92975.21982.4680.01599.31236.71228.602237.92823.94393.73395.12429.72145.801672.42914.01622.72133.701958.702374.43013.502253.42109.4593.0145.703608.803127.62192.902510.900000000606.40707.9002765.10002114.00000000612.00086.50000000000606.40224.70002114.0000612.0116.2000000231.0000612.0130.80000626.50130.80626.50130.80000000000000606.40505.7002765.10002114.00000626.500612.000102.200000021571018734913241213031210237201224901602836823183249211759874217354332188667362276583611655215545453638764514168336no\_rank1881.501789.62153.82304.02894.92975.21982.4680.01599.31236.71228.602237.92823.94393.73395.12429.72145.801672.42914.01622.72133.701958.702374.43013.502253.42109.4593.0145.703608.803127.62192.902510.900000000606.40707.9002765.10002114.00000000612.00086.50000000000606.40224.70002114.0000612.0116.2000000231.0000612.0130.80000626.50130.80626.50130.80000000000000606.40505.7002765.10002114.00000626.500612.000102.2000000215710187349132412130312102372012248916028368231832492117598742173543321886673622765836116552155454536387645142114121412112541976phylum1881.501789.62153.82304.02894.92975.21982.4680.01599.31236.71228.602237.92823.94393.73396.22429.72145.801672.42914.01622.72133.701958.702374.43013.502253.42109.4593.0145.703608.803127.62192.902510.900000000606.40707.9002765.10002114.00000000612.00086.50000000000606.40224.70002114.0000612.0116.2000000231.0000612.0130.80000626.50130.80626.50130.80000000000000606.40505.7002765.10002114.00000626.500612.000102.20000002362262721316511130192266978516541212286363368365369117743class001785.40003151.23055.70001146.802220.12445.603062.02716.02208.202168.43411.82330.82823.301959.201498.83013.502222.30074.303693.2002259.702402.700000000606.4000000000000000000086.50000000000606.40000000000116.2000000000000000000000000000000000000606.4000000000000000000086.300000023622627213165111301922669785165412122863633683653691200644order001785.40003151.23055.70001146.802220.12445.603062.02716.02208.202168.43411.82330.82823.301959.201498.83013.502222.30074.303693.2002259.702402.700000000606.4000000000000000000086.50000000000606.40000000000116.2000000000000000000000000000000000000606.4000000000000000000086.3000000236226272131651113019226697851654121218636336836536913111132735311049546family001785.40003151.23055.70001146.802220.12445.603062.02716.02208.202168.43411.82330.82823.301959.201498.83013.502222.30075.303693.2002259.702402.700000000606.4000000000000000000086.50000000000606.40000000000116.2000000000000000000000000000000000000606.4000000000000000000086.3000000236226192121541112916224627516241208636364516424755117871761016genus001785.40003151.23055.70001450.402397.82580.503767.52716.02276.302544.43440.42540.62919.40001520.53013.502239.700003693.2002259.702402.7000000000000000000000000000000000000000000000000000000000000000000000000000000000000000000000000000000000000000000843148281611114394895441323084314828161111439489544132301017species001502.90002955.53039.60001080.7001254.20001698.402687.33680.12691.52883.70001373.83013.502047.100003164.5002310.500000000000000000000000000000000000000000000000000000000000000000000000000000000000000000000000000000000000000000000268141018species000000000000000003185.33447.000000000000000004218.2000000000000000000000000000000000000000000000000000000000000000000000000000000000000000000000000000000000000000000000002681426814521097no\_rank000000000000000003185.33447.000000000000000004218.20000000000000000000000000000000000000000000000000000000000000000000000000000000000000000000000000000000000000000000000011135129111351291019species0000003330.200001266.100000000001741.400001288.8001694.000000000000000000000000000000000000000000000000000000000000000000000000000000000000000000000000000000000000000000000000000009244411754392444117543327575species001812.9000000002397.00000000003501.22628.700001760.9002629.30000000002402.70000000000000000000000000000000000000000000000000000000000000000000000000000000000000000000000000000000000000000004757857855194115337102521586422640652no\_rank0000004107.52745.30001698.002601.64555.2002667.50003214.302960.90001944.4001498.100003985.400000000000000000000000000000000000000000000000000000000000000000000000000000000000000000000000000000000000000000000000410861341086131316593species0000004107.500000000002846.00000000002022.5001566.000004471.800000000000000000000000000000000000000000000000000000000000000000000000000000000000000000000000000000000000000000000000155615561316596species000000000001272.80000000002746.800000000000003818.0000000000000000000000000000000000000000000000000000000000000000000000000000000000000000000000000000000000000000000000004781110124781110121705617species000000000000000002702.50003247.203199.30002826.000000003752.10000000000000000000000000000000000000000000000000000000000000000000000000000000000000000000000000000000000000000000000032637175326371752545799species000000000002034.503085.200000003386.4000001590.6002169.800000000000000000000000000000000000000000000000000000000000000000000000000000000000000000000000000000000000000000000000000008888834084genus00000000000326.10000000000000000000000000000000000000326.1000000000000000000000000000000326.100000000000000000000000000000000000000000000000326.10000000000000000000000000088888834085species00000000000326.10000000000000000000000000000000000000326.1000000000000000000000000000000000000000000000000000000000000000000000000000000326.1000000000000000000000000003164416159732genus00000000000222.800000000000000000000034.70000000000000000000000000000000000049.000000000000000000000000000000000000000000000000000000000000000000000000000000049.0000000155441512593645no\_rank00000000000229.200000000000000000000059.60000000000000000000000000000000000049.000000000000000000000000000000000000000000000000000000000000000000000000000000049.00000004444442594269species00000000000000000000000000000000049.00000000000000000000000000000000000049.000000000000000000000000000000000000000000000000000000000000000000000000000000049.0000000171717171759735genus00000000000915.50000000000000000000000000000000000000915.5000000000000000000000000000000915.500000000000000000000000000000000000000000000000915.5000000000000000000000000001717171717171585976species00000000000915.50000000000000000000000000000000000000915.5000000000000000000000000000000000000000000000000000000000000000000000000000000915.5000000000000000000000000005455151221065genus00000000000000000000000000000000084.800000000000000000000000000000000000124.0000000000000000000000116.200000000000000000000000000000000000000000000000000000000116.20000004444441218801species000000000000000000000000000000000124.000000000000000000000000000000000000124.0000000000000000000000000000000000000000000000000000000000000000000000000000000124.00000001111111111501783genus00000000000332.50000000000000000000000000000000000000332.5000000000000000000000000000000332.500000000000000000000000000000000000000000000000332.500000000000000000000000000111111111111237258species00000000000332.50000000000000000000000000000000000000332.5000000000000000000000000000000000000000000000000000000000000000000000000000000332.500000000000000000000000000555555117747class000000000000000000000000000000000130.8000000000000000000000000000000000000000000000000000000000000000000000130.8000000130.8000130.8000000000000000000000000000000000130.800000055555200666order000000000000000000000000000000000130.8000000000000000000000000000000000000000000000000000000000000000000000130.8000000130.80000000000000000000000000000000000000130.800000055555584566family000000000000000000000000000000000130.8000000000000000000000000000000000000000000000000000000000000000000000130.800000000000000000000000000000000000000000000130.800000019349177287106111283910067041124844925166229571791336881820525132830422765116521538765200643class1736.401792.51911.12216.22894.92937.21719.201734.41215.61827.202246.72832.54456.23396.81781.32154.401658.02875.91358.31729.202003.802559.6002255.200481.203553.803127.62112.702592.00000000000707.9002765.10002114.00000000612.0000000000000000224.70002114.0000612.00000000231.0000612.0000000000000000000000000000505.7002765.10002114.00000000612.00000000001934917728710611128391006704112484492516622957179133688182052513283042276511652153876521214325253531255541311591171549order1736.401792.51911.12216.22894.92937.21719.201734.41215.61827.202246.72832.54456.23396.81781.32154.401658.02875.91358.31729.202003.802559.6002255.200481.203553.803127.62112.702592.00000000000707.9002765.10002114.00000000612.0000000000000000224.70002114.0000612.00000000231.0000612.0000000000000000000000000000505.7002765.10002114.00000000612.0000000000135112412205757815family0000000000000942.22332.24456.22578.200000000001134.4001678.9000000000000000000001705.2002765.10000000000000000000000000000000000000000000000000000000000000000000000000001705.2002765.10000000000000000000001351124122057578512510816genus0000000000000942.22332.24456.22578.200000000001134.4001678.9000000000000000000001705.2002765.10000000000000000000000000000000000000000000000000000000000000000000000000001705.2002765.10000000000000000000001015215817species0000000000000004380.22097.900000000000000000000000000000000000000000000000000000000000000000000000000000000000000000000000000000000000000000000000000000000000000000088272559no\_rank0000000000000004334.5000000000000000000000000000000000000000000000000000000000000000000000000000000000000000000000000000000000000000000000000000000000000000000077777728113species00000000000000002765.100000000000000000000000000000000000002765.10000000000000000000000000000000000000000000000000000000000000000000000000000002765.100000000000000000000055555528116species00000000000001705.200000000000000000000000000000000000001705.20000000000000000000000000000000000000000000000000000000000000000000000000000001705.200000000000000000000000071071028119species0000000000000000000000000001623.9002476.6000000000000000000000000000000000000000000000000000000000000000000000000000000000000000000000000000000000000000000000000000011511111111171550family0000000000000210.50000000000000000000000000000000000000241.6000000000000000000000000000000224.7000000000000000224.70000000000000000000000000000000224.700000000000000000000000011511116116239759genus0000000000000210.50000000000000000000000000000000000000241.6000000000000000000000000000000224.700000000000000000000000000000000000000000000000224.70000000000000000000000005555552364787species0000000000000241.60000000000000000000000000000000000000241.6000000000000000000000000000000000000000000000000000000000000000000000000000000241.600000000000000000000000055985681712121171551family0000000573.200000601.6002580.01107.2000191.0000001464.5001908.400000000000000000000486.7000000000000000000000000000000000000000000000000000000000000000000000000000000486.7000000000000000000000000558856817121251641836genus0000000573.200000607.7002580.01107.2000191.0000001464.5001908.400000000000000000000486.7000000000000000000000000000000000000000000000000000000000000000000000000000000486.70000000000000000000000002584825848837species0000000000000768.5002580.000000000001895.0002628.50000000000000000000000000000000000000000000000000000000000000000000000000000000000000000000000000000000000000000000000000000205828123species0000000000000470.90001107.20000000000001300.0000000000000000000000000000000000000000000000000000000000000000000000000000000000000000000000000000000000000000000000000000020582058879243no\_rank0000000000000470.90001107.20000000000001300.0000000000000000000000000000000000000000000000000000000000000000000000000000000000000000000000000000000000000000000000000000077777736874species0000000000000381.40000000000000000000000000000000000000381.4000000000000000000000000000000000000000000000000000000000000000000000000000000381.4000000000000000000000000555555393921species0000000000000634.00000000000000000000000000000000000000634.0000000000000000000000000000000000000000000000000000000000000000000000000000000634.0000000000000000000000000173371672767191233785969623464123265729361651296770715821238253111171552family1703.901793.52123.72209.42894.92933.21488.001613.11185.61753.302497.32836.503364.51928.82196.501661.72887.21251.51677.002004.402588.7001910.400003519.403127.61926.403401.3000000000000000000000000000000000000000000000000000000000000000000000000000000000000000000000000000000000000000000173371672767191233785869623464123165729361641296770715821238253427114139612201032352143171361734473838genus1703.901793.52123.72209.42894.92933.21488.001613.11185.61753.302499.92836.503364.51928.82204.701661.72887.21249.71677.002004.402588.7001910.400003519.403127.61926.403401.300000000000000000000000000000000000000000000000000000000000000000000000000000000000000000000000000000000000000000015677121251174881362222611215117484461528129species00002088.82878.02902.31670.90002546.701983.01365.202040.701781.1002833.61969.1003174.501708.0002120.500004340.5000000000000000000000000000000000000000000000000000000000000000000000000000000000000000000000000000000000000000000000001566101149211715661011492117767031no\_rank00002088.802958.31587.30002607.401845.6000000002019.0002711.100002211.400004586.90000000000000000000000000000000000000000000000000000000000000000000000000000000000000000000000000000000000000000000000025118713614210211383301431053460995314725118113544210211383151341053442985314728131species0000003788.21426.7001176.81519.401383.41427.702810.72198.71431.301216.01519.41430.4235.201736.602039.9001696.900003067.002583.52067.10000000000000000000000000000000000000000000000000000000000000000000000000000000000000000000000000000000000000000000066246198no\_rank00000000001698.200000000000000000000000000000000000000000000000000000000000000000000000000000000000000000000000000000000000000000000000000000000000000000000000071591810715918101122984no\_rank00000000000001747.70000001767.31506.4000002263.2002821.700000000000000000000000000000000000000000000000000000000000000000000000000000000000000000000000000000000000000000000000000001683312442552190714017123575783752513550677331220435514481401711951532372967850628132species0000003147.81534.3001051.4002784.23109.303506.802479.402155.13003.01206.11742.70845.402765.5002170.500003587.8001805.30000000000000000000000000000000000000000000000000000000000000000000000000000000000000000000000000000000000000000000091238197459406460229579123819745940646022957553174no\_rank0000002953.50000002604.53071.903841.400003876.81533.81945.30002854.4002192.50000000000000000000000000000000000000000000000000000000000000000000000000000000000000000000000000000000000000000000000000000594615551211187172421045946155512111871724210428135species0000002164.20000002982.31155.601643.301574.001627.51608.30002927.701105.9002673.500004142.003671.80000000000000000000000000000000000000000000000000000000000000000000000000000000000000000000000000000000000000000000004452227species0000000000000000244.200000000001305.000000000000000000000000000000000000000000000000000000000000000000000000000000000000000000000000000000000000000000000000000000004444908937no\_rank0000000000000000244.200000000001305.00000000000000000000000000000000000000000000000000000000000000000000000000000000000000000000000000000000000000000000000000000000554195905541959076123species0000002398.40001141.6000001954.000003159.0000002059.6001843.600000000000000000000000000000000000000000000000000000000000000000000000000000000000000000000000000000000000000000000000000007101156711589436species00000000000001263.91049.401298.901782.001126.72884.1913.1000000000000000000000000000000000000000000000000000000000000000000000000000000000000000000000000000000000000000000000000000000000000710115671171011567111236517no\_rank00000000000001263.91049.401298.901782.001126.72884.1913.10000000000000000000000000000000000000000000000000000000000000000000000000000000000000000000000000000000000000000000000000000000000007132250128185115661286264657132250128185115661286264651177574species00000001403.1001520.5002287.02165.103202.202493.201781.93075.81352.41955.00002411.7002022.000002254.0002050.40000000000000000000000000000000000000000000000000000000000000000000000000000000000000000000000000000000000000000000013318248129227667145112638335no\_rank1767.801828.70001950.80000002322.12665.003129.91866.42247.702585.12637.8000002244.700000000000000000000000000000000000000000000000000000000000000000000000000000000000000000000000000000000000000000000000000000001331824812922766714511652716species1767.801828.70001950.80000002322.12665.003129.91866.42247.702585.12637.8000002244.7000000000000000000000000000000000000000000000000000000000000000000000000000000000000000000000000000000000000000000000000000000013318248129227667145111331824812922766714511575614no\_rank1767.801828.70001950.80000002322.12665.003129.91866.42247.702585.12637.8000002244.700000000000000000000000000000000000000000000000000000000000000000000000000000000000000000000000000000000000000000000000000000001010101010102005473family0000000000000238.0000000000000000000000000000000000000000000000000000000000000000000000000000000000000238.00000000000000000000000000000000238.0000000000000000000000000642005519family0000000000000172.7002565.500000000000000000000000000000000000000000000000000000000000000000000000000000000000000000000000000000000000000000000000000000000000000000064397864genus0000000000000172.7002565.500000000000000000000000000000000000000000000000000000000000000000000000000000000000000000000000000000000000000000000000000000000000000000064397865species0000000000000172.7002565.50000000000000000000000000000000000000000000000000000000000000000000000000000000000000000000000000000000000000000000000000000000000000000006464880074no\_rank0000000000000172.7002565.50000000000000000000000000000000000000000000000000000000000000000000000000000000000000000000000000000000000000000000000000000000000000000005555552005523family000000000000000000000000000000612.000000000000000000000000000000000000612.000000000000000000000000612.000000000000612.0000000000000000000000000000000000000000000612.000000000055555346096genus000000000000000000000000000000612.000000000000000000000000000000000000612.000000000000000000000000612.0000000000000000000000000000000000000000000000000000000612.000000000055555185300species000000000000000000000000000000612.000000000000000000000000000000000000612.0000000000000000000000000000000000000000000000000000000000000000000000000000000612.000000000055694427no\_rank000000000000000000000000000000612.0000000000000000000000000000000000000000000000000000000000000000000000000000000000000000000000000000000000000000000000000000072743977109884175666412005525family0000003492.02704.4000001353.6004533.202803.4002209.93613.100002702.9003662.000000003044.000000000000000000002114.00000000000000000000000000002114.0000000000000000000000000000000000000000000000000002114.0000000000000000007274397798741751125195950genus0000003492.02704.4000001353.6004533.202803.40003613.100002683.6003662.000000003044.0000000000000000000000000000000000000000000000000000000000000000000000000000000000000000000000000000000000000000000006169659183992553212828112species0000003686.30000002188.8004568.903383.60003613.100002965.3003668.800000000000000000000000000000000000000000000000000000000000000000000000000000000000000000000000000000000000000000000000000007247672476203275no\_rank00000000000002662.3004665.100000000000004160.0000000000000000000000000000000000000000000000000000000000000000000000000000000000000000000000000000000000000000000000000000043266156432661561307832no\_rank0000004089.50000000004736.4000003220.300002754.0003410.90000000000000000000000000000000000000000000000000000000000000000000000000000000000000000000000000000000000000000000000000000935103993510391307833no\_rank00000000000001820.4004635.200000000003166.4004427.30000000000000000000000000000000000000000000000000000000000000000000000000000000000000000000000000000000000000000000000000000272769182620507no\_rank00000002704.400000858.600000000000002610.1003511.300000000000000000000000000000000000000000000000000000000000000000000000000000000000000000000000000000000000000000000000000002727691827276918712710species00000002704.400000858.600000000000002610.1003511.3000000000000000000000000000000000000000000000000000000000000000000000000000000000000000000000000000000000000000000000000000066666375288genus0000000000000000000002114.00000000000000000000000000000000000002114.00000000000000000000000000002114.0000000000000000000000000000000000000000000000000002114.000000000000000000666666823species0000000000000000000002114.00000000000000000000000000000000000002114.00000000000000000000000000000000000000000000000000000000000000000000000000000002114.00000000000000000044444768503class000000000000000000000000000626.500000000000000000000000000000000000000000000000000000000000000000000000000000000626.5000626.500000000000000000000000000000626.5000000000000444444768507order000000000000000000000000000626.500000000000000000000000000000000000000000000000000000000000000000000000000000000626.5000000000000000000000000000000000626.50000000000007265243274611279132501330628715112669253338116640616229152637684090176899173470212677134192648919129192416872961440693260221362185792140812699137931955833412323651771549091037323235896115951056232083298110141315616663051534470645711470825114815961163056125931037323237896115951060232083699310146131259312181548242121125112211783272no\_rank2515.22656.62176.63140.62456.03830.22944.22842.33207.42900.32780.13320.22788.63362.33610.23657.13496.92535.03272.62491.02883.03162.23009.52977.13059.93364.12879.12569.54459.93958.82978.21909.23449.84063.23915.83948.32247.74833.82561.93525.54434.200000319.23358.3654.7000381.42942.71745.0383.72046.40967.21495.6486.22677.10528.91799.801288.82423.03099.700001501.900000563.0000261.2787.3107.001578.11170.8639.5509.5159.00003243.600001170.81429.6657.4159.03237.3001170.80657.4159.03189.600143.43368.81170.80143.43339.800000319.23358.3654.7000367.72942.71745.0383.72046.40967.21397.6486.22677.10569.91799.801207.32423.03099.7143.40001501.903339.87265072632609274132501149326418112588977197306180605192502358282727176279113198454461280125270188041804165228354405032382512821437951336118641372317532629123176206712785853738352411154510561953275148166301528706477082815911630125935373837241115451060195368214125931232152431124461611357103857153263225125939631239phylum2515.22690.02149.13146.92463.93830.23072.12838.73207.52890.63292.33345.12802.03484.63657.93674.73500.32533.83321.12582.12897.03156.73017.43004.03053.43389.12884.52562.54683.04176.73171.41940.73504.94068.44198.03941.42027.34834.32578.83533.84509.600000239.23358.31399.9000381.43510.01745.0300.92046.40967.21495.602808.20528.9001361.003099.700000000001399.9000261.20107.001578.10639.5535.800003243.6000001429.6003237.30000003189.60003368.80003339.800000239.23358.31399.9000367.73510.01745.0300.92046.40967.21397.602808.20569.9001281.103099.70000003339.8888833974no\_rank00000000001399.900000000000000000000000000000000000001399.90000000000000000000000000000001399.9000000000000000000000000000000000000000000000001399.9000000000000000000000000000888881930845genus00000000001399.900000000000000000000000000000000000001399.90000000000000000000000000000001399.9000000000000000000000000000000000000000000000001399.90000000000000000000000000008888881871025species00000000001399.900000000000000000000000000000000000001399.90000000000000000000000000000000000000000000000000000000000000000000000000000001399.9000000000000000000000000000725505247254716613239933523272111798923188366138590184092266682567174398052806754131099421670169621350964023438380781982612731329531267115841370917403327123105276712785353733524111381030188131810164570647082815911630537337241113810341881318101163042174214151767116264141147181121111630347191061class2514.22687.02146.23126.92406.73830.63098.22818.63209.02889.53309.73351.32809.23521.83681.93676.83501.12479.73362.72579.22908.63148.82998.13001.43043.93380.42921.52599.64693.94215.73439.61954.83515.34071.74198.03889.22047.74835.22588.63533.84525.000000239.23358.30000381.43510.01745.0243.800967.21217.102892.50282.0003538.004135.300000000000000261.200026.00321.2000003243.6000000003237.30000003189.60003368.8000000000239.23358.30000367.73510.01745.0243.800967.21076.902892.50282.0003538.004135.30000003368.83486371319921218183605244120288107718194129777841379271413236110787134812444264732104110120544726562817242411117164262411121237173258211186232866261451411511385order02777.52087.6978.003831.71419.62580.03249.72370.62460.81952.34442.53819.84464.13762.71871.52563.52447.12422.42416.33495.12667.02492.801301.52965.32181.94733.402042.21177.83024.8522.74108.21732.94140.04857.42618.43563.54559.100000000000240.03510.0056.40001799.500000000000000000000000261.200026.00000000000000000000000000000000000000000000231.43510.0056.40001461.7000000000000000032941319810618041022778407181820533099956921106822512444316912912053346561813624113882411311211290964family02784.52844.4003832.01967.21025.33286.30004442.53957.84561.93762.73114.60696.92589.8201.64127.13531.92218.202990.92977.71654.14733.40003172.9736.24342.03358.14587.64857.84413.53563.64570.800000000000184.03510.0000002307.400000000000000000000000170.100000000000000000000000000000000000000000000000170.13510.0000002307.400000000000000003294131981061804102267831718182051308994692110682231244431691291205334656181324113241131346744135342782916828731121279genus02784.52844.4003832.01967.21025.33286.30004442.53958.24567.03762.73114.60715.52589.8197.84127.13568.42218.202990.92977.71781.24733.40003172.9736.24342.03358.14587.64857.84413.53563.64570.80000000000003510.0000002307.40000000000000000000000000000000000000000000000000000000000000000000000003510.0000002307.4000000000000000020911319862410209780871505161957453131068015124413911812053326561813209113013624101547721553851619574531310680151136139118120283259287981280species03045.82935.2003832.02725.702631.300003959.94578.23764.83487.801188.22161.205272.05322.52847.504288.22978.22679.54733.40000902.54342.03642.84825.24857.84681.33563.64570.80000000000000000000000000000000000000000000000000000000000000000000000000000000000000000000000000000000000000000001854617161201082563315271117122341085121846170subspecies000004510.200000003916.83484.93685.80000000000004448.7000000004909.103933.44805.30000000000000000000000000000000000000000000000000000000000000000000000000000000000000000000000000000000000000000001212158878no\_rank0000000000000005374.60000000000000000000000000000000000000000000000000000000000000000000000000000000000000000000000000000000000000000000000000000000000000000000100100196620no\_rank000004933.20000000000000000000000000000000000000000000000000000000000000000000000000000000000000000000000000000000000000000000000000000000000000000000000000000048304830282459no\_rank000004397.500000004009.00000000000000000000000000000000000000000000000000000000000000000000000000000000000000000000000000000000000000000000000000000000000000000000002222359786no\_rank0000000000000005033.500000000000000000000000000000000000000000000000000000000000000000000000000000000000000000000000000000000000000000000000000000000000000000008367830no\_rank00000000000000000000000000000000000004314.500000000000000000000000000000000000000000000000000000000000000000000000000000000000000000000000000000000000000000000088451515no\_rank00000000000000000000000000000000000004314.500000000000000000000000000000000000000000000000000000000000000000000000000000000000000000000000000000000000000000000088426430no\_rank0000000000000004259.500000000000000000000000000000000000000000000000000000000000000000000000000000000000000000000000000000000000000000000000000000000000000000001414681288no\_rank0000000000000004829.40000000000000000000000000000000000000000000000000000000000000000000000000000000000000000000000000000000000000000000000000000000000000000000123123869816no\_rank00000000000000000000000000000000000005719.8005061.300000000000000000000000000000000000000000000000000000000000000000000000000000000000000000000000000000000000000000034513451889933no\_rank0000000000000003548.000000000000000000000000000000000000000000000000000000000000000000000000000000000000000000000000000000000000000000000000000000000000000000005125121074252no\_rank0000000000000000000000000000000000000003812.20000000000000000000000000000000000000000000000000000000000000000000000000000000000000000000000000000000000000000000537953791074919no\_rank00000000000003637.203856.7000000000000000000000000000000000000000000000000000000000000000000000000000000000000000000000000000000000000000000000000000000000000000000010101123523no\_rank000003939.20000000000000000000000000000000000000000000000000000000000000000000000000000000000000000000000000000000000000000000000000000000000000000000000000000041196216no\_rank00000000000000000000000000000000000000004577.0000000000000000000000000000000000000000000000000000000000000000000000000000000000000000000000000000000000000000000441193576no\_rank00000000000000000000000000000000000000004577.00000000000000000000000000000000000000000000000000000000000000000000000000000000000000000000000000000000000000000009709701323661no\_rank00000000000003855.65751.3000000000000000000000000000000000000000000000000000000000000000000000000000000000000000000000000000000000000000000000000000000000000000000001114913101952568361114913101952568361282species02316.40000003441.400003596.51633.7742.9002235.22675.5004087.40000000003379.4000000000000000000000000000000000000000000000000000000000000000000000000000000000000000000000000000000000000000000000000001441441441441441441286species0000000000000003203.300000000000000000000000000000000000003203.30000000000000000000000000000000000000000000000000000000000000000000000000000003203.300000000000000000000008888881290species00000000000000000000002158.50000000000000000000000000000000000002158.50000000000000000000000000000000000000000000000000000000000000000000000000000002158.5000000000000000038383838383828035species0000000000000003761.700000000000000000000000000000000000003761.70000000000000000000000000000000000000000000000000000000000000000000000000000003761.7000000000000000000000055555529380species00000000000000000000002545.60000000000000000000000000000000000002545.60000000000000000000000000000000000000000000000000000000000000000000000000000002545.60000000000000000595959595959985002species0000000000000004096.300000000000000000000000000000000000004096.30000000000000000000000000000000000000000000000000000000000000000000000000000004096.30000000000000000000000868828269965genus00000000000000128.50000000000000000000000000000000000000184.0000000000000000000000000000000170.100000000000000000000000000000000000000000000000170.1000000000000000000000006666661855823species00000000000000184.00000000000000000000000000000000000000184.0000000000000000000000000000000000000000000000000000000000000000000000000000000184.0000000000000000000000008775697779204121278794131148131142716211226695211186817family000000099.5000212.60138.3234.70431.8235.1118.60228.90392.4273.80469.70297.3001081.200539.40544.200002173.500000000000306.00056.4000149.000000000000000000000000352.200000000000000000000000000000000000000000000000306.00056.4000149.0000000000000000066347650714221214668455455466177621779672681386genus0000000101.300000144.7238.80532.6123.3127.40194.30479.3206.30247.20386.1001418.000667.30598.200002173.500000000000232.000-49.6000149.000000000000000000000000000000000000000000000000000000000000000000000000232.000-49.6000149.00000000000000000112078675454620386786661species\_group00000000000000176.2000180.1000145.1315.50446.00276.0000000000000000000000000232.0000000149.000000000000000000000000000000000000000000000000000000000000000000000000232.0000000149.000000000000000005555551428species00000000000000232.00000000000000000000000000000000000000232.0000000000000000000000000000000000000000000000000000000000000000000000000000000232.000000000000000000000000444444580165species0000000000000000000000149.0000000000000000000000000000000000000149.0000000000000000000000000000000000000000000000000000000000000000000000000000000149.00000000000000000955545185979no\_rank00000000000000000071.10000203.60000000000000000000000000000000-49.6000000000000000000000000000000000000000000000000000000000000000000000000000000-49.600000000000000000000555555486398species000000000000000000-49.6000000000000000000000000000000000000-49.6000000000000000000000000000000000000000000000000000000000000000000000000000000-49.6000000000000000000006441653685species\_group00000000000000817.30000000000000002070.00000000002173.50000000000000000000000000000000000000000000000000000000000000000000000000000000000000000000000000000000000000000005445441423species00000000000000938.80000000000000002070.00000000002173.50000000000000000000000000000000000000000000000000000000000000000000000000000000000000000000000000000000000000000008888829331genus00000000000000352.20000000000000000000000000000000000000352.2000000000000000000000000000000352.200000000000000000000000000000000000000000000000352.200000000000000000000000888881449species00000000000000352.20000000000000000000000000000000000000352.2000000000000000000000000000000000000000000000000000000000000000000000000000000352.20000000000000000000000088698758no\_rank00000000000000352.20000000000000000000000000000000000000000000000000000000000000000000000000000000000000000000000000000000000000000000000000000000000000000000081513866124400634genus00000000000000184.5000117.10000526.9000219.8000000000000000000000000000144.7000000000000000000000000000000000000000000000000000000000000000000000000000000144.70000000000000000000066666628031species000000000000000000144.7000000000000000000000000000000000000144.7000000000000000000000000000000000000000000000000000000000000000000000000000000144.700000000000000000000779877242636778no\_rank00000000000000173.7000104.60000170.4000219.8000000000000000000000000000000000000000000000000000000000000000000000000000000000000000000000000000000000000000000000000000000074742169540species00000000000000000000000155.4000235.50000000000000000000000000000000000000000000000000000000000000000000000000000000000000000000000000000000000000000000000000000000412157794444783794186818family000000144.00000000123.7000137.200078.60073.40158.700000161.50000000000000000000000000000000000000000000000000000026.00000000000000000000000000000000000000000000000000026.000000000000000004444441372genus000000000000000000000026.0000000000000000000000000000000000000000000000000000000000000000026.00000000000000000000000000000000000000000000000000026.0000000000000000057571649genus00000000000000158.4000197.7000000000000000000000000000000000000000000000000000000000000000000000000000000000000000000000000000000000000000000000000000000000000000065145186105551122186820family0000000164.700000151.2157.40144.00169.101147.700165.0000127.6000000000000000000000000135.6000000000000000000000000000000000000000000000000000000000000000000000000000000135.600000000000000000000000551351668555553186351637genus0000000177.600000151.2153.10144.00140.501147.700181.2000127.6000000000000000000000000135.6000000000000000000000000000000000000000000000000000000000000000000000000000000135.600000000000000000000000548554851639species00000000000000173.20168.5042.00000264.4000000000000000000000000000000000000000000000000000000000000000000000000000000000000000000000000000000000000000000000000000000000005555551642species00000000000000135.60000000000000000000000000000000000000135.6000000000000000000000000000000000000000000000000000000000000000000000000000000135.6000000000000000000000007661985891787184628260911353151719569912802851643313539002no\_rank002001.700002800.602404.702962.103059.02217.201603.02691.83040.81966.02561.93789.91878.53243.40822.01793.52257.5002722.51393.00001560.2002507.00000000000000000000000000000000000000000000000000000000000000000000000000000000000000000000000000000000000000000000076619758917871846282606113531517195699128028516433539738no\_rank002001.700002814.302404.702962.103059.02217.201603.02691.83055.41966.02561.93789.91878.53243.40822.01793.52257.5002722.51393.00001560.2002507.000000000000000000000000000000000000000000000000000000000000000000000000000000000000000000000000000000000000000000000766197589178718462826061135315171956991280285164339131126681033662232531378genus002001.700002814.302404.702962.103059.02217.201603.02691.83055.41966.02561.93789.91878.53243.40822.01793.52257.5002722.51393.00001560.2002507.00000000000000000000000000000000000000000000000000000000000000000000000000000000000000000000000000000000000000000000057211652381441691119946668187511401572116523814416911199466681875114011379species001888.000002792.002332.90001827.02270.30002255.41966.02236.801149.8733.0001686.72202.1001925.01393.00002035.0002520.600000000000000000000000000000000000000000000000000000000000000000000000000000000000000000000000000000000000000000000121511646276915174971215116462769151749729391species001166.600002347.1000003121.2665.7002708.13322.002597.40000001976.2003735.100000001714.3000000000000000000000000000000000000000000000000000000000000000000000000000000000000000000000000000000000000000000001736566574323329312101833135822173656657432332931210183313582284135species002464.800003001.303026.702493.503044.82373.20003486.203351.73677.72561.03381.9002122.32513.400000000002629.900000000000000000000000000000000000000000000000000000000000000000000000000000000000000000000000000000000000000000000671062624949no\_rank000000000000000000173.700000000854.7002642.800000000000000000000000000000000000000000000000000000000000000000000000000000000000000000000000000000000000000000000000000007107101785995species000000000000000000000000000854.7002642.800000000000000000000000000000000000000000000000000000000000000000000000000000000000000000000000000000000000000000000000000007214711607540165409307230501099688611883161005866366145421074217391507272735370106092163916807131756362337727284184602813291112601153713675739911725648035653653731111271013188131810570647082815953731111271013188131810815932522576614291125559915251192174132188281591077186826order2504.32680.52178.03154.72398.23449.63101.52821.33208.42893.43309.93357.22798.12961.83248.73103.53505.72435.13390.62580.02926.43148.33001.33015.13042.23385.82904.22630.92927.44215.73459.81959.53517.34079.64326.33912.3817.03793.72586.03189.03751.500000239.23358.30000689.801745.0320.200967.2455.402892.50282.0003538.004135.300000000000000000000321.2000003243.6000000003237.30000003189.60000000000000239.23358.30000689.801745.0320.200967.2455.402892.50282.0003538.004135.30000003189.67134691562507163409072193992885761872859925856279138907517158684582570153451047217802165261264262921601271921801024528771248115064573649912045325653251111211086187064708251111211086187082134181147082181300family2501.12687.62213.53273.82403.03449.63165.03062.73247.82979.63310.03385.82796.92995.33376.93008.03821.42649.53527.72582.82957.43434.93034.03119.63066.23414.42907.02678.43154.53977.23483.21946.93517.53544.14326.34121.01043.04181.22668.53189.03911.800000239.200000689.801745.0350.100967.2003193.800003538.000000000000000000000000000003243.6000000003237.300000000000000000000239.200000689.801745.0350.100967.2003193.800003538.0000000003237.371346915614991634089941929528857318660599258562751381475171584345825684522310466176681650912613629216012719217994245287612481150645736449120450156532511112110861870645111121108618706439279429197138197745861194923555888555175736271655687868018057563869057810189557123154473319912415358706470641301genus2501.12687.62214.93312.32403.03449.63185.13076.53247.82980.43319.93385.82796.92997.03391.03008.03826.62649.53529.82628.92958.63455.53035.63123.83066.23414.42907.02680.03154.53977.23484.41946.93517.53544.14326.34126.41043.04181.22677.03189.03911.800000239.200000689.801745.0350.100967.2003193.800003538.000000000000000000000000000003243.600000000000000000000000000000239.200000689.801745.0350.100967.2003193.800003538.0000000003243.695141310505152939986919643124391511043943587516154213551198951413105051529399869196431243855110439435875161541955511571302species3230.6971.61058.82890.5001748.03185.502291.33073.93660.002929.13545.72079.92831.903409.203017.02517.51951.83394.903533.71830.92692.0003697.31066.42279.6004212.2002764.2000000000000000000000000000000000000000000000000000000000000000000000000000000000000000000000000000000000000000000006184129390no\_rank0000000000000000004117.2000000000003861.300000002872.6000000000000000000000000000000000000000000000000000000000000000000000000000000000000000000000000000000000000000000006184161841467705no\_rank0000000000000000004117.2000000000003861.300000002872.600000000000000000000000000000000000000000000000000000000000000000000000000000000000000000000000000000000000000000000512410165822232515360722415861701218102011153013810277401056629114196724348336358285301114644892335116053367276718437432486755749730474681142310848186851303species2628.802339.30001233.13035.203003.01685.62809.202693.13407.42986.23917.82146.93144.61854.73021.62641.43289.13394.23032.03417.52393.92610.0002996.01591.40003963.1002819.0000000000000000000000000000000000000000000000000000000000000000000000000000000000000000000000000000000000000000000002868354675664319187516592162013883641364590171075194728683546756643191875165921620138836413645901710751947655813no\_rank2618.802302.100002394.203172.4901.42398.802631.93825.53163.84037.903061.42317.33015.72335.83416.33153.23007.63335.12781.42637.1003644.600004434.6002508.50000000000000000000000000000000000000000000000000000000000000000000000000000000000000000000000000000000000000000000091791471724632923831624894167743233653522199179147172463292383162489416774323365352219927666no\_rank2807.901977.400002662.103218.101598.303252.23321.03039.53870.702864.71504.12939.12735.73336.03005.803131.82143.52348.8003167.200003093.5002207.7000000000000000000000000000000000000000000000000000000000000000000000000000000000000000000000000000000000000000000001808565106967133559919831202115503067512242115131911808565106967133559919831202115503067512242115131911077464subspecies2765.802667.50001151.63121.003105.01921.63345.602943.83206.32762.33865.903024.502803.03146.93502.93038.73157.53463.62649.82750.0002924.400004438.9003032.900000000000000000000000000000000000000000000000000000000000000000000000000000000000000000000000000000000000000000000101699421619151743026141474814210961610169942161915174302614147481421096161458253subspecies2557.802165.800002846.701894.00002561.02100.23292.92933.403192.803233.72674.61847.72144.702467.41168.52953.3002848.100004169.3002634.90000000000000000000000000000000000000000000000000000000000000000000000000000000000000000000000000000000000000000000029524269317615349913117187922631169716912167672571717651538501752909564484864294692623163153483031150874864266671145106469325693116314815017482295284815641304species0000001762.23374.903236.52087.83308.42730.92371.33821.42246.04095.32670.64009.72382.43395.42333.32865.83527.73616.53341.22718.33144.3003550.62326.73841.13533.704534.804425.32221.404115.200000000000000000000000000000000000000000000000000000000000000000000000000000000000000000000000000000000000000000042713121139417045202410674155953437510774271312113941704520241067415595343751077347253no\_rank00000003692.103772.103000.1004285.403947.604110.62621.33390.82113.23211.83962.003243.32538.93281.600003822.4004575.5002117.40000000000000000000000000000000000000000000000000000000000000000000000000000000000000000000000000000000000000000000013407119264697239326253134071192646972393262531048332no\_rank00000004647.60000003751.504053.304417.104377.003196.4002657.41298.43039.500004383.8004874.7002232.10000000000000000000000000000000000000000000000000000000000000000000000000000000000000000000000000000000000000000000087456301172971012622233221212511065997135871088144222408507332578482442642219514121100554937753710895916249451305species002220.63935.6003158.93098.103241.62978.03227.302363.03886.404151.82256.03374.31789.83056.33961.52177.73400.702173.12624.53158.7002753.100004430.0001437.30000000000000000000000000000000000000000000000000000000000000000000000000000000000000000000000000000000000000000000014135269538623881515116065098483615951413526953862388151511606509848361595388919no\_rank002336.63698.2003451.83318.303335.12927.53136.4004021.604902.803551.01569.22955.63716.72456.73290.302163.403485.4002277.700004256.2001958.40000000000000000000000000000000000000000000000000000000000000000000000000000000000000000000000000000000000000000000054381127174267114161210543811221742671141112101307species00217.2000419.9264.501670.5333.6000195.501776.50202.101117.11738.5192.0905.001562.40252.0000000000000000000000000000000000000000000000000000000000000000000000000000000000000000000000000000000000000000000000000000000055568814no\_rank000000000000000000000001737.200000000000000000000000000000000000000000000000000000000000000000000000000000000000000000000000000000000000000000000000000000000000551214179no\_rank0000000000212.40000000000000000000000000000000000000000000000000000000000000000000000000000000000000000000000000000000000000000000000000000000000000000000000006251199192001192252683146820777216251199192193225148314512078211308species0000002803.7856.201385.41458.400790.01132.601396.501746.42264.21332.82417.52310.22177.501886.11942.91609.700001033.9000001934.4000000000000000000000000000000000000000000000000000000000000000000000000000000000000000000000000000000000000000000001212264199no\_rank0000000000000000000002296.3000000000000000000000000000000000000000000000000000000000000000000000000000000000000000000000000000000000000000000000000000000000000017926691792669322159no\_rank00000000000000001183.20579.60000000000000688.00000000000000000000000000000000000000000000000000000000000000000000000000000000000000000000000000000000000000000000000000017171051074no\_rank00000000000000000000000001919.2000000000000000000000000000000000000000000000000000000000000000000000000000000000000000000000000000000000000000000000000000000000964316121449254945011817696431612912236995618961309species000000000230.22294.33669.101131.500185.20622.6004396.53855.32951.804044.23025.42318.700002613.000000000000000000000000000000000000000000000000000000000000000000000000000000000000000000000000000000000000000000000000000137137210007no\_rank00000000000000000000000004449.300000000000000000000000000000000000000000000000000000000000000000000000000000000000000000000000000000000000000000000000000000000056105610511691no\_rank0000000000000000001044.0005330.03967.80000000000000000000000000000000000000000000000000000000000000000000000000000000000000000000000000000000000000000000000000000000000004484481155071no\_rank00000000000000000000004450.5004197.40000000000000000000000000000000000000000000000000000000000000000000000000000000000000000000000000000000000000000000000000000000004335584335581198676no\_rank00000000000000000000005851.5004031.302691.600000000000000000000000000000000000000000000000000000000000000000000000000000000000000000000000000000000000000000000000000000003153151437447no\_rank0000000000000000000004144.10004976.000000000000000000000000000000000000000000000000000000000000000000000000000000000000000000000000000000000000000000000000000000000065651310species000000000000000000811.000000000000000000003145.6000000000000000000000000000000000000000000000000000000000000000000000000000000000000000000000000000000000000000000008339675139121881816158339275139121881816151311species0000000001116.802967.10002290.03352.73365.9799.501904.23298.32612.32472.602180.901684.1000003599.700000000000000000000000000000000000000000000000000000000000000000000000000000000000000000000000000000000000000000000000004216466no\_rank00000000000000002856.500000000000000000000000000000000000000000000000000000000000000000000000000000000000000000000000000000000000000000000000000000000000000000044208435no\_rank00000000000000002856.500000000000000000000000000000000000000000000000000000000000000000000000000000000000000000000000000000000000000000000000000000000000000000012145433882557940438203143871504738210417110161103134268298018203912145426475347340438182142271284737710416581491037382829385741313species2254.602028.12430.800501.02549.202783.62911.12271.92079.22287.42650.32314.71932.31019.72263.21730.52480.22934.42911.71597.43009.92412.12465.71992.4002459.81811.90003179.704500.03145.104026.0000000000000000000000000000000000000000000000000000000000000000000000000000000000000000000000000000000000000000000726595671628726595671628189423no\_rank0000000003423.02464.200000350.002538.601083.202047.3768.00001833.600000002447.4000000000000000000000000000000000000000000000000000000000000000000000000000000000000000000000000000000000000000000000002281276522812765487214no\_rank00000003275.2001467.200000000000002736.2002968.700000003281.000003456.0000000000000000000000000000000000000000000000000000000000000000000000000000000000000000000000000000000000000000000445445488221no\_rank00000000002640.70002815.8000000000000000000000000000000000000000000000000000000000000000000000000000000000000000000000000000000000000000000000000000000000000000000006868488222no\_rank00000000002125.30003425.50000000000000000000000000000000000000000000000000000000000000000000000000000000000000000000000000000000000000000000000000000000000000000000041788684178868512566no\_rank00000002425.904148.300003505.10001579.100000003150.21127.2000000000000000000000000000000000000000000000000000000000000000000000000000000000000000000000000000000000000000000000000000000055516950no\_rank0000000000000000004086.800000000000000000000000000000000000000000000000000000000000000000000000000000000000000000000000000000000000000000000000000000000000000001313525381no\_rank000000000000000000000002126.20000000000000000000000000000000000000000000000000000000000000000000000000000000000000000000000000000000000000000000000000000000000044561276no\_rank0000000000000000000000000002418.00000000000000000000000000000000000000000000000000000000000000000000000000000000000000000000000000000000000000000000000000000000618146618146574093no\_rank00000002669.2000000000000000000000000000004500.03446.40000000000000000000000000000000000000000000000000000000000000000000000000000000000000000000000000000000000000000000058545854869309no\_rank0000000002158.91361.80000000000000000960.5000000000000000000000000000000000000000000000000000000000000000000000000000000000000000000000000000000000000000000000000000000066869311no\_rank00000000004400.30000000000000000000000000000000000000000000000000000000000000000000000000000000000000000000000000000000000000000000000000000000000000000000000001111869312no\_rank00000000000000002881.100000000000000000000000000000000000000000000000000000000000000000000000000000000000000000000000000000000000000000000000000000000000000000051158511581130804no\_rank00000002755.4003999.200000000000000001448.000000003140.600000000000000000000000000000000000000000000000000000000000000000000000000000000000000000000000000000000000000000000000981061510115114614521228703098106151011511461452122870301314species0000002986.7699.001169.0000951.33121.404925.602398.40591.63754.22996.82409.53444.42212.62972.42282.800000000000000000000000000000000000000000000000000000000000000000000000000000000000000000000000000000000000000000000000000000001632641211840494872366163652576101415819071668189983604326053346518597634726463354527344016712123521158125211468418723630713410135522424943031318species02604.72300.53251.02791.103468.83096.803066.23463.42722.13008.13210.13709.203838.203302.02532.43226.33701.92934.93279.03052.03058.12390.82792.63167.803390.91920.53268.7003952.303324.52789.703852.3000000000000000000000000000000000000000000000000000000000000000000000000000000000000000000000000000000000000000000631266597352122186848304048175545088710029733289121808893538119423027281532843631266597352122186848304048175545088710029733289121808893538119423027281532843760570no\_rank02755.02315.33231.53107.303463.93346.002993.43452.71871.73063.93265.73808.103942.003391.52758.83687.83607.83405.83257.303141.12249.93280.02899.803420.01860.33359.5003773.2002788.904004.7000000000000000000000000000000000000000000000000000000000000000000000000000000000000000000000000000000000000000000931265572962263518078049535508805443713854992479138614045956903012296167118793126557296226351807804953550880544371385499247913861404595690301229616711871114965no\_rank02556.72354.33322.42368.603348.12966.203141.63498.12773.72974.23252.83670.303796.603380.62343.73079.23939.52744.23431.73052.03001.22449.92599.3003403.41999.53174.2004846.2002951.8000000000000000000000000000000000000000000000000000000000000000000000000000000000000000000000000000000000000000000005555551326species000000000000001253.600000000000000000000000000000000000001253.60000000000000000000000000000000000000000000000000000000000000000000000000000001253.6000000000000000000000002520754898910951315588172342021621610101962082520754898910951315588172342021621610101962081335species00000002997.603687.12266.62962.4003337.804453.403802.12421.43252.002686.22066.803806.52863.53336.300003559.43444.405300.904579.61417.90000000000000000000000000000000000000000000000000000000000000000000000000000000000000000000000000000000000000000000015646156461340species00000000000002932.50000001660.30520.50000840.0000000000000000000000000000000000000000000000000000000000000000000000000000000000000000000000000000000000000000000000000000000017591015110132492176830321939231215175910151101324921768303219392312151343species00002660.2003565.20002176.102660.200003573.52050.82102.84148.12696.4003447.22252.22003.6003649.304228.5004835.2001562.4000000000000000000000000000000000000000000000000000000000000000000000000000000000000000000000000000000000000000000005555551346species0000000239.200000000000000000000000000000000000000239.2000000000000000000000000000000000000000000000000000000000000000000000000000000239.2000000000000000000000000000006666661349species00000000000000220.00000000000000000000000000000000000000220.0000000000000000000000000000000000000000000000000000000000000000000000000000000220.00000000000000000000000013873137703723700112784533021020213156396328081465289785279646408811440859532213823126333372751112552422161020641504725267213681831826344637156240242726328037species1944.101714.60002467.92940.402567.93274.21887.62869.92110.53353.32383.01398.32698.33090.62640.93032.13210.33112.22958.72569.62420.92246.92634.7003170.81930.90003893.9002618.900000000000000000000000000000000000000000000000000000000000000000000000000000000000000000000000000000000000000000000812204091314815143202224589659838266529059812204091314815143202224589659838266529059246201no\_rank00000002953.403277.73518.504085.82599.33341.2002662.03154.73960.95260.23226.43632.92910.502681.802931.9002995.500004088.4003148.600000000000000000000000000000000000000000000000000000000000000000000000000000000000000000000000000000000000000000000532515540107271639216112391052124107678532515540107271639216112391052124107678365659no\_rank001281.800002920.001896.13311.102482.61717.13296.5002886.52403.82310.92937.33370.32504.22124.02724.62440.602546.10002086.20004438.500000000000000000000000000000000000000000000000000000000000000000000000000000000000000000000000000000000000000000000000599566187752204518758128125913944563373385995661877522045187581281259139445633733833040species00002535.63681.8003566.03132.403797.903486.02932.43591.74552.504094.503565.43839.23533.93486.62985.93231.601849.00000003925.54695.5002951.13470.40000000000000000000000000000000000000000000000000000000000000000000000000000000000000000000000000000000000000000000162914913261743427754753983391115416711210384407111327206961344219845634species001687.60003019.02996.1002503.81716.302471.43056.002001.003251.52718.82864.73396.01300.03205.42601.4002814.6003381.11250.60004179.0002724.900000000000000000000000000000000000000000000000000000000000000000000000000000000000000000000000000000000000000000000825109615173023152741723081101658495825109615173023152741723081101658495889201no\_rank002158.60003240.33180.5003374.22006.302471.43560.30003440.52718.83259.43582.103241.22818.6002961.2003381.11155.60004424.5002800.5000000000000000000000000000000000000000000000000000000000000000000000000000000000000000000000000000000000000000000004195384894195384891302863no\_rank00000000000000002001.001626.4002999.31300.02944.2000000000003380.7000000000000000000000000000000000000000000000000000000000000000000000000000000000000000000000000000000000000000000000006752959310species002060.700000000000000000002241.100500.80000000000002679.90000000000000000000000000000000000000000000000000000000000000000000000000000000000000000000000000000000000000000000067529675291116231no\_rank002060.700000000000000000002241.100500.80000000000002679.9000000000000000000000000000000000000000000000000000000000000000000000000000000000000000000000000000000000000000000007152133683166102917115140874640543242320715213368316610291711514087464054324232078535species002220.30001995.91517.402072.23415.00002799.803766.903549.51303.93190.23276.42362.92491.001068.0848.82694.9002709.003007.7000001919.70000000000000000000000000000000000000000000000000000000000000000000000000000000000000000000000000000000000000000000071071082348species000000000000000000245.1000000338.700000000000000000000000000000000000000000000000000000000000000000000000000000000000000000000000000000000000000000000000000000000065214531373410051258144593063112114016574397108342211886726166521453137341005125814459306311211401657439710834221188672616113107species1004.32953.22304.93138.0002380.92843.302531.63622.42349.102086.03616.72665.93422.802699.42915.43111.22971.62482.42778.72486.42555.02166.52573.4002506.11940.52963.6003615.8003063.90000000000000000000000000000000000000000000000000000000000000000000000000000000000000000000000000000000000000000000041167215245926454672145119603species\_group0001857.20001676.3000001913.0873.10001296.803014.42904.22672.41560.402133.700001227.500000001146.800000000000000000000000000000000000000000000000000000000000000000000000000000000000000000000000000000000000000000000695195725645111711334species00000002500.700000000001027.603014.43414.62672.41912.302190.8000000000000000000000000000000000000000000000000000000000000000000000000000000000000000000000000000000000000000000000000000000000584245141119602subspecies0000000000000000001648.8002134.03242.0002179.0000000000000000000000000000000000000000000000000000000000000000000000000000000000000000000000000000000000000000000000000000000000723723617121no\_rank0000000000000000000002402.30002262.000000000000000000000000000000000000000000000000000000000000000000000000000000000000000000000000000000000000000000000000000000000051251336species0000000687.000000000001498.700964.400000000000000000000000000000000000000000000000000000000000000000000000000000000000000000000000000000000000000000000000000000000000005125512540041subspecies0000000687.000000000001498.700964.4000000000000000000000000000000000000000000000000000000000000000000000000000000000000000000000000000000000000000000000000000000000000032691019150055species0000000320.80000000000910.70002178.4898.40000000000000000000000000000000000000000000000000000000000000000000000000000000000000000000000000000000000000000000000000000000000031610316101076934no\_rank0000000327.90000000000910.70000898.400000000000000000000000000000000000000000000000000000000000000000000000000000000000000000000000000000000000000000000000000000000000555555197614species00000000000000003687.0000000105.600000000000772.8000000000000000000000000000000000000000000000000000000000000000000000000000000000000000000000000000000000000000000000003916539931926961063233113481589277757272289103670257758species1437.700000762.62475.303163.53006.41787.61935.01673.92482.20694.902031.72368.72455.81231.13406.81925.63174.5001882.40001423.00004088.300000000000000000000000000000000000000000000000000000000000000000000000000000000000000000000000000000000000000000000000391653993192696106323311348158927775727228910367039165399319269610632331134815892777572722891036701054460no\_rank1437.700000762.62475.303163.53006.41787.61935.01673.92482.20694.902031.72368.72455.81231.13406.81925.63174.5001882.40001423.00004088.30000000000000000000000000000000000000000000000000000000000000000000000000000000000000000000000000000000000000000000000021195759561252536315405species0000002543.01666.4000002084.0516.0000270.4002290.6000378.40000000000002149.00000000000000000000000000000000000000000000000000000000000000000000000000000000000000000000000000000000000000000000091956595553354subspecies0000002704.91666.4000000250.00000002211.5000378.40000000000000000000000000000000000000000000000000000000000000000000000000000000000000000000000000000000000000000000000000000000001010990317no\_rank00000001360.000000000000000000000000000000000000000000000000000000000000000000000000000000000000000000000000000000000000000000000000000000000000000000000000000096961258574no\_rank0000002704.9000000000000002211.50000000000000000000000000000000000000000000000000000000000000000000000000000000000000000000000000000000000000000000000000000000000000777777361101species000000000000000000282.3000000000000000000000000000000000000282.3000000000000000000000000000000000000000000000000000000000000000000000000000000282.3000000000000000000007575400065species00000001508.600000000000002243.200000000000000000000000000000000000000000000000000000000000000000000000000000000000000000000000000000000000000000000000000000000000003918244511119211215533574542471870170341225433881303711418562561833211412031315621132244143671232species\_group2862.90002351.43705.81597.22116.22907.62348.92213.73457.403047.52567.72886.12298.603412.93169.93425.43821.52530.63298.702296.32316.92775.5003786.83030.61336.53514.004808.004527.73036.02872.800000000000000000000000000000000000000000000000000000000000000000000000000000000000000000000000000000000000000000001623291110993463217475712351652874182255130164852353316192911109410027178396301564741312326321485212211328species00002425.43751.701802.92907.601540.93552.002913.12722.32578.62315.102050.23400.13288.93883.72809.83207.002406.72377.72740.5003790.402091.0004579.304739.63083.02357.60000000000000000000000000000000000000000000000000000000000000000000000000000000000000000000000000000000000000000000423399551072779423399551072779862970no\_rank000004188.500000003061.7002432.602433.002509.200003820.82733.93552.0004049.700000003266.200000000000000000000000000000000000000000000000000000000000000000000000000000000000000000000000000000000000000000000521859911295521859911295862971no\_rank000000000003136.602862.92586.00002350.3001911.42939.400000003797.3000000000000000000000000000000000000000000000000000000000000000000000000000000000000000000000000000000000000000000000000000056131697514121272910subspecies00000000000002010.4000003294.7003137.100002782.4004525.002279.6000002310.43131.20000000000000000000000000000000000000000000000000000000000000000000000000000000000000000000000000000000000000000000561316975141256131697514121353243no\_rank00000000000002010.4000003294.7003137.100002782.4004525.002279.6000002310.43131.2000000000000000000000000000000000000000000000000000000000000000000000000000000000000000000000000000000000000000000036415514756729281220103055133111764751444638201215102255586111338species3003.3000001597.2002418.41517.03122.103055.503568.22187.902440.903745.2002679.202003.62192.62905.40002383.6431.2004893.5002655.52177.400000000000000000000000000000000000000000000000000000000000000000000000000000000000000000000000000000000000000000008853388533591365no\_rank0000000002220.500000000003747.50000320.40000000004802.7000000000000000000000000000000000000000000000000000000000000000000000000000000000000000000000000000000000000000000000003141020631410206862966no\_rank00000000000003365.6002871.203262.400000000000000005160.20002828.000000000000000000000000000000000000000000000000000000000000000000000000000000000000000000000000000000000000000000003011822530118225862967no\_rank3176.8000000000000000001925.5000000003946.500000004542.0002664.800000000000000000000000000000000000000000000000000000000000000000000000000000000000000000000000000000000000000000000215811250158573415613076860species00000002632.2004342.22874.203568.102956.0004089.02412.02829.1003792.502694.90000000000002224.73097.200000000000000000000000000000000000000000000000000000000000000000000000000000000000000000000000000000000000000000002158112501585734156130511331674518184250subspecies00000002632.2004342.22874.203568.102956.0004089.02412.02829.1003792.502694.90000000000002224.73097.200000000000000000000000000000000000000000000000000000000000000000000000000000000000000000000000000000000000000000002114471171021144711710696216no\_rank00000002632.2000003548.602883.0005501.600004381.003229.400000000000000000000000000000000000000000000000000000000000000000000000000000000000000000000000000000000000000000000000000000000074960111127496011112862968no\_rank000000000002600.003982.600004213.100003929.30000000000000002951.000000000000000000000000000000000000000000000000000000000000000000000000000000000000000000000000000000000000000000003671512636715126862969no\_rank00000000000003777.200004127.22412.00003983.7000000000000002224.7000000000000000000000000000000000000000000000000000000000000000000000000000000000000000000000000000000000000000000007171717171711111760species00000000000000000000000003745.3000000000000000000000000000000000003745.30000000000000000000000000000000000000000000000000000000000000000000000000000003745.30000000000000038452823571043673116463308111750163199435573701726021621384528235710436731164633081117501631994355737017260216211433513species002489.100003182.502296.03494.002603.52691.43627.02737.31554.02822.23254.02680.23021.93305.72747.42413.00997.62226.72883.1003005.82024.20003992.2002241.5000000000000000000000000000000000000000000000000000000000000000000000000000000000000000000000000000000000000000000005555551825069species000000000000000000490.0000000000000000000000000000000000000490.0000000000000000000000000000000000000000000000000000000000000000000000000000000490.0000000000000000000006666661917441species0000000000000000000000000329.700000000000000000000000000000000000329.7000000000000000000000000000000000000000000000000000000000000000000000000000000329.7000000000000009999992021971species000000000000000000325.1000000000000000000000000000000000000325.1000000000000000000000000000000000000000000000000000000000000000000000000000000325.10000000000000000000088136212469121931461362675271571535881362124691219314613626752715715352382163species001938.42658.2002077.32844.50621.03534.72835.8003493.302280.403042.903049.13231.61291.42868.902051.803058.300000000002298.4000000000000000000000000000000000000000000000000000000000000000000000000000000000000000000000000000000000000000000009856142399856142392490633species00000000000003324.30000003180.60000773.30000001985.7000003168.52637.60000000000000000000000000000000000000000000000000000000000000000000000000000000000000000000000000000000000000000000442694902708637295140421748163633161113824685293518274384180525926227695839814741586683107931529523843801212985111091811109188697572513251362024520144746152238101665711012453122608887no\_rank2491.52801.22334.73505.72367.203114.83354.23888.02965.53461.32667.62827.12339.33429.33057.23829.32693.23135.72763.72777.43483.73218.83105.02902.03283.02591.12540.83990.603025.22001.73262.9003929.503330.02733.704255.600000000000001745.0000967.200753.000003538.00000000000000000000000000000000000000000000000000000000000000000001745.0000967.200753.000003538.0000000000473933512351525777257912113011139961527360112102617861481147818473933512351525777257912113011139961527360112102617861481147818712633species001909.90002869.62543.002942.51574.72159.302168.12684.92356.33423.62140.82460.41863.02456.42777.82860.42544.503099.91890.72222.6002392.800003327.0001836.90000000000000000000000000000000000000000000000000000000000000000000000000000000000000000000000000000000000000000000061418104415228523128284152429713071215614181044152285231282841524297130712151156431species2136.2000003081.32684.701521.03027.4002756.63803.703053.401668.403328.72825.92426.82557.502704.32543.43005.9002592.303161.4000002556.000000000000000000000000000000000000000000000000000000000000000000000000000000000000000000000000000000000000000000000255513445102241017721471801438923171501025551344510224101772147180143892317150101156433species0000002916.32809.802807.33647.5001421.34001.71790.82905.602593.72155.73957.23407.11721.42646.501995.42146.92942.400002867.7000001921.4000000000000000000000000000000000000000000000000000000000000000000000000000000000000000000000000000000000000000000006282227591028142499127911566141727264708136282227591028142499127911566141727264708131759399species001984.70002223.62986.5003683.62427.302902.83917.804140.503373.13024.13390.22455.12164.62231.501551.71859.52997.400002135.5002394.500000000000000000000000000000000000000000000000000000000000000000000000000000000000000000000000000000000000000000000000133216114531164791648731277713613341060016713321611453116479164873127771361334106001671839799species0000002492.43387.002859.32544.101485.504476.904361.604084.003950.92192.52412.92937.702823.82785.13075.3003344.01637.53677.1005294.6002449.10000000000000000000000000000000000000000000000000000000000000000000000000000000000000000000000000000000000000000000097771552256161316615529152869135977715522561613166155291528691351902136species2145.201772.600002587.501961.702013.001766.42629.92964.63923.902477.202333.92239.61956.91777.402284.001912.4002029.300003183.8000000000000000000000000000000000000000000000000000000000000000000000000000000000000000000000000000000000000000000000001010101010101940319species000000000000000000000967.2000000000000000000000000000000000000967.2000000000000000000000000000000000000000000000000000000000000000000000000000000967.2000000000000000009999992173853species0000000000000000000000000753.000000000000000000000000000000000000753.0000000000000000000000000000000000000000000000000000000000000000000000000000000753.0000000000000001111111111112420308species0000000000000000000000000000004037.2000000000000000000000000000000000004037.20000000000000000000000000000000000000000000000000000000000000000000000000000004037.20000000007777772420309species0000000000000000000000000000002753.6000000000000000000000000000000000002753.60000000000000000000000000000000000000000000000000000000000000000000000000000002753.6000000000211853441744714134110931761380712721238815541421185344174471413411093176138071272123881554142576376species2828.302623.2000812.02612.902440.8803.402938.02296.93352.53170.94164.503222.61494.93204.81527.43083.02797.003525.02450.92157.100000003950.4000000000000000000000000000000000000000000000000000000000000000000000000000000000000000000000000000000000000000000000001111111111112583585species00000000000000001745.000000000000000000000000000000000000001745.00000000000000000000000000000000000000000000000000000000000000000000000000000001745.000000000000000000000044438847149110611374286372425296444388471491106113742863724252962598453species00572.000001977.101593.92908.701543.71531.41883.201297.001026.82768.31921.92500.13168.11518.40002030.100000002783.9002385.7000000000000000000000000000000000000000000000000000000000000000000000000000000000000000000000000000000000000000000002633992638135984209480343763481402209305391519101619398546643722476373717449490824927317102409123052633992638135984209480343763481402209305391519101619398546643722476373717449490824927317102409123052610896species02808.72426.53553.52381.003136.13469.13888.03151.93382.13082.72945.43025.53783.003852.53219.33569.82786.03166.73593.53435.43471.43105.83325.62628.22906.33879.403096.92026.63137.4004235.903521.62772.604255.60000000000000000000000000000000000000000000000000000000000000000000000000000000000000000000000000000000000000000001951952684658species000000000000000000468.1000188.40000000000000000000000000000000000000000000000000000000000000000000000000000000000000000000000000000000000000000000000000000000000008781046876251712261339291651723719151357genus000871.600858.3503.500614.9000825.20471.40363.5608.6824.0710.2910.61285.1000912.90000000164.800975.2000000000000000000000000000000000000000000000000000000000000000000000000000000000000000000000000000000000000000000008781026873251012251332915171341511358species000871.600858.3510.100614.9000852.70471.40550.9608.6938.0710.201285.1000956.40000000000975.200000000000000000000000000000000000000000000000000000000000000000000000000000000000000000000000000000000000000000000878101687025121133291498564442552110310529721359subspecies000871.600858.3514.300614.9000880.50471.400612.70710.201285.10001019.00000000000844.1000000000000000000000000000000000000000000000000000000000000000000000000000000000000000000000000000000000000000000002257261541828772257261541828771104322no\_rank000000641.3599.200642.5000695.30408.000542.40658.1000001058.30000000000987.00000000000000000000000000000000000000000000000000000000000000000000000000000000000000000000000000000000000000000000068611360subspecies000000000000000000160.700000000000000000001122.80000000000000000000000000000000000000000000000000000000000000000000000000000000000000000000000000000000000000000000077684738no\_rank00000000000000000000000000000000000000884.30000000000000000000000000000000000000000000000000000000000000000000000000000000000000000000000000000000000000000000051182616109687752536309897682013112594061618243071753511043736497410373649741011521714113233958family2548.401794.52919.0002127.31939.23208.304329.33139.60461.6670.63314.41721.1249.1496.62451.163.24382.72721.11540.71040.03247.62826.4171.3002639.703491.71006.403651.7002611.902469.00000003358.30000000215.70001133.502675.40274.000004135.30000000000000000000000000000000000000000000000000000000003358.30000000215.70001133.502675.40274.000004135.3000000045451253genus000000255.000000000000170.8000000000000000000000000000000000000000000000000000000000000000000000000000000000000000000000000000000000000000000000000000000000000000051182215109687751343098863201211259366161723277175331104373649741037364974105812631473410838101261520636623275131941578genus2548.401794.52919.0002467.72055.53208.304329.33164.000704.83314.41228.80522.42451.165.24382.72721.11600.91040.03249.52943.9127.0002639.703491.71006.403876.1002611.902469.00000003358.30000000215.70001133.502675.40274.000004135.30000000000000000000000000000000000000000000000000000000003358.30000000215.70001133.502675.40274.000004135.300000006306301580species00000000917.80000003556.800000000000000000000000000000000000000000000000000000000000000000000000000000000000000000000000000000000000000000000000000000000000000000007777771587species00000000000000000000000001601.1000000000000000000000000000000000001601.10000000000000000000000000000000000000000000000000000000000000000000000000000001601.10000000000000012254122541588species000000002553.100000000000000002936.40000000000000000000000000000000000000000000000000000000000000000000000000000000000000000000000000000000000000000000000000000000002091122031121590species000000002965.600000000000000002668.800000000000000000000000000000000000000000000000000000000000000000000000000000000000000000000000000000000000000000000000000000000066337330subspecies000000002905.0000000000000000000000000000000000000000000000000000000000000000000000000000000000000000000000000000000000000000000000000000000000000000000000000005723187711471596species000000002794.0003482.600000000003340.4003922.50000000000003936.000000000000000000000000000000000000000000000000000000000000000000000000000000000000000000000000000000000000000000000472218347221831380360no\_rank000000002552.5003482.600000000003438.7003951.300000000000000000000000000000000000000000000000000000000000000000000000000000000000000000000000000000000000000000000000000000000067671598species0000000000000003251.5000000000451.40000000000000000000000000000000000000000000000000000000000000000000000000000000000000000000000000000000000000000000000000000000004444441601species00000000000000000000001133.50000000000000000000000000000000000001133.50000000000000000000000000000000000000000000000000000000000000000000000000000001133.5000000000000000020151316072106513130721613species000000003092.400000000003135.2003467.4003010.70000000000002564.400000000000000000000000000000000000000000000000000000000000000000000000000000000000000000000000000000000000000000000953095301381124no\_rank000000003127.100000000000000003372.80000000000000000000000000000000000000000000000000000000000000000000000000000000000000000000000000000000000000000000000000000000004444441622species000000000000000000000000000274.000000000000000000000000000000000000274.0000000000000000000000000000000000000000000000000000000000000000000000000000000274.0000000000000592110771637332710121624species2548.40000002118.12706.202985.43457.9000000130.000001807.603849.20000000000000000000000000000000000000000000000000000000000000000000000000000000000000000000000000000000000000000000000000000000005610763756107637362948no\_rank2548.40000002613.8002985.43457.9000000000002526.203219.800000000000000000000000000000000000000000000000000000000000000000000000000000000000000000000000000000000000000000000000000000000019324193241194971no\_rank000000002705.400000000000000003939.00000000000000000000000000000000000000000000000000000000000000000000000000000000000000000000000000000000000000000000000000000000006141202056141202051633species000000000002177.70000000003375.30002282.50000000005336.2002114.40000000000000000000000000000000000000000000000000000000000000000000000000000000000000000000000000000000000000000000066666628038species000000000000000000215.7000000000000000000000000000000000000215.7000000000000000000000000000000000000000000000000000000000000000000000000000000215.700000000000000000000545433959species0000000000000000000005264.83504.500000000000000000000000000000000000000000000000000000000000000000000000000000000000000000000000000000000000000000000000000000000000015151515151535787species00000000000000000000000001061.5000000000000000000000000000000000001061.50000000000000000000000000000000000000000000000000000000000000000000000000000001061.50000000000000055928565319117559250559701747715species000000002796.9003536.20003494.90003349.204799.20004132.03360.5000000000000000000000000000000000000000000000000000000000000000000000000000000000000000000000000000000000000000000000000000000006767568703no\_rank0000000000000003979.1000000000000000000000000000000000000000000000000000000000000000000000000000000000000000000000000000000000000000000000000000000000000000000015111511568704no\_rank0000000000000004034.5000004009.700000000000000000000000000000000000000000000000000000000000000000000000000000000000000000000000000000000000000000000000000000000000002052051088720no\_rank0000000000000003678.7000005203.400000000000000000000000000000000000000000000000000000000000000000000000000000000000000000000000000000000000000000000000000000000000006956951316933no\_rank0000000000000004207.00000000003478.800000000000000000000000000000000000000000000000000000000000000000000000000000000000000000000000000000000000000000000000000000000056561318634no\_rank0000000000000004318.8000005303.0000000000000000000000000000000000000000000000000000000000000000000000000000000000000000000000000000000000000000000000000000000000000017516175161380361no\_rank0000000000000003339.60000000004487.100000000000000000000000000000000000000000000000000000000000000000000000000000000000000000000000000000000000000000000000000000000064646464646447770species00000000000000000000000003358.4000000000000000000000000000000000003358.40000000000000000000000000000000000000000000000000000000000000000000000000000003358.4000000000000006666697478species00000000000000000000000001254.3000000000000000000000000000000000001254.30000000000000000000000000000000000000000000000000000000000000000000000000000001254.300000000000000661130798no\_rank00000000000000000000000001254.3000000000000000000000000000000000000000000000000000000000000000000000000000000000000000000000000000000000000000000000000000000000555555109790species00000000000000000000000001984.4000000000000000000000000000000000001984.40000000000000000000000000000000000000000000000000000000000000000000000000000001984.400000000000000107107107107107107152331species000000003262.6000000000000000000000000000000000000003262.60000000000000000000000000000000000000000000000000000000000000000000000000000003262.60000000000000000000000000000171717171717227942species000000002627.7000000000000000000000000000000000000002627.70000000000000000000000000000000000000000000000000000000000000000000000000000002627.70000000000000000000000000000515151515151304207species000000002932.2000000000000000000000000000000000000002932.20000000000000000000000000000000000000000000000000000000000000000000000000000002932.2000000000000000000000000000061010253631519675617457198198533024655183species\_group002364.70004521.003225.404573.63730.2000677.0002698.3004592.90003323.800003651.200000003239.7000000003582.50000000000000000000000000000000000000000000000000000000000000000000000000000003582.5000000000000000000000000000019819819821981981582species000000003573.4000000000000000000000000000000000000003582.50000000000000000000000000000000000000000000000000000000000000000000000000000003582.500000000000000000000000000001961961215914no\_rank000000003582.6000000000000000000000000000000000000000000000000000000000000000000000000000000000000000000000000000000000000000000000000000000000000000000000000006597256315194756170576589906315194756156571597species002364.70004882.003221.804573.63730.2000677.8002698.3004592.90003310.000003651.200000003239.700000000000000000000000000000000000000000000000000000000000000000000000000000000000000000000000000000000000000000000662142461447714subspecies000000003417.700000000000000003981.4000000000000000000000000000000000000000000000000000000000000000000000000000000000000000000000000000000000000000000000000000000000262262537973no\_rank000000003592.2000000000000000000000000000000000000000000000000000000000000000000000000000000000000000000000000000000000000000000000000000000000000000000000000001541541226298no\_rank000000002938.6000000000000000000000000000000000000000000000000000000000000000000000000000000000000000000000000000000000000000000000000000000000000000000000000002828321967no\_rank000000002794.40000000000000000000000000000000000000000000000000000000000000000000000000000000000000000000000000000000000000000000000000000000000000000000000000045451446494no\_rank000000002732.900000000000000000000000000000000000000000000000000000000000000000000000000000000000000000000000000000000000000000000000000000000000000000000000000101010110102107999species000000000000000000000000000000003908.3000000000000000000000000000000000004135.30000000000000000000000000000000000000000000000000000000000000000000000000000004135.3000000099525326no\_rank000000000000000000000000000000004160.60000000000000000000000000000000000000000000000000000000000000000000000000000000000000000000000000000000000000000000000000056562620435no\_rank000000000000000000116.00000002222.000000000000000000000000000000000000000000000000000000000000000000000000000000000000000000000000000000000000000000000000000000000025111793560270263348343127147529115963371612635610161276724121413623237136148959543313282211328661065281852family00710.5505.400410.41556.50218.8178.0491.90515.4409.23608.1190.4503.31352.50514.11744.7614.0374.50969.71961.4827.104442.91311.83269.23414.14083.001671.803451.8225.000000000000000000000154.001920.40000000000000000000000000000000000000000000000000000000000000000000000000000154.001920.4000000000000002161623525264262028239127146715987403716802988755215718121413623213136146454526141912018261810351410141614316221761350genus00794.3630.700433.81568.20218.1178.0546.00520.8410.53608.1190.0591.21507.90604.81744.7690.9349.401058.42159.1924.404442.91701.53269.23414.14083.001805.503451.8224.800000000000000000000210.501920.40000000000000000000000000000000000000000000000000000000000000000000000000000210.501920.4000000000000007965651261554413266653178101313623137965651261553813266651178101313404131351species001463.40001470.8379.70406.40000397.43602.61102.7516.8369.50782.60527.4573.50580.00505.904442.93554.63889.63669.04083.00505.8000000000000000000000000000000000000000000000000000000000000000000000000000000000000000000000000000000000000000000000003434226185no\_rank0000000000000000000000000000000004056.3000000000000000000000000000000000000000000000000000000000000000000000000000000000000000000000000000000000000000000000000066699186no\_rank000000000000000000461.0000000000000000000000000000000000000000000000000000000000000000000000000000000000000000000000000000000000000000000000000000000000000000020102201021201292no\_rank000000000000000000000000000474.6000003507.9000000000000000000000000000000000000000000000000000000000000000000000000000000000000000000000000000000000000000000000000065651206105no\_rank0000000000000000000000000000000003141.5000000000000000000000000000000000000000000000000000000000000000000000000000000000000000000000000000000000000000000000000018181287066no\_rank0000000000000000000000000000000005016.30000000000000000000000000000000000000000000000000000000000000000000000000000000000000000000000000000000000000000000000000812334922582660144079053706281726352111818113614081233492258266014407905370628172635211181811361401352species00196.8000282.71577.90214.2178.0000426.40180.701758.501251.21748.91080.4223.801200.52277.61272.700187.800002018.003451.8212.40000000000000000000000000000000000000000000000000000000000000000000000000000000000000000000000000000000000000000000081081033945species00000000000000781.0000528.8000000000000000000000000000000000000000000000000000000000000000000000000000000000000000000000000000000000000000000000000000000000000000044444437734species0000000000000000000000210.5000000000000000000000000000000000000210.5000000000000000000000000000000000000000000000000000000000000000000000000000000210.5000000000000000041681040688830471241681040688830471244008species00828.2000955.1556.500000626.3543.1000719.40497.80600.8566.6000698.40000000716.5000000000000000000000000000000000000000000000000000000000000000000000000000000000000000000000000000000000000000000000001275127553345species00000000000000172.2000111.1000070.80000000000000000000000000000000000000000000000000000000000000000000000000000000000000000000000000000000000000000000000000000000000011411411411453346species00000000000000112.90126.50195.100000000110.00000000000000000000000000000000000000000000000000000000000000000000000000000000000000000000000000000000000000000000000000000000172296172296417368species00000000000000416.2000474.70000531.9000628.700000000000000000000000000000000000000000000000000000000000000000000000000000000000000000000000000000000000000000000000000000005552608891no\_rank00000000000000000000000001920.4000000000000000000000000000000000001920.40000000000000000000000000000000000000000000000000000000000000000000000000000001920.4000000000000005555552582830species00000000000000000000000001920.4000000000000000000000000000000000001920.40000000000000000000000000000000000000000000000000000000000000000000000000000001920.400000000000000514325101890612154203850851191535532511122737genus000355.000193.6397.60182.80451.20552.2426.60283.0428.3474.50360.00554.7502.40510.10425.00000000561.600000000000000000000000000000000000000000000000000000000000000000000000000000000000000000000000000000000000000000000000612967612967519472species0000000169.3000000223.8000166.00000250.7000393.100000000000000000000000000000000000000000000000000000000000000000000000000000000000000000000000000000000000000000000000000000005814758147633807species0000000000000123.2332.2000250.300000000360.000000000000000000000000000000000000000000000000000000000000000000000000000000000000000000000000000000000000000000000000000000004923108706913118334373617413312331242648499no\_rank000394.800163.6439.3000451.201011.8472.10283.0463.6519.70355.60584.4528.70549.00444.60000000581.10000000000000000000000000000000000000000000000000000000000000000000000000000000000000000000000000000000000000000000000065307212821653072128212562451species0000000522.7000001260.8474.8000579.6000660.6244.5000486.500000000000000000000000000000000000000000000000000000000000000000000000000000000000000000000000000000000000000000000000000000008177405756152134515138177405756152134515132571750species000000141.2409.8000514.900470.10315.2428.9433.90367.20540.8597.40695.80385.80000000595.8000000000000000000000000000000000000000000000000000000000000000000000000000000000000000000000000000000000000000000000006765567151668genus0000000000000070.7000136.3000124.0000000000000000000000000000000000000108.8000000000000000000000000000000000000000000000000000000000000000000000000000000108.8000000000000000055555551669species0000000000000000000000108.8000000000000000000000000000000000000108.8000000000000000000000000000000000000000000000000000000000000000000000000000000108.800000000000000008136193561343114186827family000000974.2676.200000430.7384.7000818.0000466.7576.5000244.5000000000000000000000000000000000000000000000000000000000000000000000000000000000000000000000000000000000000000000000000000000012519186132589661375genus0000000723.500000483.2384.7000308.6000466.7576.5000000000000000000000000000000000000000000000000000000000000000000000000000000000000000000000000000000000000000000000000000000000001091091377species0000000634.80000000000152.9000000000000000000000000000000000000000000000000000000000000000000000000000000000000000000000000000000000000000000000000000000000000000011711751665species00000000000000264.700000000174.9000000000000000000000000000000000000000000000000000000000000000000000000000000000000000000000000000000000000000000000000000000000005172689587genus0000001322.0000000000001357.500000000000000000000000000000000000000000000000000000000000000000000000000000000000000000000000000000000000000000000000000000000000000005175172036206species0000001322.0000000000001357.5000000000000000000000000000000000000000000000000000000000000000000000000000000000000000000000000000000000000000000000000000000000000000041217559172422528112555275993851235211959411334852044155122186828family00325.01037.800685.8580.901195.10470.20658.4664.00670.9369.8573.50705.10602.2578.50657.6632.4713.30000000622.100470.4000000000000000000000000285.60000000000000000000000000321.20000000000000000000000000000000000000000000000000000285.6000000000000483214201071369519314518522894443367156453162492747genus000669.200797.8589.5000510.10745.6543.70444.5461.0370.00683.90456.5302.50628.30471.00000000635.700440.7000000000000000000000000241.0000000000000000000000000000000000000000000000000000000000000000000000000000000241.0000000000000598135585114161612211959813558511416161221192748species0000001188.0706.6000706.201001.6762.10490.20514.60858.20663.7453.10742.90702.00000000823.50000000000000000000000000000000000000000000000000000000000000000000000000000000000000000000000000000000000000000000000012291648912124892751species0000000548.0000000327.2000210.9000261.0197.8000405.6000000000000000000000000000000000000000000000000000000000000000000000000000000000000000000000000000000000000000000000000000000017124171241234679no\_rank00000000000000253.9000207.0000261.0000000000000000000000000000000000000000000000000000000000000000000000000000000000000000000000000000000000000000000000000000000000000881086147709species0000000682.8000000155.2000193.60000393.5000176.000000000000000000000000000000000000000000000000000000000000000000000000000000000000000000000000000000000000000000000000000000008810868810861266845subspecies0000000682.8000000155.2000193.60000393.5000176.000000000000000000000000000000000000000000000000000000000000000000000000000000000000000000000000000000000000000000000000000000001414812145448981255257487no\_rank00000000000000308.7000179.6000258.5118.8000219.40000000179.2000000000000000000000000000241.0000000000000000000000000000000000000000000000000000000000000000000000000000000241.0000000000000655655208596species00000000000000278.7000291.600000000182.000000000000000000000000000000000000000000000000000000000000000000000000000000000000000000000000000000000000000000000000000000004444442592353species000000000000000000000000000241.000000000000000000000000000000000000241.0000000000000000000000000000000000000000000000000000000000000000000000000000000241.000000000000055555191769genus000000000000000000000000000321.200000000000000000000000000000000000321.20000000000000000000000000321.20000000000000000000000000000000000000000000000000000321.20000000000005552632303no\_rank000000000000000000000000000321.200000000000000000000000000000000000321.2000000000000000000000000000000000000000000000000000000000000000000000000000000321.20000000000005555551911586species000000000000000000000000000321.200000000000000000000000000000000000321.2000000000000000000000000000000000000000000000000000000000000000000000000000000321.20000000000008822611015140294053156522112131121470540genus0001222.100641.4580.301430.00000794.10867.20666.60727.30717.1817.50775.50943.60000000635.8000000000000000000000000000000000000000000000000000000000000000000000000000000000000000000000000000000000000000000000005121051210708126species00000000000000816.8000575.80000788.9000000000000000000000000000000000000000000000000000000000000000000000000000000000000000000000000000000000000000000000000000000000007722610513128283742146322163142621505no\_rank0001015.900718.1580.301430.00000793.00963.20675.10751.20757.4838.70814.60955.80000000635.800000000000000000000000000000000000000000000000000000000000000000000000000000000000000000000000000000000000000000000000761691911125333113581876169191112533311358181903686species0001015.900852.2592.2000000882.801077.60744.30803.50816.4983.10851.60995.00000000708.000000000000000000000000000000000000000000000000000000000000000000000000000000000000000000000000000000000000000000000000614417411561441741152496265species0000000548.7000000209.40706.00223.6000271.0431.9000500.200000000000000000000000000000000000000000000000000000000000000000000000000000000000000000000000000000000000000000000000000000001362405331603215373648086682261432281203582713515441011751911411186801class0000003270.21843.5002664.21363.602028.32275.21479.21459.402664.72425.02894.52305.32155.82212.60669.6902.41511.7001280.60518.4138.501673.0001847.8000000000000000002046.4000000842.700960.80510.500000000000000000000798.6619.900000000001429.60000000000000000000000000000000002046.4000000854.200619.90510.5000000013624053316032153726480866822614322812025827135154410117519114362341524786561161186802order0000003270.21843.5002664.21363.602028.32275.21479.21459.402671.62425.02894.52305.32155.82212.60669.6902.41511.7001286.50518.4138.501673.0001847.8000000000000000002046.4000000842.700960.80510.500000000000000000000798.6619.900000000001429.60000000000000000000000000000000002046.4000000854.200619.90510.5000000010491051114474434151331979family0000000000000136.400102.00198.20510.00131.80000144.700245.60510.5143.40000000000000000000000000000000000510.5000000000000000000000000000000000000000000000000000000000000000000000000000000510.50000000785106474478510671485genus0000000000000118.00000197.80130.4000000147.600162.70510.5143.40000000000000000000000000000000000510.5000000000000000000000000000000000000000000000000000000000000000000000000000000510.500000004444441491species00000000000000000000000000000000510.500000000000000000000000000000000000510.5000000000000000000000000000000000000000000000000000000000000000000000000000000510.50000000551649459genus00000000000000000000889.6000000000383.0000000000000000000000000000000000000000000000000000000000000000000000000000000000000000000000000000000000000000000000000000055154046species00000000000000000000889.6000000000383.000000000000000000000000000000000000000000000000000000000000000000000000000000000000000000000000000000000000000000000000000005555742737no\_rank00000000000000000000889.6000000000383.0000000000000000000000000000000000000000000000000000000000000000000000000000000000000000000000000000000000000000000000000000030115523132694687482113426183522175114645124113511413683810577186803family0000000775.7001340.41363.601107.81319.71479.21686.501359.92425.01473.01566.31031.51486.5001047.51152.9001077.300001850.0001233.1000000000000000002046.4000000538.900960.80000000000000000000000212.4960.8000000000000000000000000000000000000000000002046.4000000518.500960.800000000044444841genus000000000000000000000000000000960.800000000000000000000000000000000000960.800000000000000000000000960.8000000000000000000000000000000000000000000000000000000960.800000000044444166486species000000000000000000000000000000960.800000000000000000000000000000000000960.8000000000000000000000000000000000000000000000000000000000000000000000000000000960.800000000044536231no\_rank000000000000000000000000000000960.800000000000000000000000000000000000000000000000000000000000000000000000000000000000000000000000000000000000000000000000000008135326568271356561451871186928no\_rank00000000001764.8001874.91737.61479.21686.502027.62425.01707.50000001168.5001661.800000000000000000000000002046.4000000824.7000000000000000000000000000000000000000000000000000000000000000000000002046.4000000824.700000000000079326514127932651412712991species00000000001963.1002595.801479.21686.502027.6000000001643.8001791.900000000000000000000000000000000000000000000000000000000000000000000000000000000000000000000000000000000000000000000000000005555551898203species00000000000000000002046.40000000000000000000000000000000000002046.40000000000000000000000000000000000000000000000000000000000000000000000000000002046.400000000000000000006666662109691species000000000000000000000000000824.700000000000000000000000000000000000824.7000000000000000000000000000000000000000000000000000000000000000000000000000000824.70000000000006566161207244genus000000000000000000000000000294.300000000000000000000000000000000000196.00000000000000000000000000212.40000000000000000000000000000000000000000000000000000212.4000000000000555555649756species000000000000000000000000000196.000000000000000000000000000000000000196.0000000000000000000000000000000000000000000000000000000000000000000000000000000196.0000000000000562572511genus000000000000001180.4000000000000235.3000000000000000000000000000000000000000000000000000000000000000000000000000000000000000000000000000000000000000000000000000000054542648079no\_rank000000000000001180.4000000000000143.00000000000000000000000000000000000000000000000000000000000000000000000000000000000000000000000000000000000000000000000000000000192817767340181262612321141164882genus00000001036.000000978.11493.60001419.001536.61785.11140.61509.4001047.51350.4001240.300001662.6000000000000000000000000000000000000000000000000000000000000000000000000000000000000000000000000000000000000000000000001928177673401812626123211419281776734018126261232114617123species00000001036.000000978.11493.60001419.001536.61785.11140.61509.4001047.51350.4001240.300001662.600000000000000000000000000000000000000000000000000000000000000000000000000000000000000000000000000000000000000000000000119731506553genus000000000000000000000000000390.400385.200000000000000000000000000000000000000000000000000000000000000000000000000000000000000000000000000000000000000000000000000004646208479species000000000000000000000000000265.500439.200000000000000000000000000000000000000000000000000000000000000000000000000000000000000000000000000000000000000000000000000008132812529161151465135186804family00000003378.5002841.1001506.500002103.203030.403584.100002375.2001435.500000003042.4000000000000000000000000000000000000000000000000000000000000000000000000000000000000000000000000000000000000000000007912844259genus00000002961.7000002420.000000000000002869.4001117.0000000000000000000000000000000000000000000000000000000000000000000000000000000000000000000000000000000000000000000000000000079128143361species00000002961.7000002420.000000000000002869.4001117.000000000000000000000000000000000000000000000000000000000000000000000000000000000000000000000000000000000000000000000000000007912879128546269no\_rank00000002961.7000002420.000000000000002869.4001117.00000000000000000000000000000000000000000000000000000000000000000000000000000000000000000000000000000000000000000000000000000131562841870884genus00000000002841.1001271.700002246.70003700.10000892.5000000000000000000000000000000000000000000000000000000000000000000000000000000000000000000000000000000000000000000000000000000013156284131562841496species00000000002841.1001271.700002246.70003700.10000892.5000000000000000000000000000000000000000000000000000000000000000000000000000000000000000000000000000000000000000000000000000000010181423825425237634138565911977538999no\_rank0000003462.63464.4003867.9002391.33810.801484.503310.403295.93564.71564.03403.601090.702759.2001510.600000000000000000000000000000000000000000000000000000000000425.1000000000000000000000000000000000000000000000000000000425.1000000000101814238254252376341385659119771121261543314family0000003462.63464.4003867.9002391.33810.801484.503310.403295.93564.71564.03403.601090.702759.2001510.600000000000000000000000000000000000000000000000000000000000425.1000000000000000000000000000000000000000000000000000000425.10000000001017141731641821663283366986331genus0000003462.63571.8003867.9002663.74035.601484.503433.603426.43734.103476.30003154.5001477.000000000000000000000000000000000000000000000000000000000000000000000000000000000000000000000000000000000000000000000000000001017141731641821663283366910171417316418216632833669114527species0000003462.63571.8003867.9002663.74035.601484.503433.603426.43734.103476.30003154.5001477.0000000000000000000000000000000000000000000000000000000000000000000000000000000000000000000000000000000000000000000000000000065969210122342143393species00000000000001666.43411.10003030.703192.701540.200002140.6001778.800000000000000000000000000000000000000000000000000000000000000000000000000000000000000000000000000000000000000000000000000006596921012234265969210122342888727no\_rank00000000000001666.43411.10003030.703192.701540.200002140.6001778.80000000000000000000000000000000000000000000000000000000000000000000000000000000000000000000000000000000000000000000000000000777772060094genus000000000000000000000000000000425.100000000000000000000000000000000000000000000000000000000000425.1000000000000000000000000000000000000000000000000000000425.1000000000772623050no\_rank000000000000000000000000000000425.10000000000000000000000000000000000000000000000000000000000000000000000000000000000000000000000000000000000000000000000000000744773732304686family0000000000000000000000000001098.3000000000000000000000000000000000001678.000000000000000000000000001678.0000000000001429.600000000000000000000000000000000000000001429.6000000000000444441637257genus0000000000000000000000000001678.0000000000000000000000000000000000001678.000000000000000000000000001678.000000000000000000000000000000000000000000000000000001678.000000000000044444884684species0000000000000000000000000001678.0000000000000000000000000000000000001678.00000000000000000000000000000000000000000000000000000000000000000000000000000001678.000000000000044699246no\_rank0000000000000000000000000001678.00000000000000000000000000000000000000000000000000000000000000000000000000000000000000000000000000000000000000000000000000000000851911201141761711417526524class0000000000000924.8124.8000402.300000000162.200469.8000000000000000000000000278.10000000155.000481.4000000000000000000107.00000481.40000000000000000000000000000000000000000000278.10000000155.000481.4000000000851911201141761711417526525order0000000000000924.8124.8000402.300000000162.200469.8000000000000000000000000278.10000000155.000481.4000000000000000000107.00000481.40000000000000000000000000000000000000000000278.10000000155.000481.40000000008519112011417617114178533128827family0000000000000924.8124.8000402.300000000162.200469.8000000000000000000000000278.10000000155.000481.4000000000000000000107.00000481.40000000000000000000000000000000000000000000278.10000000155.000481.40000000007455241647genus000000000000000000395.300000000208.5000000000000000000000000000483.4000000000000000000000000000000000000000000000000000000000000000000000000000000483.4000000000000000000005555551648species000000000000000000483.4000000000000000000000000000000000000483.4000000000000000000000000000000000000000000000000000000000000000000000000000000483.40000000000000000000066666191303genus000000000000000000107.0000000000000000000000000000000000000107.000000000000000000000000000000107.0000000000000000000000000000000000000000000000000107.0000000000000000000006662638206no\_rank000000000000000000107.0000000000000000000000000000000000000107.0000000000000000000000000000000000000000000000000000000000000000000000000000000107.0000000000000000000006666661712675species000000000000000000107.0000000000000000000000000000000000000107.0000000000000000000000000000000000000000000000000000000000000000000000000000000107.0000000000000000000006444433334no\_rank000000000000000000705.700000000155.000000000000000000000000000000000000155.0000000000000000000000000000000000000000000000000000000000000000000000000000000155.000000000000064446544447no\_rank000000000000000000705.700000000155.000000000000000000000000000000000000155.0000000000000000000000000000000000000000000000000000000000000000000000000000000155.00000000000004444442109692species000000000000000000000000000155.000000000000000000000000000000000000155.0000000000000000000000000000000000000000000000000000000000000000000000000000000155.0000000000000444441918538genus000000000000000000000000000000300.000000000000000000000000000000000000300.000000000000000000000000300.0000000000000000000000000000000000000000000000000000000300.00000000004442636055no\_rank000000000000000000000000000000300.000000000000000000000000000000000000300.0000000000000000000000000000000000000000000000000000000000000000000000000000000300.00000000004444442584943species000000000000000000000000000000300.000000000000000000000000000000000000300.0000000000000000000000000000000000000000000000000000000000000000000000000000000300.000000000013131313132679910genus000000000000000000000000000000537.200000000000000000000000000000000000537.200000000000000000000000537.2000000000000000000000000000000000000000000000000000000537.20000000001313131313132487118species000000000000000000000000000000537.200000000000000000000000000000000000537.2000000000000000000000000000000000000000000000000000000000000000000000000000000537.2000000000157541082145307377508343415224843169339626124235011632428294873238836595554442691252591255736573614121909932class002202.53399.52551.702957.23014.82998.73043.02940.72788.82518.72950.33153.303643.703076.33179.92786.63233.13239.23055.84005.73439.52323.52459.13212.202590.91487.13130.1004159.404096.42540.701877.200000000000000785.000000542.90000732.30000000000000000000000000000000000000000000000000000000000000000000785.000000542.90000732.30000000005107107171010171973029715160232311909929order001247.00002039.21968.1001785.92026.4001881.502431.201282.501846.01849.81354.91543.402585.41956.91342.5001860.700002215.8001323.10000000000000000000000000000000000000000000000000000000000000000000000000000000000000000000000000000000000000000000051071071710101719630297151592323121111843491family001247.00002039.21968.1001785.92026.4001881.502431.201282.501846.01849.81566.01543.402585.41956.91342.5001871.500002215.8001323.10000000000000000000000000000000000000000000000000000000000000000000000000000000000000000000000000000000000000000000059710717108161963029714159232227121321243970genus001247.00002254.21968.1001785.92026.4001881.502431.201583.601956.01849.81566.01543.402585.41956.91426.7001871.500002215.8001379.0000000000000000000000000000000000000000000000000000000000000000000000000000000000000000000000000000000000000000000005105226569823species0000003040.80000000002431.20001954.200002635.000002299.100000000000000000000000000000000000000000000000000000000000000000000000000000000000000000000000000000000000000000000000000005105226551052265546271no\_rank0000003040.80000000002431.20001954.200002635.000002299.100000000000000000000000000000000000000000000000000000000000000000000000000000000000000000000000000000000000000000000000000005478166101642977129419191418610747732812637378no\_rank001247.00001271.01968.1001852.60001822.40001629.802058.71810.81688.51584.402429.41956.91227.3001575.800002040.1001180.7000000000000000000000000000000000000000000000000000000000000000000000000000000000000000000000000000000000000000000005971659716712538species000000000000002918.40000001729.201810.00000001972.60000000000000000000000000000000000000000000000000000000000000000000000000000000000000000000000000000000000000000000000000000115237115237713030species000000000000001324.2000000002164.60000001959.70000000896.60000000000000000000000000000000000000000000000000000000000000000000000000000000000000000000000000000000000000000000046179531111461795311111884263species001126.200002032.20000000000000001320.80001485.3001315.400001639.6001335.50000000000000000000000000000000000000000000000000000000000000000000000000000000000000000000000000000000000000000000015254108213530667749824271522082615933862512253482162542529484423813642539444269122948895573657361843489order002233.93399.52551.702961.53017.12998.73091.02954.72986.42518.72990.33179.503720.003081.63276.72799.73240.73247.33066.44005.73444.62324.62463.73212.202887.51487.13130.1004195.704831.82572.201877.200000000000000785.000000542.90000732.30000000000000000000000000000000000000000000000000000000000000000000785.000000542.90000732.300000000015254108213530667749824271522082615933862512253482162542529484423813642539444269122948895573657361222624232131977family002233.93399.52551.702961.53017.12998.73091.02954.72986.42518.72990.33179.503720.003081.63276.72799.73240.73247.33066.44005.73444.62324.62463.73212.202887.51487.13130.1004195.704831.82572.201877.200000000000000785.000000542.90000732.30000000000000000000000000000000000000000000000000000000000000000000785.000000542.90000732.30000000007636363676906genus0000000000000771.60000000626.000000000715.200000000000000000000000000000000000732.3000000000000000000000000000000000000000000000000000000000000000000000000000000732.3000000000999599907species000000000000000000000000000000499.100000000000000000000000000000000000567.5000000000000000000000000000000000000000000000000000000000000000000000000000000567.5000000000441064535no\_rank000000000000000000000000000000653.000000000000000000000000000000000000000000000000000000000000000000000000000000000000000000000000000000000000000000000000000008888881675036species000000000000000000000000000000242.800000000000000000000000000000000000242.8000000000000000000000000000000000000000000000000000000000000000000000000000000242.80000000001919191919192144175species0000000000000000000000000000001016.5000000000000000000000000000000000001016.50000000000000000000000000000000000000000000000000000000000000000000000000000001016.50000000001525410821353066774982325151728241593366251219347616164248948332379363953414426912204888514115356242132341211112013215362111110411529465genus002233.93399.52551.702961.53017.12998.73091.02957.43021.42518.72984.33186.603720.003088.73276.72804.43245.23246.23069.04005.73451.32326.32464.23212.203110.11487.13130.1004219.304831.82574.901877.2000000000000000000000000000000000000000000000000000000000000000000000000000000000000000000000000000000000000000000864475768115971413625151088863576112396099121648846579719242313217882049703675699113857412825158363404052220586737147583831971515231201786993729466species002387.53531.72625.503045.23247.72989.53154.12389.43021.42518.72981.23097.804049.303153.53030.93093.73063.93393.63230.84098.93491.61866.42723.4003173.51554.83326.6004479.5002162.60000000000000000000000000000000000000000000000000000000000000000000000000000000000000000000000000000000000000000000016869211482525231719192317517382640912121121686921148252523171919231751738264091212112479436no\_rank002121.13644.9003344.13820.93014.602710.6003170.32844.404320.002982.33033.83012.13259.33429.13126.903588.202498.20001409.40004361.7002269.800000000000000000000000000000000000000000000000000000000000000000000000000000000000000000000000000000000000000000000461317536213845255313306214655097923199279986761373094613175362138452553133062146550979231992799867613730939777species002128.50002970.13098.903116.71719.3003190.33287.80003281.23413.82897.83479.13188.23076.601902.32432.52111.0003033.61287.41127.3004007.4002682.4000000000000000000000000000000000000000000000000000000000000000000000000000000000000000000000000000000000000000000001963212161113762242659421976465931131467126248699423585243519196321216111376224265942197646593113146712624869942358524351939778species001882.43260.32430.602927.12775.9003018.9002726.23124.303520.503049.902698.72841.42933.52970.302539.21824.52274.43352.802884.01333.42860.1003631.5002579.7000000000000000000000000000000000000000000000000000000000000000000000000000000000000000000000000000000000000000000005666717212222661056667172122226610248315species000000890.4983.70000002002.2000881.701501.4826.51321.3983.802244.2332.7986.200000002749.100000000000000000000000000000000000000000000000000000000000000000000000000000000000000000000000000000000000000000000000411871117757573242739948genus00000000000003394.000002091.60004094.000470.200003021.900001150.90000000000000000000785.000000542.9000000000000000000000000000000000000000000000000000000000000000000000000785.000000542.9000000000000003811715381171539950species00000000000003593.000003010.00004094.000000003355.500000000000000000000000000000000000000000000000000000000000000000000000000000000000000000000000000000000000000000000000000007777772161821species0000000000000000000000000542.900000000000000000000000000000000000542.9000000000000000000000000000000000000000000000000000000000000000000000000000000542.9000000000000005555552582419species000000000000000000785.0000000000000000000000000000000000000785.0000000000000000000000000000000000000000000000000000000000000000000000000000000785.000000000000000000000610612267612410011372813514214146022411262626211737404class0003321.703440.702695.8003875.4003302.91671.72853.503038.03258.503293.12211.43212.03449.703432.101994.73049.502750.62179.04049.5004092.3000000000000000000000001816.90000000000000000000000000001816.9000000000000000000000000000000000000000000000000001816.9000000000000000061061226761221001127281351421414602241126262611737405order0003321.703440.702695.8003875.4003302.91671.72862.803038.03285.703293.12211.43212.03449.703432.101994.73049.502750.62179.04049.5004092.3000000000000000000000001816.90000000000000000000000000001816.9000000000000000000000000000000000000000000000000001816.900000000000000006106122676122100111728135142141460224112626261367738122310221570339family0003321.703440.702695.8003875.4003302.91671.72862.803038.03312.503293.12211.43212.03449.703432.101994.73049.502750.62179.04049.5004092.3000000000000000000000001816.90000000000000000000000000001816.9000000000000000000000000000000000000000000000000001816.90000000000000000699150022genus00000000000002180.203020.400000000000000000000000000000000000000000000000000000000000000000000000000000000000000000000000000000000000000000000000000000000000000000006991260species00000000000002180.203020.40000000000000000000000000000000000000000000000000000000000000000000000000000000000000000000000000000000000000000000000000000000000000000000699699334413no\_rank00000000000002180.203020.4000000000000000000000000000000000000000000000000000000000000000000000000000000000000000000000000000000000000000000000000000000000000000000016621162289genus0000000000000001807.90000002217.700000000000000000000000000000000000000000000000000000000000000000000000000000000000000000000000000000000000000000000000000000000000014514554005species0000000000000002014.80000002425.60000000000000000000000000000000000000000000000000000000000000000000000000000000000000000000000000000000000000000000000000000000000002626262626165779genus00000000000000000000001816.90000000000000000000000000000000000001816.90000000000000000000000000001816.9000000000000000000000000000000000000000000000000001816.90000000000000000151515151533034species00000000000000000000002295.90000000000000000000000000000000000002295.90000000000000000000000000000000000000000000000000000000000000000000000000000002295.900000000000000001515525919no\_rank00000000000000000000002295.90000000000000000000000000000000000000000000000000000000000000000000000000000000000000000000000000000000000000000000000000000000000001111111111111870984species00000000000000000000001163.80000000000000000000000000000000000001163.80000000000000000000000000000000000000000000000000000000000000000000000000000001163.80000000000000000610512258100104691021219384502049543311genus0003321.703440.703215.8003875.4003348.20003038.03502.603309.203640.63299.303627.502059.43049.503009.42264.74049.5004058.10000000000000000000000000000000000000000000000000000000000000000000000000000000000000000000000000000000000000000000000061051225810010469102121938450204961051225810010469102121938450204933033species0003321.703440.703215.8003875.4003348.20003038.03502.603309.203640.63299.303627.502059.43049.503009.42264.74049.5004058.100000000000000000000000000000000000000000000000000000000000000000000000000000000000000000000000000000000000000000000000747200795phylum00000000000000003899.000000000000003152.40000000000000000000000000000000000000000000000000000000000000000000000000000000000000000000000000000000000000000000000000000747292625class00000000000000003899.000000000000003152.40000000000000000000000000000000000000000000000000000000000000000000000000000000000000000000000000000000000000000000000000000747292629order00000000000000003899.000000000000003152.40000000000000000000000000000000000000000000000000000000000000000000000000000000000000000000000000000000000000000000000000000747292628family00000000000000003899.000000000000003152.400000000000000000000000000000000000000000000000000000000000000000000000000000000000000000000000000000000000000000000000000007471324991no\_rank00000000000000003899.000000000000003152.400000000000000000000000000000000000000000000000000000000000000000000000000000000000000000000000000000000000000000000000000007477471889813species00000000000000003899.000000000000003152.4000000000000000000000000000000000000000000000000000000000000000000000000000000000000000000000000000000000000000000000000000017641518132296827614073459113660279013635562710722961412173241198351244188219280419507283463255563102751522465552389610131486611115224655523891110132113111111201174phylum01663.22289.702023.002133.32884.83058.83215.72062.92991.62050.82721.33211.82592.02331.52710.32706.42422.52605.73278.62556.22575.13180.52813.51732.22650.5884.13196.22597.91323.12669.33352.204122.33526.03963.32231.31343.53165.600000399.20628.100002733.901278.20000486.2423.2001799.80386.72423.0000001501.900000517.80000787.300000386.700000000000657.4000000657.4000000000000000399.20628.100002733.901278.20000486.2423.2001799.80657.42423.0000001501.9001763917851952427514060459113276271713608623397229415110824384834123011120858041511717846316355277275052246555238910131486522465552389101352444353661453312862244184455211760class01663.22292.20002120.43026.02953.53226.02062.12991.62050.82812.83230.92592.13452.22710.32784.82422.52793.93305.22711.32693.83195.62834.91792.02687.9884.13196.22715.41315.12719.73352.204214.73526.04123.62213.91343.53178.500000399.20628.100002733.901278.20000486.2423.2001799.8002423.0000001501.900000517.80000787.30000000000000000000000000000000000000000399.20628.100002733.901278.20000486.2423.2001799.8002423.0000001501.90023622056572061394281944643412117843145127374893631628565236565232037order001550.60001719.51972.802518.71810.02643.201572.91725.71194.22452.82710.32093.901771.41829.91749.31254.102396.11389.81804.6002291.200002141.8001259.60000000399.2000000787.301278.20000486.200000000000000000000000787.30000000000000000000000000000000000000000399.2000000787.301278.20000486.2000000000000000236220565720613942819446434121178431451273748936316285652365652341421312133131412049family001550.60001719.51972.802518.71810.02643.201572.91725.71194.22452.82710.32093.901771.41829.91749.31254.102396.11389.81804.6002291.200002141.8001259.60000000399.2000000787.301278.20000486.200000000000000000000000787.30000000000000000000000000000000000000000399.2000000787.301278.20000486.2000000000000000152488520489511714518249562910712619141125151055235523315574102421988241941127101654genus001224.10001424.01978.201962.002654.801398.82504.61230.72452.83017.21966.901891.91813.41847.81213.402414.41096.02152.8002433.900001985.900884.50000000399.2000000001278.20000486.20000000000000000000000000000000000000000000000000000000000000000399.2000000001278.20000486.20000000000000004391110516485304391110516485301655species001695.200002155.60002094.70000002126.2003096.00975.802794.81555.62209.700000000000000000000000000000000000000000000000000000000000000000000000000000000000000000000000000000000000000000000000000000001132131812841132131812841656species000000000001390.40000001640.70001781.11058.101653.101245.5001688.8000000000000000000000000000000000000000000000000000000000000000000000000000000000000000000000000000000000000000000000000000055555552774species0000000399.200000000000000000000000000000000000000399.2000000000000000000000000000000000000000000000000000000000000000000000000000000399.200000000000000000000000000000555555111015species0000000000000000001278.20000000000000000000000000000000000001278.20000000000000000000000000000000000000000000000000000000000000000000000000000001278.2000000000000000000007181368283519871813682835198544580species000000000002388.00000001487.001679.51686.81931.2002734.202452.1001252.800002202.500000000000000000000000000000000000000000000000000000000000000000000000000000000000000000000000000000000000000000000000823287854116957227135259582328785411695722713525951852377species00684.20001401.12047.80001154.601373.82763.91242.503017.22870.002053.22643.400001438.02531.5002728.300000000000000000000000000000000000000000000000000000000000000000000000000000000000000000000000000000000000000000000000000001197342449195325232311112111015752609248no\_rank00000002204.90003094.20000001466.2001642.701431.402610.7893.62178.2002291.800000000000000000000000000000486.2000000000000000000000000000000000000000000000000000000000000000000000000000000486.200000000000000090167649739species000000000003145.8000000000003271.504041.10000000000000000000000000000000000000000000000000000000000000000000000000000000000000000000000000000000000000000000000000000000009016790167649743no\_rank000000000003145.8000000000003271.504041.1000000000000000000000000000000000000000000000000000000000000000000000000000000000000000000000000000000000000000000000000000000000622131125706438species000000000002274.80000001629.4002518.70001770.902149.50000000000000000000000000000000000000000000000000000000000000000000000000000000000000000000000000000000000000000000000000000000622131125622131125706439no\_rank000000000002274.80000001629.4002518.70001770.902149.500000000000000000000000000000000000000000000000000000000000000000000000000000000000000000000000000000000000000000000000000000007777772079536species00000000000000000000000584.6000000000000000000000000000000000000584.6000000000000000000000000000000000000000000000000000000000000000000000000000000584.60000000000000001616161616162560010species00000000000000000000000443.1000000000000000000000000000000000000443.1000000000000000000000000000000000000000000000000000000000000000000000000000000443.1000000000000000666661653174genus000000000000000787.30000000000000000000000000000000000000787.3000000000000000000000000000000787.300000000000000000000000000000000000000000000000787.3000000000000000000000066666659505species000000000000000787.30000000000000000000000000000000000000787.3000000000000000000000000000000000000000000000000000000000000000000000000000000787.300000000000000000000008381135349377142507111914351534723418114214312529408genus002162.80001906.12005.1001817.3001894.11620.3976.6002198.301705.11865.01545.31438.4001851.11666.6002238.300000001467.900000000000000000000000000000000000000000000000000000000000000000000000000000000000000000000000000000000000000000000838104524036913244671171434153342311783810452403691324467117143415334231171660species002162.80001906.12041.3001777.8001986.41639.5969.8002234.601710.91872.31545.31458.0001851.11658.5002257.300000001517.000000000000000000000000000000000000000000000000000000000000000000000000000000000000000000000000000000000000000000000998699986952773species00000001586.9000001484.0737.2000720.8000000002297.6000000000000000000000000000000000000000000000000000000000000000000000000000000000000000000000000000000000000000000000000000000097154724595525223467178452198148219885004order00000002149.802265.42750.200003346.30004034.402399.23804.43612.101784.300003353.700004550.8002973.9169.600000000543.9000000000000423.200000000000000000517.800000000000000000000000000000000000000000000000543.9000000000000423.2000000000000009715472459552522346717845219814821984262531331131953family00000002149.802265.42750.200003346.30004034.402399.23804.43612.101784.300003353.700004550.8002973.9169.600000000543.9000000000000423.200000000000000000517.800000000000000000000000000000000000000000000000543.9000000000000423.2000000000000001354222210046768447187185712273241678genus00000000003062.900003260.800000003874.802072.900003353.700005030.3002973.9217.500000000598.4000000000000423.200000000000000000000000000000000000000000000000000000000000000000598.4000000000000423.20000000000000085851680species0000000000166.500000000000000287.60000000000000000000000000000000000000000000000000000000000000000000000000000000000000000000000000000000000000000000000000000000001212121212121681species00000000001704.700000000000000000000000000000000000001704.70000000000000000000000000000000000000000000000000000000000000000000000000000001704.70000000000000000000000000009999991683species0000000000203.10000000000000000000000000000000000000203.1000000000000000000000000000000000000000000000000000000000000000000000000000000203.1000000000000000000000000000111196561961685species00000000002168.400000000000000000001788.900000002030.30000000000000000000000000000000000000000000000000000000000000000000000000000000000000000000000000000000000000000000066936351no\_rank00000000001796.5000000000000000000000000000000000000000000000000000000000000000000000000000000000000000000000000000000000000000000000000000000000000000000000000551385938no\_rank00000000004178.400000000000000000000000000000000000000000000000000000000000000000000000000000000000000000000000000000000000000000000000000000000000000000000000037371385939no\_rank00000000001476.1000000000000000000000000000000000000000000000000000000000000000000000000000000000000000000000000000000000000000000000000000000000000000000000000771385941no\_rank00000000004151.0000000000000000000000000000000000000000000000000000000000000000000000000000000000000000000000000000000000000000000000000000000000000000000000000383838438381686species0000000000390.60000000000000000000000000000000000000429.2000000000000000000000000000000000000000000000000000000000000000000000000000000429.2000000000000000000000000000345630129subspecies0000000000394.000000000000000000000000000000000000000000000000000000000000000000000000000000000000000000000000000000000000000000000000000000000000000000000000029291150460no\_rank0000000000440.60000000000000000000000000000000000000000000000000000000000000000000000000000000000000000000000000000000000000000000000000000000000000000000000001121205481661811689species0000000000723.000003412.100000003994.503499.800004024.700000000000000000000000000000000000000000000000000000000000000000000000000000000000000000000000000000000000000000000000000005145351453401473no\_rank00000000001099.60000000000003695.303550.30000000000000000000000000000000000000000000000000000000000000000000000000000000000000000000000000000000000000000000000000000000001561561150423no\_rank0000000000000003475.300000004692.7000000000000000000000000000000000000000000000000000000000000000000000000000000000000000000000000000000000000000000000000000000000001212127121228026species0000000000273.80000000000000000000000000000000000000324.4000000000000000000000000000000000000000000000000000000000000000000000000000000324.400000000000000000000000000055547043no\_rank0000000000395.200000000000000000000000000000000000000000000000000000000000000000000000000000000000000000000000000000000000000000000000000000000000000000000000088888833905species0000000000000000000000000423.200000000000000000000000000000000000423.2000000000000000000000000000000000000000000000000000000000000000000000000000000423.20000000000000094158787species0000000000790.400000000000000000001331.0000000000000000000000000000000000000000000000000000000000000000000000000000000000000000000000000000000000000000000000000000094941150461no\_rank0000000000790.400000000000000000001331.00000000000000000000000000000000000000000000000000000000000000000000000000000000000000000000000000000000000000000000000000000108763606762536301660216816species00000000003474.100000000000000177.000003319.900005030.3003038.80000000000000000000000000000000000000000000000000000000000000000000000000000000000000000000000000000000000000000000078037166971531679subspecies00000000003617.100000000000000000002905.100000003296.900000000000000000000000000000000000000000000000000000000000000000000000000000000000000000000000000000000000000000000115115565042no\_rank00000000003404.40000000000000000000000000003758.6000000000000000000000000000000000000000000000000000000000000000000000000000000000000000000000000000000000000000000003816838168759350no\_rank00000000003520.400000000000000000003183.200000003160.2000000000000000000000000000000000000000000000000000000000000000000000000000000000000000000000000000000000000000000005656890402no\_rank00000000002503.200000000000000000002989.2000000000000000000000000000000000000000000000000000000000000000000000000000000000000000000000000000000000000000000000000000016161035817no\_rank00000000004394.100000000000000000000000000000000000000000000000000000000000000000000000000000000000000000000000000000000000000000000000000000000000000000000000013131300227no\_rank00000000004020.2000000000000000000000000000000000000000000000000000000000000000000000000000000000000000000000000000000000000000000000000000000000000000000000000282228141682subspecies00000000002734.500000000000000000003119.4000000000000000000000000000000000000000000000000000000000000000000000000000000000000000000000000000000000000000000000000000088565040no\_rank0000000000000000000000000000004532.000000000000000000000000000000000000000000000000000000000000000000000000000000000000000000000000000000000000000000000000000002626205913no\_rank00000000002846.40000000000000000000000000000000000000000000000000000000000000000000000000000000000000000000000000000000000000000000000000000000000000000000000001481481481481482701genus0000000000522.00000000000000000000000000000000000000517.8000000000000000000000000000000517.800000000000000000000000000000000000000000000000517.80000000000000000000000000001481481481041481482702species0000000000522.00000000000000000000000000000000000000517.8000000000000000000000000000000000000000000000000000000000000000000000000000000517.80000000000000000000000000002323553190no\_rank0000000000389.700000000000000000000000000000000000000000000000000000000000000000000000000000000000000000000000000000000000000000000000000000000000000000000000021211009464no\_rank0000000000637.40000000000000000000000000000000000000000000000000000000000000000000000000000000000000000000000000000000000000000000000000000000000000000000000005345456196081genus0000000003063.6734.7000000000003869.400886.8000000000000000000000000000000000000000000000000000000000000000000000000000000000000000000000000000000000000000000000000000000000534545678259species0000000003063.6734.7000000000003869.400886.8000000000000000000000000000000000000000000000000000000000000000000000000000000000000000000000000000000000000000000000000000000000534545653454561150468no\_rank0000000003063.6734.7000000000003869.400886.80000000000000000000000000000000000000000000000000000000000000000000000000000000000000000000000000000000000000000000000000000000005566410196082genus00000003637.200490.400000000003002.20002169.20000000004615.400000000000000000000000000000000000000000000000000000000000000000000000000000000000000000000000000000000000000000000000556641078258species00000003637.200490.400000000003002.20002169.20000000004615.40000000000000000000000000000000000000000000000000000000000000000000000000000000000000000000000000000000000000000000000055664105566410864564no\_rank00000003637.200490.400000000003002.20002169.20000000004615.400000000000000000000000000000000000000000000000000000000000000000000000000000000000000000000000000000000000000000000000609170817272591241124411312522744631876722428590913765626874721558662357805983149446161345226285006order002315.90002130.03162.403294.41973.73325.52050.82872.63528.02904.0002963.02421.43217.93630.02594.72995.23495.43171.61972.62988.0002509.81486.82732.33201.504261.4001894.62852.53080.400000000000000000000000000000000000000000000000000000000000000000000000000000000000000000000000000000000000000000060917081682259124052441131242271459187172242839071376502687472131566035780598314944612121242142111268family002315.90002130.03156.003294.41974.63325.52050.82873.53532.22925.6002969.92421.43239.63636.82594.73018.63495.43171.61972.62877.6002515.51486.82732.33201.504261.4001894.62852.53080.4000000000000000000000000000000000000000000000000000000000000000000000000000000000000000000000000000000000000000000609170716822591240324411312422714511869722428390613564626872711311658347795983149461321211311535413132207genus002315.90002131.13156.003294.41974.83325.52050.82873.53532.22940.6002972.92421.43239.63640.82591.13036.43495.43174.71998.82876.6002521.61524.32732.63201.504261.4001894.603080.40000000000000000000000000000000000000000000000000000000000000000000000000000000000000000000000000000000000000000001643152569102574151389561945157236071713152130198715214335139543444514281371621721315830192047species002469.600003512.703318.41982.73482.22350.03441.83751.70003731.603501.93755.43563.03257.73509.13560.22235.52515.6003784.8003059.204282.2002693.100000000000000000000000000000000000000000000000000000000000000000000000000000000000000000000000000000000000000000000771631136515359745115172073901513771631136515359745115172073901513762948no\_rank002518.900003683.703425.91840.33440.92350.03471.83764.90003506.903492.23678.43649.72800.03514.93653.12197.20000003325.500000000000000000000000000000000000000000000000000000000000000000000000000000000000000000000000000000000000000000000000004451707135512392141630482115451144472232248868948426054127964330779450129464081316925835497114011383391365665419959481322187498674674616444883643675species002259.20002131.13083.7001975.13234.71801.52860.83516.12940.6002755.92421.63171.63642.32126.42990.502289.41924.32875.6002492.11368.32732.62253.004254.7001780.303080.40000000000000000000000000000000000000000000000000000000000000000000000000000000000000000000000000000000000000000003739143040384451647732601079569252928162735412176261636411037391430403844516477326010795692529281627354121762616364110680646no\_rank002397.20002349.73092.4001924.73346.61997.02796.63568.73195.6002706.62472.12852.53791.53145.63123.002396.02103.02866.8002724.71428.23077.7003835.7001869.603585.5000000000000000000000000000000000000000000000000000000000000000000000000000000000000000000000000000000000000000000115351933115351933172042species00000002120.00000003701.20003538.3000000003189.5000003376.80000000000000000000000000000000000000000000000000000000000000000000000000000000000000000000000000000000000000000000000000742078575genus0000000000000002334.0000000000000000000000002852.5000000000000000000000000000000000000000000000000000000000000000000000000000000000000000000000000000000000000000000074741618207species0000000000000002334.0000000000000000000000002852.5000000000000000000000000000000000000000000000000000000000000000000000000000000000000000000000000000000000000000000044243577468no\_rank00000003472.700000000000000000003585.70000000000000000000000000000000000000000000000000000000000000000000000000000000000000000000000000000000000000000000000000000000442432038genus00000003472.700000000000000000003585.70000000000000000000000000000000000000000000000000000000000000000000000000000000000000000000000000000000000000000000000000000000442433332039species00000003472.700000000000000000003585.700000000000000000000000000000000000000000000000000000000000000000000000000000000000000000000000000000000000000000000000000000002219622196203267no\_rank00000003543.000000000000000000003626.4000000000000000000000000000000000000000000000000000000000000000000000000000000000000000000000000000000000000000000000000000000019141914218496no\_rank00000003570.500000000000000000004231.90000000000000000000000000000000000000000000000000000000000000000000000000000000000000000000000000000000000000000000000000000000125772169673196971125761028421417649910136499101311313112185007order01240.92582.40002698.42426.9002283.8002340.01392.82754.5001802.804441.31392.22305.92843.40001485.6778.101925.61201.00004438.8002685.41365.100000000000002751.900000000001799.8002423.0000001501.900000000000000000000000000000000000000000000000000000000002751.900000000001799.8002423.0000001501.9001156718686701869711247682842141664991013649910131653family01317.22582.40003115.02426.9002561.2002340.01231.52764.8001865.604441.31392.22305.92843.40001541.8778.102452.01201.00004438.8002685.41438.600000000000002751.900000000001799.8002423.0000001501.900000000000000000000000000000000000000000000000000000000002751.900000000001799.8002423.0000001501.90011567186867018697112476828421416649910136499101321786811119741447818131716genus01317.22582.40003115.02426.9002561.2002340.01231.52764.8001865.604441.31392.22305.92843.40001541.8778.102452.01201.00004438.8002685.41438.600000000000002751.900000000001799.8002423.0000001501.900000000000000000000000000000000000000000000000000000000002751.900000000001799.8002423.0000001501.9009999991717species00000000000000000000000000001799.8000000000000000000000000000000000001799.80000000000000000000000000000000000000000000000000000000000000000000000000000001799.8000000000005510757104213551075710421343768species002582.40003331.40004096.600000004030.004961.2003762.40002855.700000004438.8002853.300000000000000000000000000000000000000000000000000000000000000000000000000000000000000000000000000000000000000000000989843770species0959.8000000000000000000000000001225.500000000000000000000000000000000000000000000000000000000000000000000000000000000000000000000000000000000000000000000000000000010101010101043990species00000000000000000000000000000002423.0000000000000000000000000000000000002423.00000000000000000000000000000000000000000000000000000000000000000000000000000002423.000000000207207146827species0000000000000003336.2000000000000688.6000000000000000000000000000000000000000000000000000000000000000000000000000000000000000000000000000000000000000000000000000000649649649649649161879species0000000000000002751.900000000000000000000000000000000000002751.90000000000000000000000000000000000000000000000000000000000000000000000000000002751.90000000000000000000000649649645127no\_rank0000000000000002751.900000000000000000000000000000000000000000000000000000000000000000000000000000000000000000000000000000000000000000000000000000000000000000001313131313169292species0000000000000000000000000000000000000001501.90000000000000000000000000000000001501.90000000000000000000000000000000000000000000000000000000000000000000000000000001501.9001313548476no\_rank0000000000000000000000000000000000000001501.90000000000000000000000000000000000000000000000000000000000000000000000000000000000000000000000000000000000000000000552624378no\_rank0000000000000000000000000000291.2000000000000000000000000000000000000000000000000000000000000000000000000000000000000000000000000000000000000000000000000000000419655911854555185009order0000003842.00004706.42193.30002972.4002156.400001606.20002527.7002911.100007046.400004307.000000004314.40000000000000000000000000000000000000000000000000000000000000000000000000000004314.400000000000000000000000000041965811854554158431957family0000003842.00004706.42193.30000002156.400001785.50002527.7002911.100007046.400004307.000000004314.40000000000000000000000000000000000000000000000000000000000000000000000000000004314.40000000000000000000000000001944551912216genus00000000004706.400000000000000003569.80000000000004307.000000004314.40000000000000000000000000000000000000000000000000000000000000000000000000000004314.400000000000000000000000000014446441747species00000000004846.400000000000000003569.80000000000004307.0000000000000000000000000000000000000000000000000000000000000000000000000000000000000000000000000000000000000000000881734925subspecies00000000004625.100000000000000000000000000000000000000000000000000000000000000000000000000000000000000000000000000000000000000000000000000000000000000000000000055555533011species00000000004314.400000000000000000000000000000000000004314.40000000000000000000000000000000000000000000000000000000000000000000000000000004314.400000000000000000000000000057451912217genus000000000002525.60000000000000001932.3003117.200007046.400000000000000000000000000000000000000000000000000000000000000000000000000000000000000000000000000000000000000000000000574557451750species000000000002525.60000000000000001932.3003117.200007046.400000000000000000000000000000000000000000000000000000000000000000000000000000000000000000000000000000000000000000000000528344412384734737119910981350147710643850923366111111214131221122284998class00002023.002954.72083.33164.003170.9001940.52500.802140.702212.102213.33008.42090.72287.70932.31646.01932.0002194.301879.5003958.7002377.5000000000000000000000000000386.700000000000000000000000386.700000000000657.4000000657.400000000000000000000000000000000000657.4000000000528342411380734636819810779349137710442750903121121284999order00002023.002954.72069.33164.002886.3001940.42500.802176.302209.002206.13001.82125.52284.00682.01646.01957.2002225.501879.5003993.1002377.100000000000000000000000000000000000000000000000000000000000000000000000000000000000000000000000000000000000000000000528342411378734636819710677349137610442550903152411331221121643824family00002023.002954.72069.33164.002886.3001950.22500.802176.302209.002216.23024.12160.02284.00682.01654.81957.2002234.201879.5003993.1002377.1000000000000000000000000000000000000000000000000000000000000000000000000000000000000000000000000000000000000000000005193374374723536019498623471175873885079291380genus00002023.002652.02081.63164.000001931.92512.501560.702225.702216.43094.52315.62292.60776.91671.31786.6002097.501879.5003990.7002356.2000000000000000000000000000000000000000000000000000000000000000000000000000000000000000000000000000000000000000000005193374374723536019498623471175873885079291382species00002023.002652.02081.63164.000001931.92512.501560.702225.702216.43094.52315.62292.60776.91671.31786.6002097.501879.5003990.7002356.200000000000000000000000000000000000000000000000000000000000000000000000000000000000000000000000000000000000000000000519337437472353601949862347117587388507929519337437472353601949862347117587388507929521095no\_rank00002023.002652.02081.63164.000001931.92512.501560.702225.702216.43094.52315.62292.60776.91671.31786.6002097.501879.5003990.7002356.200000000000000000000000000000000000000000000000000000000000000000000000000000000000000000000000000000000000000000000991058141737101514133925genus0000003593.80003215.2000004513.901971.6002161.51577.000002830.1003667.900004395.700000000000000000000000000000000000000000000000000000000000000000000000000000000000000000000000000000000000000000000000499326133926species0000003454.00000000004361.8000002343.600000003861.800003808.700000000000000000000000000000000000000000000000000000000000000000000000000000000000000000000000000000000000000000000000499326499326633147no\_rank0000003454.00000000004361.8000002343.600000003861.800003808.700000000000000000000000000000000000000000000000000000000000000000000000000000000000000000000000000000000000000000000000598516512638792no\_rank0000003705.60003215.200000000002161.5197.200002973.1002426.6000000000000000000000000000000000000000000000000000000000000000000000000000000000000000000000000000000000000000000000000000059841655984165712411species0000003705.60003215.200000000002161.5332.000002973.1002426.600000000000000000000000000000000000000000000000000000000000000000000000000000000000000000000000000000000000000000000000000001166111111111643822order000000000000000000000000000000982.200000000000000000000000000000000000386.700000000000000000000000386.700000000000657.4000000657.400000000000000000000000000000000000657.40000000001166111151151643826family000000000000000000000000000000982.200000000000000000000000000000000000386.700000000000000000000000386.700000000000657.4000000000000000000000000000000000000000000657.40000000006666684108genus000000000000000000000000000000386.700000000000000000000000000000000000386.700000000000000000000000386.7000000000000000000000000000000000000000000000000000000386.70000000006666684110species000000000000000000000000000000386.700000000000000000000000000000000000386.7000000000000000000000000000000000000000000000000000000000000000000000000000000386.700000000066471855no\_rank000000000000000000000000000000386.7000000000000000000000000000000000000000000000000000000000000000000000000000000000000000000000000000000000000000000000000000064446666544448phylum000000000000000000000000000000000112.3000000000000000000000000000000000000000000000000000000000159.000000000000159.0000000159.0000143.4000143.400000000000000000000000000000143.400000064446626231969class000000000000000000000000000000000112.3000000000000000000000000000000000000000000000000000000000159.000000000000159.0000000159.0000143.4000000000000000000000000000000000143.40000004444442085order000000000000000000000000000000000159.0000000000000000000000000000000000000000000000000000000000159.000000000000159.0000000159.00000000000000000000000000000000000000159.0000000444442092family000000000000000000000000000000000159.0000000000000000000000000000000000000000000000000000000000159.000000000000159.000000000000000000000000000000000000000000000159.00000004444442093genus000000000000000000000000000000000159.0000000000000000000000000000000000000000000000000000000000159.000000000000000000000000000000000000000000000000000000000159.000000055555551798711no\_rank00000000000000000000000001170.8000000000000000000000000000000000001170.8000000000000000000000000001170.8000000000001170.80000001170.8000000001170.8000000000000000000000001170.800000000000000555555551117phylum00000000000000000000000001170.8000000000000000000000000000000000001170.8000000000000000000000000001170.8000000000001170.80000001170.8000000001170.8000000000000000000000001170.8000000000000005555551301283subclass00000000000000000000000001170.8000000000000000000000000000000000001170.8000000000000000000000000001170.8000000000001170.80000001170.8000000000000000000000000000000001170.80000000000000055555551150order00000000000000000000000001170.8000000000000000000000000000000000001170.8000000000000000000000000001170.8000000000001170.80000001170.8000000000000000000000000000000001170.8000000000000005555551892254family00000000000000000000000001170.8000000000000000000000000000000000001170.8000000000000000000000000001170.8000000000001170.80000000000000000000000000000000000000001170.800000000000000555551158genus00000000000000000000000001170.8000000000000000000000000000000000001170.8000000000000000000000000001170.80000000000000000000000000000000000000000000000000001170.80000000000000055555482564species00000000000000000000000001170.8000000000000000000000000000000000001170.80000000000000000000000000000000000000000000000000000000000000000000000000000001170.80000000000000055179408no\_rank00000000000000000000000001170.8000000000000000000000000000000000000000000000000000000000000000000000000000000000000000000000000000000000000000000000000000000000596924605172481212841254112157superkingdom0000000268.600337.800000144.300000000000330.9000238.6220.1236.800240.00265.2000000000000000000000000000000000000000000000000000131.800000000000117.5000000117.5000131.800089.000000000000000000000000000000117.500000041828568888821528890phylum0000000000000000000000000000211.5000253.0287.4000216.80255.3000000000000000000000000000000000000000000000000000131.800000000000131.8000000131.8000131.8000000000000000000000000000000000131.80000004162768888812290931no\_rank0000000000000000000000000000211.5000267.2292.700000255.3000000000000000000000000000000000000000000000000000131.800000000000131.8000000131.8000131.8000000000000000000000000000000000131.800000041519612183963class0000000000000000000000000000211.5000279.7360.500000255.300000000000000000000000000000000000000000000000000000000000000000000000000000000000000000000000000000000000000000004141761644055order0000000000000000000000000000211.5000285.0380.700000255.300000000000000000000000000000000000000000000000000000000000000000000000000000000000000000000000000000000000000000004141761963271family0000000000000000000000000000211.5000285.0380.700000255.3000000000000000000000000000000000000000000000000000000000000000000000000000000000000000000000000000000000000000000041417656688genus0000000000000000000000000000211.5000285.0380.700000255.300000000000000000000000000000000000000000000000000000000000000000000000000000000000000000000000000000000000000000004141764722642239no\_rank0000000000000000000000000000211.5000285.0380.700000255.300000000000000000000000000000000000000000000000000000000000000000000000000000000000000000000000000000000000000000007127121537265species00000000000000000000000000000000258.0346.0000000000000000000000000000000000000000000000000000000000000000000000000000000000000000000000000000000000000000000000000054542518119species000000000000000000000000000000000464.000000261.000000000000000000000000000000000000000000000000000000000000000000000000000000000000000000000000000000000000000000008888888224756class000000000000000000000000000000000131.8000000000000000000000000000000000000000000000000000000000131.800000000000131.8000000131.8000131.8000000000000000000000000000000000131.800000088888894695order000000000000000000000000000000000131.8000000000000000000000000000000000000000000000000000000000131.800000000000131.8000000131.80000000000000000000000000000000000000131.8000000888882206family000000000000000000000000000000000131.8000000000000000000000000000000000000000000000000000000000131.800000000000131.800000000000000000000000000000000000000000000131.80000008888882207genus000000000000000000000000000000000131.8000000000000000000000000000000000000000000000000000000000131.800000000000000000000000000000000000000000000000000000000131.80000005556324121844441783275no\_rank0000000000172.400000156.400000000000426.4000195.3161.2225.500249.70268.500000000000000000000000000000000000000000000000000000000000000089.000000089.0000000089.00000000000000000000000000000089.00000005556284121828889phylum0000000000172.400000156.400000000000426.4000195.3171.5225.500249.70268.5000000000000000000000000000000000000000000000000000000000000000000000000000000000000000000000000000000000000000000055562841218183924class0000000000172.400000156.400000000000426.4000195.3171.5225.500249.70268.50000000000000000000000000000000000000000000000000000000000000000000000000000000000000000000000000000000000000000000555628412182281order0000000000172.400000156.400000000000426.4000195.3171.5225.500249.70268.5000000000000000000000000000000000000000000000000000000000000000000000000000000000000000000000000000000000000000000055562841218118883family0000000000172.400000156.400000000000426.4000195.3171.5225.500249.70268.50000000000000000000000000000000000000000000000000000000000000000000000000000000000000000000000000000000000000000000555628412182284genus0000000000172.400000156.400000000000426.4000195.3171.5225.500249.70268.5000000000000000000000000000000000000000000000000000000000000000000000000000000000000000000000000000000000000000000055562841218555628412182285species0000000000172.400000156.400000000000426.4000195.3171.5225.500249.70268.50000000000000000000000000000000000000000000000000000000000000000000000000000000000000000000000000000000000000000000444444651137phylum00000000000000000000000000000000089.000000000000000000000000000000000000000000000000000000000000000000000089.000000089.0000000089.00000000000000000000000000000089.00000004444431932order00000000000000000000000000000000089.000000000000000000000000000000000000000000000000000000000000000000000089.000000089.0000000000000000000000000000000000000089.0000000444444338190family00000000000000000000000000000000089.000000000000000000000000000000000000000000000000000000000000000000000089.00000000000000000000000000000000000000000000089.00000001622113291791581727196453210679149986421986572864875916299642198751110239superkingdom002638.100001239.602396.2506.8002794.91788.21982.81745.601848.802166.41714.41074.30002931.7985.6000003364.502033.805914.42251.10002597.400000956.70000573.001608.70000003048.6990.5000003906.106101.03050.0003010.800000000573.01608.70002469.203906.106074.33050.00000571.10000000000000000000002626.800000956.70000571.101608.70000003048.6990.5000003906.106074.33050.0000162211329179157172719645329679149986421986572864875916299642198752142131428883order002638.100001239.602396.2506.8002794.91788.21982.81283.301848.802166.41714.41074.30002931.7985.6000003724.302033.805914.42251.10002597.400000956.70000573.001608.70000003048.6990.5000003906.106101.03050.0003010.800000000573.01608.70002469.203906.106074.33050.00000571.10000000000000000000002626.800000956.70000571.101608.70000003048.6990.5000003906.106074.33050.0000616555510662family0000000000000000001995.502539.1000000000000000003050.00000000000000000000000000000000003050.00000000000000000000003050.0000000000000000000000000000000000000000000000000000000003050.0000616196896no\_rank0000000000000000001995.502539.10000000000000000000000000000000000000000000000000000000000000000000000000000000000000000000000000000000000000000000000000000000000000061661612402species0000000000000000001995.502539.1000000000000000000000000000000000000000000000000000000000000000000000000000000000000000000000000000000000000000000000000000000000000005555857479subfamily000000000000000000000000000000000000003050.00000000000000000000000000000000003050.00000000000000000000003050.0000000000000000000000000000000000000000000000000000000003050.0000555551196844genus000000000000000000000000000000000000003050.00000000000000000000000000000000003050.00000000000000000000003050.0000000000000000000000000000000000000000000000000000000003050.000055555510690species000000000000000000000000000000000000003050.00000000000000000000000000000000003050.00000000000000000000000000000000000000000000000000000000000000000000000000000003050.0000721912291756511619545259575996421586726877296421587456111231510699family003036.200001218.302377.8506.8002794.91599.64123.3766.601768.701619.01714.4993.20002931.7727.1000003724.302309.405914.400003021.200000956.70000001608.70000003048.6596.2000003906.106101.00003010.80000000001608.7000003906.106074.30000000000000000000000000003010.800000956.70000001608.70000003048.6596.2000003906.106074.3000019122945518522915915423455757196894no\_rank00000001218.302377.8506.800000609.001960.801896.81804.9993.20000726.300000000000000000000956.700000000000000596.2000000000000000000000000000000000000000000000000000000000000000956.700000000000000596.200000000000055555559241species000000000000000000000000000429.600000000000000000000000000000000000429.6000000000000000000000000000000000000000000000000000000000000000000000000000000429.6000000000000555555157924species000000000000000000000000000872.000000000000000000000000000000000000872.0000000000000000000000000000000000000000000000000000000000000000000000000000000872.0000000000000717717537874species00000001510.700313.60000000000000000000000000000000000000000000000000000000000000000000000000000000000000000000000000000000000000000000000000000000000000000000000008101181011644007species00000001184.202376.8000000000002459.000000000000000000000000000000000000000000000000000000000000000000000000000000000000000000000000000000000000000000000000000000000000009999991566990species0000000000956.70000000000000000000000000000000000000956.7000000000000000000000000000000000000000000000000000000000000000000000000000000956.70000000000000000000000000005555551701837species000000000000000000000000000487.000000000000000000000000000000000000487.0000000000000000000000000000000000000000000000000000000000000000000000000000000487.000000000000076771711623273genus00000000000000000000000000000000000005914.40000000000000000000000000000000006101.00000000000000000000006074.3000000000000000000000000000000000000000000000000000000006074.300006662029063no\_rank00000000000000000000000000000000000006101.00000000000000000000000000000000006101.00000000000000000000000000000000000000000000000000000000000000000000000000000006101.000006666661168612species00000000000000000000000000000000000006101.00000000000000000000000000000000006101.00000000000000000000000000000000000000000000000000000000000000000000000000000006101.0000064516464864131623286genus002974.40000000000000000000000000000000000000002941.4000000000000000000000000000000002945.90000000000000000000000000000000000000000000002945.900000000000000000000000000000000056515152115965no\_rank002945.90000000000000000000000000000000000000002941.40000000000000000000000000000000000000000000000000000000000000000000000000000002941.40000000000000000000000000000000001515151515151223261species002687.40000000000000000000000000000000000000002687.40000000000000000000000000000000000000000000000000000000000000000000000000000002687.40000000000000000000000000000000003636363636361223262species003047.20000000000000000000000000000000000000003047.20000000000000000000000000000000000000000000000000000000000000000000000000000003047.200000000000000000000000000000000013434242131623298genus00000000000003376.00000000000003001.8000000000000000000000000000000000003048.60000000000000000000000000000000000000000000000000000000000000000000000000000003048.6000000000000043424211636203no\_rank000000000000000000000000003001.8000000000000000000000000000000000003048.60000000000000000000000000000000000000000000000000000000000000000000000000000003048.600000000000001414141414141195068species000000000000000000000000002894.7000000000000000000000000000000000002894.70000000000000000000000000000000000000000000000000000000000000000000000000000002894.700000000000002828282828281308897species000000000000000000000000003125.5000000000000000000000000000000000003125.50000000000000000000000000000000000000000000000000000000000000000000000000000003125.50000000000000888881623299genus0000000000000000000000000000000003906.1000000000000000000000000000000000003906.10000000000000000000003906.1000000000000000000000000000000000000000000000000000000003906.1000000888881633149species0000000000000000000000000000000003906.1000000000000000000000000000000000003906.10000000000000000000000000000000000000000000000000000000000000000000000000000003906.100000088673832no\_rank0000000000000000000000000000000003906.10000000000000000000000000000000000000000000000000000000000000000000000000000000000000000000000000000000000000000000000000666661623304genus0000000000000000001608.70000000000000000000000000000000000001608.7000000000000000000000000000001608.70000000000000000000000000000000000000000000000001608.7000000000000000000006662231643no\_rank0000000000000000001608.70000000000000000000000000000000000001608.70000000000000000000000000000000000000000000000000000000000000000000000000000001608.7000000000000000000006666661051631species0000000000000000001608.70000000000000000000000000000000000001608.70000000000000000000000000000000000000000000000000000000000000000000000000000001608.700000000000000000000888882560098genus003530.20000000000000000000000000000000000000003530.2000000000000000000000000000000003530.20000000000000000000000000000000000000000000003530.2000000000000000000000000000000000888882560663species003530.20000000000000000000000000000000000000003530.20000000000000000000000000000000000000000000000000000000000000000000000000000003530.2000000000000000000000000000000000881225793no\_rank003530.2000000000000000000000000000000000000000000000000000000000000000000000000000000000000000000000000000000000000000000000000000000000000000000000000000000009054904490410744family002319.6000000000000000001630.40000002469.2000000000000002319.6000000000000000000002469.200000000000000000000000002469.200000000000000000000000000000002319.6000000000000000000002469.2000000000000909090196895no\_rank002319.60000000000000000000000000000000000000002319.60000000000000000000000000000000000000000000000000000000000000000000000000000002319.60000000000000000000000000000000009090909090901449437species002319.60000000000000000000000000000000000000002319.60000000000000000000000000000000000000000000000000000000000000000000000000000002319.6000000000000000000000000000000000544445542836subfamily000000000000000000001630.40000002469.2000000000000000000000000000000000002469.200000000000000000000000002469.200000000000000000000000000000000000000000000000000002469.2000000000000444441982583genus0000000000000000000000000002469.2000000000000000000000000000000000002469.200000000000000000000000002469.200000000000000000000000000000000000000000000000000002469.2000000000000444441982584species0000000000000000000000000002469.2000000000000000000000000000000000002469.20000000000000000000000000000000000000000000000000000000000000000000000000000002469.20000000000004410747no\_rank0000000000000000000000000002469.2000000000000000000000000000000000000000000000000000000000000000000000000000000000000000000000000000000000000000000000000000000098899912560065family000000000000000555.80000000000000000000000000000000000000573.0000000000000000000000000000000573.000000000000000571.100000000000000000000000000000000571.10000000000000000000000988812560081subfamily000000000000000555.80000000000000000000000000000000000000573.0000000000000000000000000000000573.000000000000000000000000000000000000000000000000573.00000000000000000000000888881980928genus000000000000000573.00000000000000000000000000000000000000573.0000000000000000000000000000000573.000000000000000000000000000000000000000000000000573.00000000000000000000000888881980930species000000000000000573.00000000000000000000000000000000000000573.0000000000000000000000000000000000000000000000000000000000000000000000000000000573.00000000000000000000000881340769no\_rank000000000000000573.00000000000000000000000000000000000000000000000000000000000000000000000000000000000000000000000000000000000000000000000000000000000000000000212411264828384no\_rank0000000000000000000000000002582.30000611.72403.50003295.0002500.6000000000000000000000000000000000000000000000000000000000000000000000000000000000000000000000000000000000000000000212411264881077no\_rank0000000000000000000000000002582.30000611.72403.50003295.0002500.60000000000000000000000000000000000000000000000000000000000000000000000000000000000000000000000000000000000000000002124112648212411264832630species0000000000000000000000000002582.30000611.72403.50003295.0002500.6000000000000000000000000000000000000000000000000000000000000000000000000000000000000000000000000000000000000000000
