## Supplementary Material for "BugSeq: a highly accurate cloud platform for long-read metagenomic analyses"

**Supplementary Information**

**Methods**

Precision (the positive predictive value) was defined by the number of correctly called reads divided by the total number of classified reads at the specified rank. Recall was defined as the number of correctly called reads divided by the total number of reads (unclassified and classified). F1 score was calculated as 2 × (Precision × Recall) / (Precision + Recall). Processing time and memory were calculated using the linux time utility, with the “-v” option.

The RefSeq reference database included all complete bacterial genomes; and all fungal, viral, protozoal and archaeal genomes, regardless of completion status. Additionally, the human genome and a database of contaminants (Univec) were included. The database was generated on February 23, 2020. For use with an alternative reference database, please get in touch with the corresponding author.

|  | ZymoBIOMICS Microbial Community Standards Dataset | Level | Mean F-Score | F-Score | Mean Precision | Precision, (%) | Mean Recall | Recall |
| --- | --- | --- | --- | --- | --- | --- | --- | --- |
| Bugseq | Even | Kingdom | 0.91 | 0.91 | 99.74 | 99.86 | 83.97 | 84.22 |
|  |  | Phylum |  | 0.91 |  | 99.86 |  | 84.14 |
|  |  | Order |  | 0.91 |  | 99.78 |  | 84.02 |
|  |  | Family |  | 0.91 |  | 99.71 |  | 83.93 |
|  |  | Genus |  | 0.91 |  | 99.65 |  | 83.81 |
|  |  | Species |  | 0.91 |  | 99.57 |  | 83.69 |
|  | Log | Kingdom | 0.95 | 0.95 | 99.95 | 99.97 | 90.49 | 90.54 |
|  |  | Phylum |  | 0.95 |  | 99.97 |  | 90.50 |
|  |  | Order |  | 0.95 |  | 99.94 |  | 90.48 |
|  |  | Family |  | 0.95 |  | 99.94 |  | 90.47 |
|  |  | Genus |  | 0.95 |  | 99.94 |  | 90.47 |
|  |  | Species |  | 0.95 |  | 99.92 |  | 90.47 |
| Centrifuge –min-hit 16 | Even | Kingdom | 0.92 | 0.98 | 96.72 | 99.95 | 88.05 | 96.12 |
|  |  | Phylum |  | 0.96 |  | 98.82 |  | 93.77 |
|  |  | Order |  | 0.94 |  | 96.93 |  | 90.75 |
|  |  | Family |  | 0.93 |  | 96.35 |  | 89.99 |
|  |  | Genus |  | 0.92 |  | 95.26 |  | 88.53 |
|  |  | Species |  | 0.79 |  | 93.00 |  | 69.17 |
|  | Log | Kingdom | 0.95 | 0.98 | 97.51 | 99.94 | 92.35 | 96.01 |
|  |  | Phylum |  | 0.97 |  | 98.82 |  | 94.38 |
|  |  | Order |  | 0.94 |  | 97.13 |  | 91.92 |
|  |  | Family |  | 0.94 |  | 96.71 |  | 91.46 |
|  |  | Genus |  | 0.94 |  | 96.51 |  | 91.24 |
|  |  | Species |  | 0.92 |  | 95.93 |  | 89.07 |
| Centrifuge –min-hit 22 | Even | Kingdom | 0.92 | 0.96 | 97.72 | 99.97 | 86.36 | 91.90 |
|  |  | Phylum |  | 0.95 |  | 99.25 |  | 91.06 |
|  |  | Order |  | 0.94 |  | 98.13 |  | 89.69 |
|  |  | Family |  | 0.93 |  | 97.65 |  | 89.17 |
|  |  | Genus |  | 0.92 |  | 96.63 |  | 87.80 |
|  |  | Species |  | 0.80 |  | 94.71 |  | 68.53 |
|  | Log | Kingdom | 0.95 | 0.97 | 98.33 | 99.95 | 91.51 | 93.99 |
|  |  | Phylum |  | 0.96 |  | 99.21 |  | 93.12 |
|  |  | Order |  | 0.95 |  | 98.14 |  | 91.37 |
|  |  | Family |  | 0.94 |  | 97.85 |  | 91.04 |
|  |  | Genus |  | 0.94 |  | 97.71 |  | 90.90 |
|  |  | Species |  | 0.93 |  | 97.14 |  | 88.65 |
| MetaMaps (miniSeq+H) | Even | Kingdom | 0.89 | 0.90 | 99.58 | 99.89 | 81.04 | 81.33 |
|  |  | Phylum |  | 0.90 |  | 99.85 |  | 81.21 |
|  |  | Order |  | 0.89 |  | 99.56 |  | 81.05 |
|  |  | Family |  | 0.89 |  | 99.51 |  | 81.01 |
|  |  | Genus |  | 0.89 |  | 99.51 |  | 80.96 |
|  |  | Species |  | 0.89 |  | 99.13 |  | 80.71 |
|  | Log | Kingdom | 0.94 | 0.94 | 99.65 | 99.69 | 88.33 | 88.46 |
|  |  | Phylum |  | 0.94 |  | 99.92 |  | 88.39 |
|  |  | Order |  | 0.94 |  | 99.54 |  | 88.31 |
|  |  | Family |  | 0.94 |  | 99.51 |  | 88.29 |
|  |  | Genus |  | 0.94 |  | 99.80 |  | 88.29 |
|  |  | Species |  | 0.94 |  | 99.46 |  | 88.25 |
| Metamaps (Refseq) | Even | Kingdom | N/A – Out of RAM | | | | | |
|  |  | Phylum |  |  |  |  |  |  |
|  |  | Order |  |  |  |  |  |  |
|  |  | Family |  |  |  |  |  |  |
|  |  | Genus |  |  |  |  |  |  |
|  |  | Species |  |  |  |  |  |  |
|  | Log | Kingdom | 0.94 | 0.94 | 99.81 | 99.98 | 88.72 | 88.88 |
|  |  | Phylum |  | 0.94 |  | 99.84 |  | 88.74 |
|  |  | Order |  | 0.94 |  | 99.77 |  | 88.69 |
|  |  | Family |  | 0.94 |  | 99.76 |  | 88.68 |
|  |  | Genus |  | 0.94 |  | 99.75 |  | 88.67 |
|  |  | Species |  | 0.94 |  | 99.73 |  | 88.66 |

**Supplementary Table 1:** Full performance characteristics on ZymBIOMICS mock communities.

| Sample | Sample type | Organism cultured by microbiology | Organism identified by BugSeq | Organism identified from original metagenomic pipeline (WIMP) | qPCR |
| --- | --- | --- | --- | --- | --- |
| S1 | ETA | *E. coli* | *E. coli* | *E. coli* |  |
| S2 | Sputum | *K. pneumoniae* | *K. pneumoniae* | *K. pneumoniae* |  |
| S3 | Sputum | *P. aeruginosa* | *P. aeruginosa* | *P. aeruginosa* |  |
| S4 | Sputum | *S. marcescens* | *S. marcescens* | *S. marcescens* |  |
| S5 | Sputum | *K. oxytoca* | *K. pneumoniae/K. oxytoca* | *K. pneumoniae/K. oxytoca* | *K. pneumoniae* not detected. |
| S6 | Sputum | *S. aureus* | *S. aureus* | *S. aureus* |  |
| S7 | Sputum | *H. influenzae* | *H. influenzae/P. aeruginosa* | *H. influenzae/P. aeruginosa* | *P. aeruginosa* detected. |
| **S8** | **Sputum** | ***M. catarrhalis*** | ***M. catarrhalis*** | ***M. catarrhalis/S. pneumoniae*** | ***S. pneumoniae* not detected.** |
| S9 | Sputum | *P. aeruginosa/E. coli* | *E. coli* | *E. coli* | *P. aeruginosa* not detected. |
| S10 | Sputum | NSG | *H. influenzae/S. pneumoniae* | *H. influenzae/S. pneumoniae* | *H. influenzae* detected.  *S. pneumoniae* detected. |
| S11 | Sputum | NRF | *S. pneumoniae* | *S. pneumoniae* | *S. pneumoniae* detected. |
| **S12** | **Sputum** | **NRF** | ***M. catarrhalis*** | ***H. influenzae/M. catarrhalis*** | ***H. influenzae* not detected. *M. catarrhalis* detected*.*** |
| S13 | Sputum | *S. marcescens* | *S. marcescens* | *S. marcescens* |  |
| S14 | Sputum | *S. aureus* | *S. aureus/M. catarrhalis* | *S. aureus/M. catarrhalis* | *M. Catarrhalis* detected*.* |
| **S15** | **Sputum** | ***S. aureus*** | ***S. aureus*** | ***S. aureus/S. pneumoniae*** | ***S. pneumoniae* not detected.** |
| S16 | Sputum | *S. aureus* | *S. aureus* | *S. aureus* |  |
| S17 | Sputum | NRF | NRF | None |  |
| S18 | Sputum | *H. influenzae* | *H. influenzae* | *H. influenzae* |  |
| S19 | Sputum | NRF | NRF | None |  |
| S20 | Sputum | *H. influenzae* | *H. influenzae* | *H. influenzae* |  |
| **S21** | **Sputum** | **NRF** | ***H. influenzae*** | ***H. influenzae/S. pneumoniae*** | ***H. influenzae* detected.**  ***S. pneumoniae* not detected.** |
| S22 | Sputum | NRF | NRF | None |  |
| S23 | Sputum | *H. influenzae* | *H. influenzae* | *H. influenzae* |  |
| S24 | Sputum | *H. influenzae* | *H. influenzae* | *H. influenzae* |  |
| S25 | Sputum | *H. influenzae* | *H. influenzae* | *H. influenzae* |  |
| S26 | Sputum | *M. catarrhalis* | *M. catarrhalis* | *M. catarrhalis* |  |
| S27 | Sputum | *H. influenzae/S. aureus* | *H. influenzae/S. aureus/S. pyogenes* | *H. influenzae/S. aureus/S. pyogenes* | *S. pyogenes* detected. |
| S28 | Sputum | NRF | *S. pneumoniae* | *S. pneumoniae* | *S. pneumoniae* not detected. |
| S29 | Sputum | *P. aeruginosa* | *P. aeruginosa/S. aureus* | *P. aeruginosa/S. aureus* | *S. aureus* detected. |
| S30 | BAL | *P. aeruginosa* | *P. aeruginosa* | *P. aeruginosa* |  |
| S31 | Sputum | NRF | *H. influenzae* | *H. influenzae* | *H. influenzae* detected. |
| **S32** | **Sputum** | **NSG** | ***E. coli/S. flexneri*** | ***E. coli*** | ***E. coli* detected. *S. flexneri* not tested.** |
| S33 | Sputum | NRF | NRF | None |  |
| S34 | Sputum | NSG | None | None |  |
| S35 | Sputum | *E. coli* | *E. coli* | *E. coli* |  |
| S36 | Sputum | *H. influenzae* | *H. influenzae* | *H. influenzae* |  |
| S37 | Sputum | *P. aeruginosa* | *P. aeruginosa* | *P. aeruginosa* |  |
| S38 | Sputum | *S. aureus/P. aeruginosa* | *S. aureus/P. aeruginosa* | *S. aureus/P. aeruginosa* |  |
| S39 | Sputum | *H. influenzae* | *H. influenzae/M. catarrhalis* | *H. influenzae/M. catarrhalis* | *M. Catarrhalis* detected*.* |
| S40 | ETA | *S. aureus* | *S. aureus* | *S. aureus* |  |
| S41 | Sputum | *H. influenzae/S. aureus* | *H. influenzae/S. aureus* | *H. influenzae/S. aureus* |  |

**Supplementary Table 2:** Performance characteristics on lower respiratory tract specimens. A 1% or greater abundance of any clinically significant microbe (defined by UK Standards for Microbiology Investigations) in a sample was called as present, as performed in the original publication. Differences in microbial identification are bolded. NRF=Normal respiratory flora. NSG=No significant growth.
